# Discovery of a Novel Chimeric Transposase-Transposon System for Advanced Genome Engineering

**DOI:** 10.64898/2025.12.02.691755

**Authors:** Daniel Heinzelmann, Franziska Reuss, Nikolas Zeh, Robin Nilson, Emely Walker, Juergen Fieder, Benjamin Lindner, Benjamin Renner, Patrick Schulz, Simon Fischer, Moritz Schmidt

## Abstract

Transposases revolutionized the field of genetic engineering, yet the scarcity of functional systems remains a challenge. In response, we conducted a metagenomic screening and identified a novel transposase system from *Acyrthosiphon pisum*. Through systematic optimization, we enhanced nuclear localization, transposon composition, and created a hyperactive transposase variant to boost transposition efficiency. Intriguingly, the combined application of the newly discovered transposase with inverted terminal repeat sequences from a related pea aphid species, *Aphis craccivora*, further enhanced transposition activity, resulting in the first chimeric transposase system reported so far. We investigated the genomic integration events following transposition in mammalian cells, to understand the underlying mechanisms and optimize the efficiency of transgene integration. This optimized system can expedite the generation of recombinant protein producing CHO cell lines, even surpassing the hyperactive *piggyBac* system with regards to cell specific productivity. These findings introduce a significant addition to the field of semi-targeted transgene integration technologies, offering substantial potential for enhancing biologics manufacturing.

## Introduction

Transposons are mobile DNA sequences possessing the unique ability to change their location within a genome^1^. In general, transposons can be categorized into two classes: class I and class II transposons ^2^. Class I transposons (e.g. retrotransposons) mobilize through a ‘copy and paste’ mechanism and constitute approx. 45 % of the human genome, whilst class II transposons (e.g. DNA transposons), many of which use a ‘cut and paste’ mechanism, can change their genomic location and only account for 3 % of the human genome ^3^. Most of the attention on transposon system over the recent years was gained by class II transposons as members of this family have emerged as attractive tools for genome engineering applications. For example, the semi-targeted integration of transgenes into mammalian cell expression systems revolutionized the production of biotherapeutics ^4–8^. Class II transposable elements encode a transposase enzyme, which is flanked by inverted terminal repeat (ITR) sequences that are recognized by the transposase, to carry out the process of transposition into a genomic locus^9^. The repurposing of this mechanism has transformed numerous genetic engineering applications, including genetic manipulation of pluripotent stem cells ^10^, engineering of bacteria ^11^ and the generation of transgenic animals ^12^. Additionally, it has found significant applications in gene therapy and biopharmaceutical cell line development. For instance, cell line development leverages the ‘cut and paste’ mechanism for transgene integration into the genome of mammalian cells ^8^. Prior to this, stable genomic integration of a transgene for biopharmaceutical cell line development was achieved via transfecting linearized DNA material into the cell, where integration events are based on randomly occurring genomic double strand breaks in the host cell DNA ^13^. This process is inefficient, error prone and often results in an uncontrolled integration of DNA fragments or concatemers at various genomic loci leading to genetic rearrangements and in turn to genetic instability of the recombinant mammalian production cell lines ^14,15^. Transposases usually target transcriptionally active and/or accessible genomic loci and result in a defined integration of the transposon cargo within the ITR sequences (e.g. transgene) without fragmentation, concatemerization or genomic rearrangements ^16,17^. Thus, transposase-mediated transgene integration results in exquisitely stable recombinant cell lines with high expression levels and moderate transgene copy numbers ^18,19^.

Although most published genomic sequences harbor transposase/transposon-like sequences, only very few have since been demonstrated to be active in mammalian cell lines. In nature, mobile genetic elements may have contributed to evolution but can also be a potential source of genomic instability, potentially detrimental to cells or organisms. This instability has presumably led to the inactivation of most transposase/transposon elements over time. Despite extensive research and debates on molecular evolution at the genomic level ^20,21^, the prevalent discovery of non-functional transposase/transposon pairs emphasizes that the mere presence of sequences resembling transposase/transposon does not necessarily indicate their activity. This further complicates the task of identifying new active pairs of transposase and transposon.

One of the first discovered and functionally active transposases is the *piggyBac* transposase/transposon pair, derived from the cabbage looper moth *Trichoplusia ni* (*T. ni*). This pair has been demonstrated to be active in a variety of cell species ^22,23^. Another prominent example is the Tc1/mariner-type transposase *Sleeping Beauty* derived from salmon fish ^24^. The identification and potential use of these systems have sparked a rush, leading to the discovery of additional active transposase/transposon pairs in organisms such as *Xenopus tropicalis*, *Bombyx mori*, *Myotis lucifugus, and Nilaparvata lugens* ^25–28^. And although large-language model aided design has even expedited the discovery of additional transposase systems ^29^, only a few functional and mostly hyperactivated transposase/transposon pairs are currently available for commercial use ^30^. Despite the scarcity of the systems, wide applicability of these demonstrates the transformative potential for various applications across different cell types. In this regard, relevant implications comprise drastically reduced development timelines for recombinant therapeutic proteins ^5,6^, increased productivity ^31–33^, improved cell line stability and facilitated development of complex multi-specific antibody formats ^4^.

Due to the proven value for diverse biopharmaceutical applications in combination with the limited availability, expanding the number of functional transposase/transposon systems has become critical. In this context, the continuously increased availability of new genomic sequences from various species and organisms supports discovery efforts to identify novel functional genetic elements. To address this, we embarked to discover new functional transposase/transposon systems in this study.

We present the identification of a novel and yet unknown transposase system in the pea aphid species *Acyrthosiphon pisum* (AP). This discovery was enabled through metagenomic screening across diverse species, which led to the identification of approximately 400 putative transposase and ITR sequence combinations. Among these, two transposases found in aphid species were confirmed to be functional in mammalian cells, with the one transposase from *Acyrthosiphon pisum* even showing exceeding activity to the hyperactive *piggyBac* transposase for the generation of Chinese hamster ovary (CHO) manufacturing cell lines. Furthermore, we delved into the systematic optimization of this newly identified transposase system, revealing its potential for superior performance across a diverse set of biopharmaceutical applications and cell types. This optimization process finally led to the establishment of the first chimeric hyperactive transposase system reported so far, further enhancing its activity and broadening its applicability.

## Results

### Identification of putatively novel transposase-transposon pairs via a metagenomic screening across various species results in the discovery of a yet unknown and functional transposase-transposon pair in the genome of the pea aphid species Acyrthosiphon pisum

In our pursuit to identify novel and functional transposase systems, a metagenomic screen was initiated. The aim was to search for genomically encoded transposase genes in proximity to ITR-like sequences across a variety of species (Fig. 1a). The initial screen yielded ∼400 hits, which were subsequently analyzed and assessed for their potential functionality and supporting evidence thereof. Transposases that deviated strongly (>100 AA) from the size of the *piggyBac* transposase from *T. ni* (594 AA) were de-selected. In contrast, we preferably selected transposase sequences that harbored the catalytic DDD triad motif (Fig. 1c) and exhibited a dimeric structural prediction using AlphaFold ^34,35^, similar to published *piggyBac* structures ^9^. Sequence homology to *piggyBac* transposase from *T. ni* was not considered a factor, with the focus on maintaining high sequence and species diversity. Additionally, the proximal 1 – 2 kb genomic sequences in 5’ and 3’ of the putative transposase genes were scanned for TTAA motifs, aiming to discover ITR-like repetitive motifs in proximity. By stringent de-selection, we aimed to experimentally confirm transposition capability for the most promising transposase/transposon pairs.

**Figure 1:**
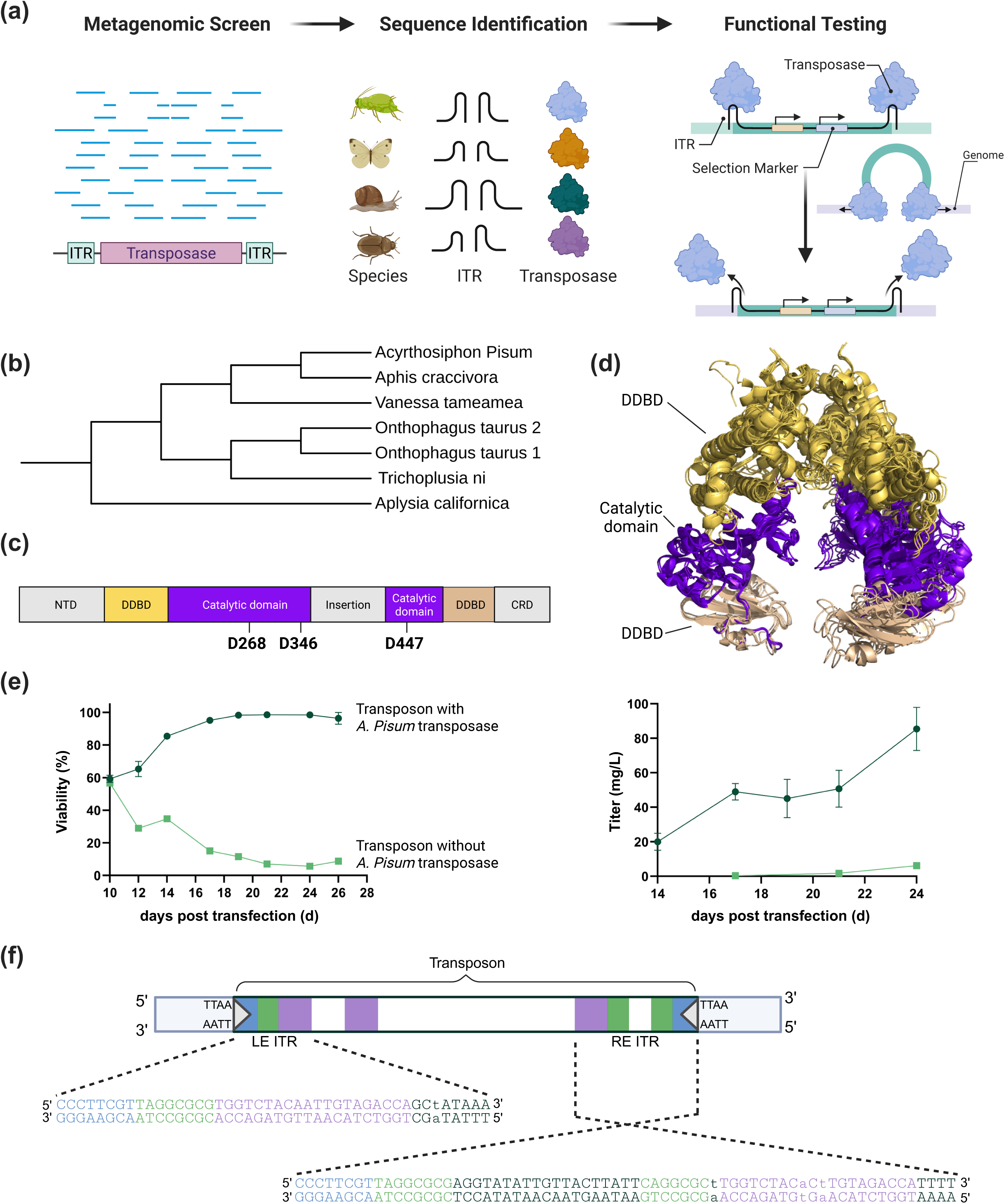
Discovery of novel, functional transposases in a diverse set of animal species. (a) Schematic representation of the metagenomic screening approach, including database mining, sequence identification, in-silico characterization, and functional testing in mammalian cells. (b) Phylogenetic tree of the top hits selected for experimental confirmation, including the originally discovered wildtype *piggyBac* transposase from *Trichoplusia ni* as a reference ^51^. (c) Schematic illustration of structural and functional domains, with the catalytic triad highlighted in bold (numbering according to the *piggyBac* transposase from *T. ni*). NTD = N-terminal domain; DDBD = Dimerization and DNA-binding domain; CRD = Cysteine-rich domain. (d) AlphaFold-modeled transposase dimers, overlaid for all hits (without NTD and CRD). (e) Functional test of the transposase identified in *Acyrthosiphon pisum*. Selection-marker based survival and recombinant protein expression was monitored following the co-transfection of a transposon with and without *Acyrthosiphon pisum* transposase. Error bars indicate standard deviation from two biological replicates. (f) Schematic representation of the left (LE) and right (RE) Inverted Terminal Repeats (ITR) of the Acyrthosiphon pisum (AP) and Aphis craccivora (AC) transposon sequences. Core transposase binding sites are highlighted in green, palindromic motifs predicted to be recognized by the CRD dimer are shown in purple, and the conserved terminal motif is marked in blue. Lowercase letters indicate sequence differences between the AC and AP variants.

Using this stringent selection process, six potential transposase sequences remained for functional testing in a mammalian cell line model (Fig. S1). Although species diversity was initially enforced, five of the six transposase/ITR combinations originated from insect species, with one additional hit found in gastropods (Fig. 1b). When examining the predicted structural dimeric scaffold in detail, all remaining hits adopt a dimeric state, comparable to *T. ni* (Fig. 1d). For comparison, the N-terminal domain (NTD) was excluded, as this domain adopts a rather unstructured and flexible conformation in its native state ^36^. The Cysteine-rich Domain (CRD) adopts a cross-brace zinc finger topology, consistent with the canonical fold observed in *piggyBac* transposases, and is likely involved in sequence-specific DNA binding ^37^. Overall, the identified hits exhibited a structural alignment to the reference cryo-EM structure of *T.ni* with root mean square deviation (RMSD) values of the models between 1.10 and 1.45 (Fig. 1d and Fig. S3), probably hinting on a shared biological function despite a rather low sequence homology (Fig. S2).

The identified *Onthophagus taurus 2* structure appears to lack a CRD domain (Fig. S3). Nevertheless, it was included for further experimental validation, as CRDs are known to mediate sequence-specific DNA binding at transposon ends ^37^, and their absence may reflect lineage-specific adaptations or alternative mechanisms of transposition. Additionally, monomeric structural comparison was conducted to elucidate whether certain domains deviate from the functional reference. In line with the overall high RMSD values for the dimeric scaffold, the monomers exhibited good alignment among each other, with only the structures from *Vanessa tameamea* and *Onthophagus taurus 2* showing some signs of misalignment, especially in the C-terminal area of the dimerization and DNA-binding domain (DDBD, Fig. S4).

To experimentally test the identified transposase systems for functional transposition in mammalian cells and to investigate their genome engineering capabilities, an auxotrophic glutamine synthetase (GS)-deficient CHO host cell line was transfected with a plasmid encoding the transposase under the control of a CMV promoter. Additionally, a transposon harboring the identified putative ITR sequences and genomic flanking regions was co-transfected with the transposase-encoding plasmid. For transposon construction, we selected approximately 300–400 nucleotides downstream of the genomic TTAA insertion motif for both the left end (LE) and right end (RE), aiming to encompass the core and palindromic binding motifs relevant for transposase activity. On the LE, a second palindromic motif was identified ∼200 nucleotides downstream of the initial TTAA site and was included due to its potential functional relevance.

Overall, the transposon encoded for an expression cassette to drive the expression of the heavy and light chain genes of an IgG1 antibody, as well as a GS selection marker gene to rescue the glutamine deficient cells during the metabolic selection. Cells were transferred to L-glutamine-free medium 24 hours post transfection, followed by monitoring of cell survival and recombinant IgG production over time (Fig. 1e). Despite stringent selection criteria and characterization, most identified top hits of the potential transposase/ITR combinations remained functionally inactive. Strikingly, the transposase hits from aphid species showed detectable transposition activity. Cells survived the selection process, leading to the establishment of stable IgG expressing pools. Hence, these findings rendered two highly homologous (98% protein sequence identity – Fig. S2) transposases identified from *Aphis craccivora* (AC) and *Acyrthosiphon pisum* (AP) functional for transposition in mammalian cells (Fig. 1e and Fig. S5), with their respective ITR sequences also exhibiting high similarity (Fig. S6 and Fig. S7).

To dissect the impact of differences between AP and AC derived ITR sequences, especially the ones within core transposase binding sites and palindromic motifs (Fig. 1f), and to identify a minimal ITR sequence different truncated variants of the ITR sequences were evaluated (Fig. S8). This analysis further suggested that, in addition to the core functional binding motifs and palindromic repeats, the flanking sequences of the LE and RE were essential for complete activity. Truncating regions outside these domains significantly reduced function, highlighting the broader contribution of surrounding sequence context beyond the more subtle differences observed in palindromic repeats.

### Systematic engineering of newly identified transposase and transposon sequences gives rise to a hyperactive transposase system for diverse biotechnological applications

Following the initial discovery of the functional transposase systems in the two aphid species, a systematic optimization process for both the transposase protein and the transposon was performed to enhance the genome engineering capabilities. Since the AP and AC transposases demonstrated a very high sequence identity, we focused on one of the enzymes and decided to further optimize the AP transposase. Optimization experiments were conducted as described above, by measuring the viability and recombinant IgG expression levels of stable pools based on GS-deficient CHO cells in course of selection.

First, we set out to optimize the nuclear import of the transposase enzyme after transfection. Although transposases typically encode natural nuclear localization signals (NLS), we investigated whether additional, artificial NLS sequences could further improve translocation of the transposase and thus affect transposition efficiency. We directly added a NLS either to the N-terminus or to both the N- and C-terminus of the AP transposase protein without any additional linkers and tested its efficiency in generating stable CHO cell pools (Fig. 2a). Adding a single NLS sequence at the N-terminus had only minor effects on cellular viability and the behavior in the selection process. In contrast, the attachment of NLS sequences to both N- and C-terminus resulted in accelerated cellular outgrowth during selection and significantly higher viabilities, suggesting improved transposition efficiency due to enhanced nuclear import. Stable recombinant CHO cell pools generated with the dual NLS harboring transposase reached a viable cell density (VCD) of 3.0×10^6^ viable cells per mL in shake flasks within 14 days post transfection. In comparison, constructs with only a single additional N-terminal NLS sequence or the wildtype NLS sequence without additional NLS achieved VCDs of 0.9×10^6^ and 1.5×10^6^ viable cells per mL, respectively. A similar trend was observed for cell viability during selection. Here, the dual NLS construct led to superior outgrowth during selection with cells reaching a viability of approx. 90% at day 9 post transfection, whereas the other variants remained below 90% viability even after 14 days into selection.

**Figure 2:**
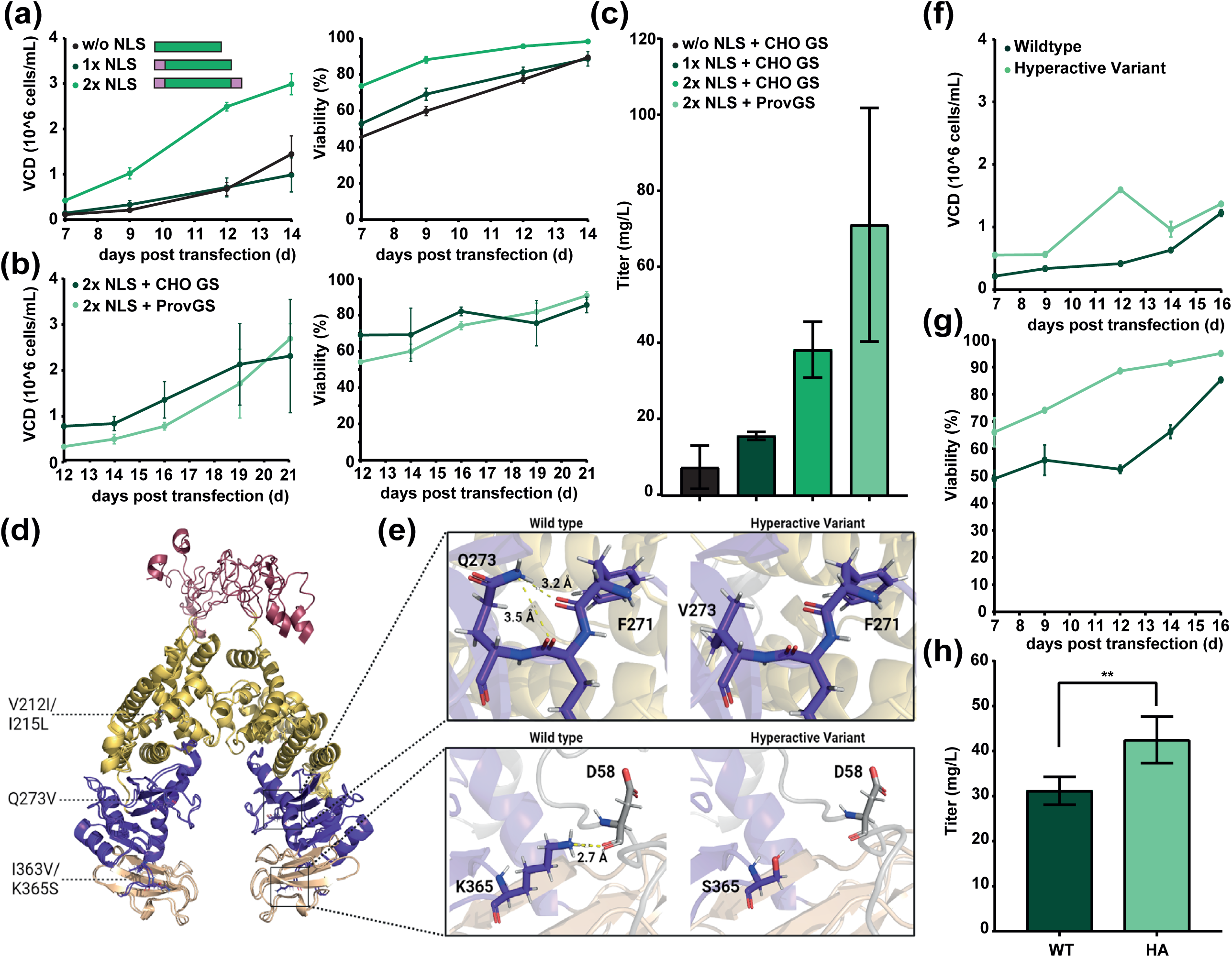
Systematic Optimization of the *Acyrthosiphon pisum* (AP) Transposase/Transposon System. (a) Enhancement in system efficiency by addition of N-terminal (1x NLS) or N- and C-terminal (2x NLS) nuclear localization signal (NLS) sequences to the transposase protein sequence. Data shows the cellular outgrowth behavior and cell viability of stable transfection pools in a GS selection system using different transposase-NLS constructs. NLS configurations are visualized in purple. (b) Comparison of the metabolic selection behavior after co-transfection of the dual NLS AP transposase variant (2xNLS) with transposons either harboring a CHO GS or ProvGS selection marker. (c) IgG titers in a 3-day batch cultivation of stable pools generated with various NLS-transposase constructs and GS transposon combinations. (d) AlphaFold-predicted dimer structure of AP wildtype (WT) transposase and its hyperactive variant. Mutated amino acid residues in the hyperactive variant (HA) are indicated. (e) Comparison of the wildtype AP transposase with its hyperactive variant at the mutated amino acid residues Q273V and K365S, showing impact of residue mutation on structural flexibility. (f) Outgrowth comparison of stable pools generated with wildtype AP transposase to its hyperactive variant during metabolic selection. (g) Cell viability comparison of stable pools generated with wildtype AP transposase to its hyperactive variant during metabolic selection. (h) Comparison of IgG titer of stable pools generated with wildtype AP transposase to its hyperactive variant at the end of a 3-day batch cultivation process. Error bars indicate standard deviation from two biological replicates.

To further enhance stable pool generation and recombinant IgG expression, we aimed to modulate selection stringency by replacing the endogenous GS selection marker derived from *Cricetulus griseus* (CHO GS) with a bacterially derived selection marker from *Providencia vermicola* (ProvGS) ^38,39^ on the transposon. Selection stringency plays a crucial role in enhancing cell productivity ^40,41^ and by integrating a bacterial enzyme, which may not exhibit optimal enzyme activity in eukaryotic cell culture environment, stringency may be increased. Cells transfected with the transposon harboring the ProvGS selection marker exhibited a comparable behavior during the selection process to those cells that were transfected with the CHO GS containing transposon (Fig. 2b). However, the productivity of cells generated with ProvGS was significantly increased (Fig. 2c) potentially indicating an increased selection stringency. The transposase variant without any additional NLS sequence and in combination with the transposon harboring the CHO GS achieved an IgG titer of 7.5 mg/L at the end of a 3-day batch culture, while the variant with one additional NLS sequence and the CHO GS transposon reached a titer of 16.0 mg/L. The dual NLS variant in combination with the CHO GS transposon showed a notable increase in IgG titer attaining 38.4 mg/L. The combination of the 2x NLS transposase variant with the ProvGS transposon eventually resulted in the highest IgG titer, reaching a mean value of 71.0 mg/L. In summary, the optimization of the transposase by addition of NLS sequences in combination with a transposon that harbors an attenuated GS selection marker led to an overall 9-fold increase in IgG titer without prolonging selection time, as compared to the initial transposase/transposon combination.

Finally, we aimed to engineer the AP transposase enzyme for functional hyperactivity to further increase transposition efficiency and recombinant protein expression in mammalian cells. We screened a range of mutations across the protein (Table S1) using structure-guided engineering, targeting enhanced DNA-binding, catalytic domain stabilization via disulfide bridges, and improved dimerization through increased C-terminal hydrophobicity. Following individual assessments of selection behavior and protein expression of the resulting cell pools, beneficial mutations were combined to optimize activity. The AlphaFold structural model of the transposase dimer was applied to rationalize the introduced mutations and investigate their structure-function relationship. After several rounds of screening and combining activity-enhancing mutations, we developed a hyperactive transposase variant (AP-HA) comprising five mutations: V212I, I215L, I363V, K365S, and Q273V (Fig. 2d). Structural predictions revealed that hyperactive variants may adopt a more flexible conformation while retaining a high degree of similarity to the wild-type dimeric scaffold. The Q273V mutation disrupts hydrogen bonding with the peptide backbone of F271 in the catalytic domain, potentially leading to increased conformational dynamics (Fig. 2e). Furthermore, the K365S mutation disrupts hydrogen bonding with D58 from the NTD, facilitating additional flexibility at the interface between the catalytic domain and the insertion domain (Fig. 2e).

The engineered AP-HA variant was benchmarked against the AP wildtype enzyme in the previously described experimental setup. The hyperactive variant demonstrated increased cellular outgrowth (Fig. 2f) and a faster recovery during metabolic selection (Fig. 2g), significantly outperforming the wildtype AP transposase. Again, IgG titers at the end of a batch cultivation process were compared. The hyperactive variant achieved an IgG titer of 42.5 mg/L thereby outperforming the wildtype AP transposase which reached 31.2 mg/L (Fig. 2h). In summary, these findings collectively underscore the effectiveness of incorporating additional NLS sequences, introducing an attenuated GS selection marker into the transposon, and integrating protein engineering strategies to enhance the overall performance and functionality of the AP transposase system.

### A cross species transposase-transposon exchange strategy gives rise to the first chimeric transposase system

Given that both the AP and AC transposase systems were functional in mammalian cells and exhibited high sequence homology – 98% at the protein level (Fig. S2) and between 87.5% and 95.6% for the left and right ITR and flanking sequences (Fig. S6 and Fig. S7) – we decided to further explore potential benefits of combining the two identified systems and hypothesized that a cross-species transposase-transposon exchange strategy could enhance the engineered system.

In such a chimeric approach, we tested the AP transposase variant in combination with the ITR sequences identified in AC and *vice versa*. The experimental design was driven by the hypothesis that such a combination could potentially harness remained functional integrity retained in each individual element, as it might be possible that the evolution of these homologous transposon pairs was not shaped by selective pressure for increased transposition activity and this lack of evolutionary constraint may have allowed the systems to diverge functionally. However, combining the AC transposase with the AP transposon resulted in cells not recovering from selection (Fig. S5), rendering this combination dysfunctional in the experimental setup. In contrast, combining the AP transposase with the AC transposon improved selection behavior when compared to the combination of the AP transposase and AP ITR sequences. Notably, the combination of the AC transposase and the AC ITR variants does not demonstrate significantly lower antibody titer values in comparison to the AC-ITR and AP-transposase combination. However, due to its improved outgrowth behavior during selection, the chimeric combination AP transposase with the AC transposon was chosen for further optimization.

Consequently, the experimental testing was expanded to incorporate all previously determined improvements, focusing on the evaluation of the AC- and AP-ITR sequences using both the wildtype and hyperactive AP transposase variants. As previously observed, the chimeric combination of the hyperactive AP transposase with the AC ITR sequences resulted in markedly improved recovery of stable cell pools post-transfection, achieving over 90% viability within less than 7 days of selection — outperforming the reference combination of the hyperactive AP transposase with AP ITR sequences (Fig. 3a). This performance significantly exceeded all other combinations, as none of the other tested variants achieved viabilities above 90% until 17 days post-transfection. Interestingly, while this improvement in performance was clearly measurable in the selection behavior, the absolute levels of recombinant protein expression remained comparable between the tested transposon/transposase pairs.

**Figure 3:**
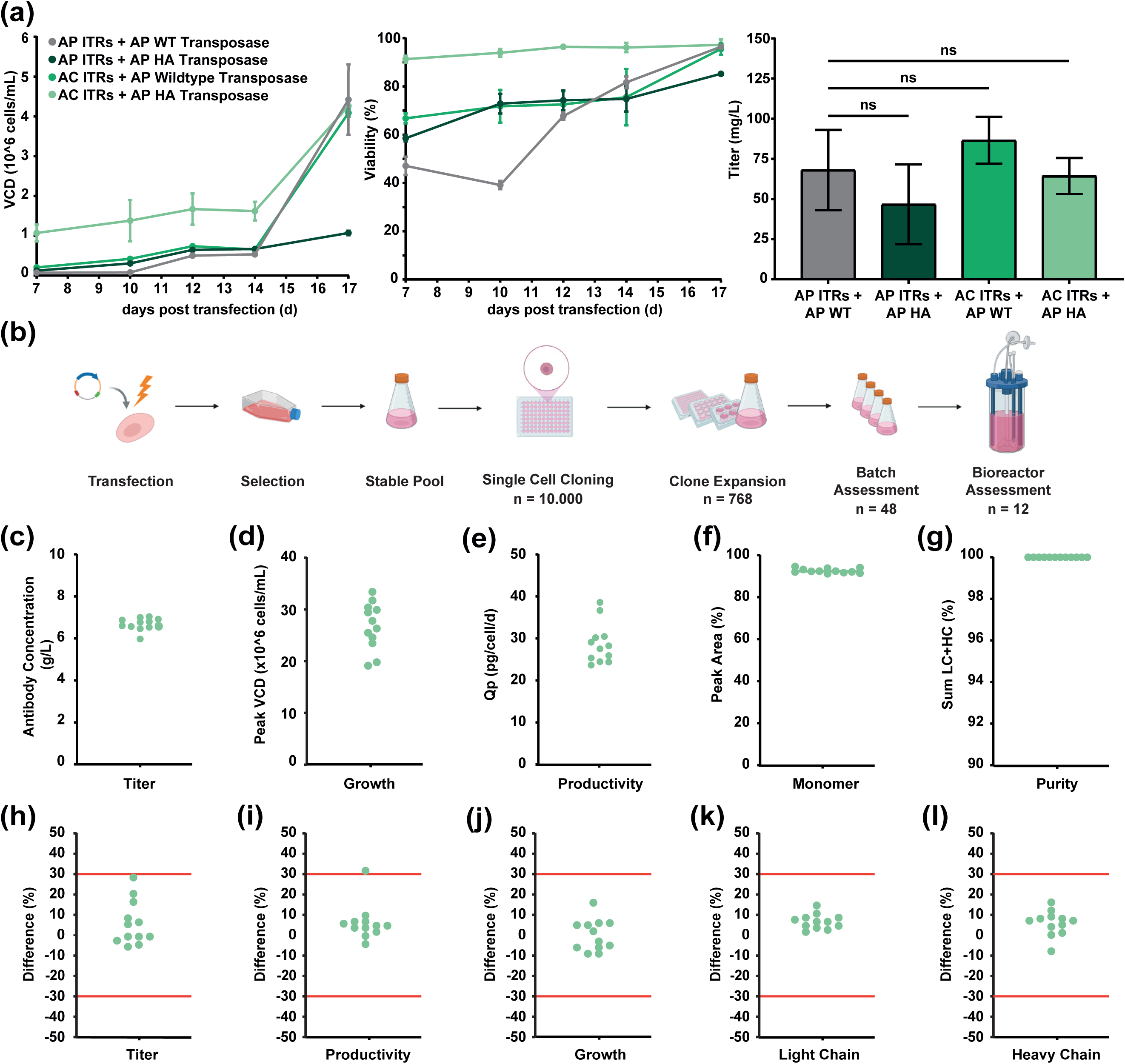
Optimization and characterization of a chimeric hyperactive transposase system shows improved cell culture performance in CHO cells with high titers, excellent product quality and long term stability. (a) Functionality of the cross-species combination of the wildtype (WT) and hyperactive (HA) *Acyrthosiphon pisum* (AP) transposase with the *Aphis craccivora* (AC) Inverted Terminal Repeat (ITR) sequences. Cellular outgrowth and viability of transfected pools during the metabolic selection process in L-glutamine-free medium was monitored and IgG production was determined in a 3 day batch cultivation process. Error bars indicate standard deviation from two biological replicates. (b) Schematic representation of the applied cell line development process to generate IgG expressing clonal manufacturing cell lines after transposase-mediated semi-targeted transgene integration in GS-deficient CHO host cells. Top 12 clones isolated out of the presented approach were assessed in a 14-day fed-batch bioprocess. (c) IgG product titer and (d) cell-specific productivity (Qp) at day 14. (d) Monomer content was determined by size exclusion chromatography, and (e) antibody purity (reported as a sum of LC and HC peaks) was determined by reduced capillary gel electrophoresis. All shown product quality parameters were analyzed following Protein A purification of harvested cell culture fluid. Phenotypic expression stability of the integrated vector copies was verified by comparing the cells at an age of 55 days in culture to cells that were harvested after 37 days in culture. Relative difference between the two time points for (h) antibody titer, (i) Qp, (j) cell growth, and (k) light chain and (l) heavy chain copy numbers were calculated. Criteria for stability (± 30 %) is indicated in red.

Due to its improved metabolic selection behavior and IgG productivity, the chimeric AP-HA/AC system — comprising the hyperactive AP transposase variant (V212I, I215L, I363V, K365S, Q273V) and AC ITR sequences — was selected for further evaluation.

To finally demonstrate its fit-for-purpose in relevant biopharmaceutical processes and for further in-depth clonal characterization in CHO cells, stable cell pools were subjected to single-cell deposition (Fig. 3b) to isolate clonal cell lines. Subsequently, cells were screened for growth behavior and IgG productivity to identify the best-performing clonal cell lines. Finally, after the expansion phase and batch assessment in shake flasks, 12 clones were selected for small-scale bioprocess assessment from the initial pool of over 10,000 deposited single CHO cells. The product titer produced by these clones ranged from 5.9 to 7.0 g/L over a 14-day process (Fig. 3c) with a consistent peak VCD between 19.1 to 33.4 ×10^6^ cells/mL (Fig. 3d). The cell-specific productivity (qP) was observed to be between 26.6 to 38.6 pg/cell/day (Fig. 3e). Focusing on antibody product quality parameters, the monomer content was consistent among all AP-HA/AC clones, maintaining a tight range around 91% (Fig. 3f). The purity of antibody produced by these twelve clones, as determined by reduced capillary gel electrophoresis and calculated based on the sum of the peak areas corresponding to the light and heavy chain, was found to be close to 100% (Fig. 3g).

Moreover, these 12 clones from the bioprocess assessment have undergone long term cell line stability assessment by continuous cultivation for up to 55 days in culture (dic). Phenotypic expression stability of the integrated vector copies was verified by comparing the cells at an age of 55 dic to cells that were harvested after 37 dic. Phenotypic parameters from a small-scale bioprocess run were compared between these two time points and only one of the analyzed clones showed a change in antibody titer (Fig. 3h), productivity (Fig. 3i), or growth (Fig. 3j) greater ± 30 %. The genetic stability was evaluated by transgene copy number determination via droplet digital Polymerase Chain Reaction (ddPCR), showing that none of the clones showed a change of copy numbers ± 30 % (Fig. 3k and Fig. 3l). Additionally, Southern blot analysis of the integrated light chain (Fig. S9) and heavy chain (Fig. S10) genes verified the genetic stabilities of the analyzed cell lines, as indicated by comparable restriction enzyme pattern between the cells after long term cultivation and the respective cryopreserved cell bank generated beforehand. Thus, the vector integrations generated with the AP-HA transposase in combination with the AC transposon showed exceptionally high genetic stability with an accompanying expression stability even after long term cultivation.

Overall, the chimeric combination of two related transposase systems reveals a design principle that significantly improves activity and genome engineering capabilities by leveraging the strengths of each evolved system, potentially circumventing detrimental mutations that may have accumulated over time in aphid species. Moreover, the presented results from the bioprocess assessment indicate that the chimeric AP-HA/AC system shows promising results in terms of both productivity and expression stability, suggesting its suitable applicability in commercial biopharmaceutical manufacturing systems.

### In-depth characterization of integration sites indicates integration preferences of the hyperactive AP/AC transposase system in CHO cell lines

In addition to the presented phenotypic characterization of clones derived from the chimeric AP-HA/AC system in a representative small-scale bioprocess, also a deepened genetic characterization was of interest to assess the genome-wide distribution of integration sites. To address this, genomic DNA (gDNA) was extracted from the 48 clones evaluated in a batch assessment (Fig. 3b) to study the precise genomic location of the integrated transposon vector via next generation sequencing (Fig. 4a). Specifically, hybridization-based capture was used to enrich the vector breakpoints, representing the junctions between the transposon ends and the host cell genome, from a sequencing library. Resulting reads covering the breakpoints were extracted and mapped to a reference genome. Additionally, ddPCR was used to determine transgene copy numbers, as potential offsets between integration sites and copy number could indicate random integration or concatemerization.

**Figure 4:**
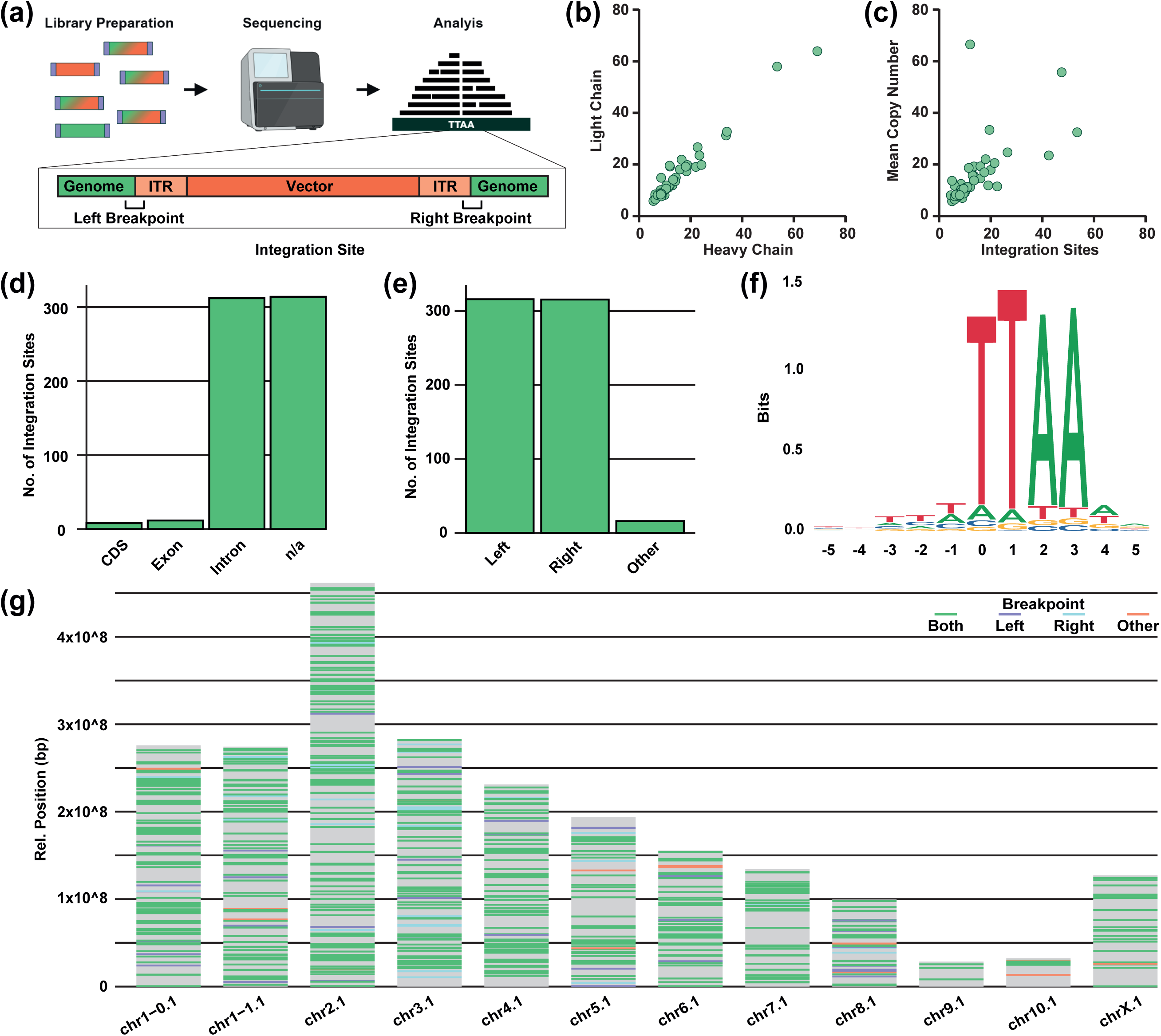
Integration site analysis of 48 clonal CHO cell lines reveals preferences of the chimeric *Acyrthosiphon pisum* transposase system on genomic context and motif for vector integration. (a) Process for integration site analysis performed in this study. Following library preparation and Next Generation Sequencing, reads are mapped to the genome and reads covering the left and right breakpoints between the genome and the Inverted Terminal Repeats (ITR) from the vector are extracted to identify vector integration sites. (b) Mean copy number calculated from both the Light Chain and Heavy Chain copies juxtaposed with the overall number of identified vector integration sites. (c) Analysis of the genomic context of the integrated expression vectors and (d) the breakpoints of the vectors based on the genomic annotation. (e) Sequence logo plot showing nucleotide conservation across integration motifs. Position 0 was defined relative to the vector breakpoint, and the plot visualizes sequence homology ±5 bases upstream and downstream of this site. The height of each nucleotide at a given position reflects its relative frequency and contribution to overall sequence conservation.. (f) Chromosome blot indicating positions within the host genome of all integration sites determined in 48 clones. Type of identified vector breakpoints for each integration site indicated by different colors.

First, the determined copy numbers for light and heavy chain were compared for each clone (Fig. 4b). Copy number valued ranged from 6 copies per cell up to 70 copies. However, indicated balance between the number of light and heavy chain genes (r = 0.98) suggest no detectable levels of fragmentation at the integration sites even for high copy number clones. Next, transgene copy number was compared with the overall number of identified vector integration sites (Fig. 4b). To account for the light and heavy chain numbers, and assuming that each locus contains only one copy each, the mean value between the two determinations was used for this analysis. A noteworthy correlation (r = 0.62) was observed between the integration sites and the transgene copy numbers, suggesting a one-copy-per-locus integration rather than concatemerization ^14^. For most of the clones, the number of transgene integration sites aligned with the LC and HC copy numbers. However, especially one clone showed a high offset with copy numbers exceeding the identified number of integration sites approximately 3 to 4-fold. The genomic context of expression vector integration sites and the breakpoints of the vectors were thoroughly analyzed based on the genomic annotation of the host cell genome (Fig. 4d and Fig. 4e). Out of the 648 determined integration sites, a distinct preference was observed for integration sites into genes, particularly into intronic regions (312 integration sites) in line with published data ^42^. Additionally, a number of 314 integration sites was found to be located in regions lacking annotations. Conversely, only a minority of 12 integration sites were found to be located in exons, and with a number of 8 even less within coding sequences (CDS) of genes. Upon examination of the vector breakpoints, it was found that 98 % of the integration sites matched both the expected left and right breakpoints, clearly indicating that the transposase has integrated its transposon exactly from the left to the right ITR. This was complemented by the observation that only 2 % of integration sites and a corresponding number of 12 breakpoints were identified in different regions of the vector than the left or right ITR sequences. The motif of the integration sites was subsequently analyzed (Fig. 4f) and showed a TTAA tetranucleotide as the most frequent motif (83 % - Fig. S11) at either the left or right breakpoint, proofing the intended controlled transposition event. Interestingly, the integration sites were found to be spread across the genome (Fig. 4g) and no distinct chromosomal region was complete void of transposase-mediated integration sites. In-depth analysis of the integration sites revealed that 16 genes were targeted by the transposase in multiple clones but having different positions. Furthermore, we identified two integration sites present in two distinct clones with identical genomic positions. Moreover, one clone had two integration sites in the same gene only kilobases apart. Taken together, genome-wide distribution analysis further underpins the preference of the transposase for a certain motif or genomic context rather than specific genes.

In conclusion, the newly discovered chimeric AP-HA/AC transposase system demonstrates high fidelity and precision in integrating transposon cargo into a wide variety of genomic loci. Additionally, a preferred integration into the tetranucleotide motif “TTAA” was observed, which is comparable to *piggyBac*-derived transposases ^43^, while being only distantly related to the *piggybac*-superfamily due to the low sequence homology.

### Benchmarking of the chimeric and hyperactive AP/AC system against piggyBac reveales superior performance of the novel system for biomanufacturing in CHO cells

To enable a robust evaluation of the final AP/AC system’s (Fig. 5a) performance and activity, a benchmarking study was conducted using the hyperactive piggyBac transposase (pB-HA) as a reference ^22^. This system was selected due to its well-characterized and commercially validated nature, as well as the close resemblance of the AP-HA transposase to the *piggyBac* transposase in terms of predicted structural features and integration site preferences. In course of the benchmarking, CHO cells were co-transfected with either the AP-HA or pB-HA transposase, along with an antibody-encoding transposon containing the corresponding ITR sequences. Following transfection and metabolic selection, cell growth and viability were monitored throughout the selection period (Fig. 5b and Fig.5c). By day 16 post transfection, both transposase systems demonstrated comparable performance in terms of cell proliferation and viability. Specifically, cell densities reached approximately 1.5 × 10^6^ cells/mL, and viabilities reached around 90% for both AP-HA and pB-HA pools. However, a significant difference was observed in antibody production. After a 3-day shake flask cultivation, the AP-HA pools yielded a mean antibody titer of 148.1 mg/L, compared to 98.3 mg/L for the pB-HA pools (Fig. 5d). Genomic copy number analysis revealed that the AP-HA pools harbored a higher mean transposon copy number per cell, calculated as the mean of light and heavy chain integrations. Specifically, AP-HA pools exhibited 13.7 mean copies per cell, whereas pB-HA pools showed 8.2 mean copies per cell (Fig. 5e). These findings suggest that the improved expression observed in AP-HA pools may be attributed to more efficient transposon integration, thereby enhancing transgene dosage and overall productivity.

**Figure 5:**
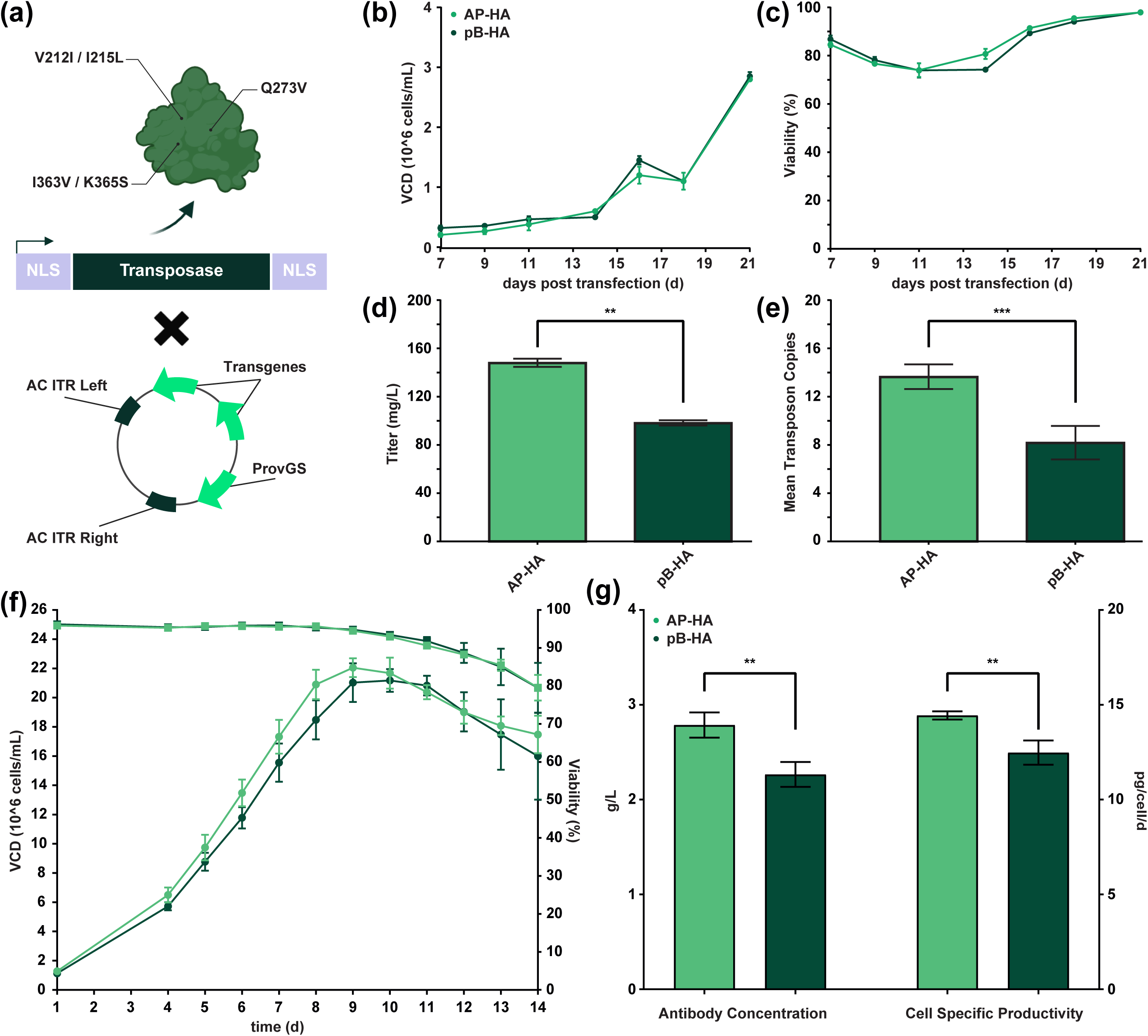
Benchmarking of the hyperactive *Acyrthosiphon pisum* (AP-HA) transposase system against the hyperactive *piggyBac* (pB-HA) demonstrates its superior performance for antibody expression in CHO cell lines. (a) Summary of all included interventions in the final AP-HA transposase system: AP-HA transposase with indicated mutations and flanking nuclear localization signals (NLS), transposon harboring Inverted Terminal Repeat (ITR) sequences from *Aphis craccivora* (AC), and a metabolic selection marker from *Providencia vermicola* (ProvGS). (b) Viable cell density (VCD) monitoring during metabolic selection, following the transfection of CHO cell pools with an antibody-encoding transposon using either the AP-HA or pB-HA transposase. (c) Percentage of viable cells in course of the selection process. (d) Antibody concentration was determined at the end of a shake flask cultivation for 3 days. (e) Mean transposon copy number per cell, calculated as an average between determined numbers of light and heavy chain copies. (f) Viable cell density (VCD) and cell viability measured in course of the 14-day fed-batch process. (g) Antibody concentration measured at day 14 (left) and average calculated cell specific productivity (Qp – right) across the fed-batch process. All error bars indicate standard deviation from two biological replicates.

To further investigate the performance of both systems under conditions more representative for biopharmaceutical production, the respective cell pools were subsequently cultivated in small-scale bioreactors. Throughout the 14-day fed-batch process, both cell pools exhibited comparable growth kinetics and viability profiles. Peak VCDs reached approximately 22×10^6^ cells/mL (Fig. 5f), and harvest viabilities were maintained at around 80 % (Fig 5g). At the end of the fermentation, the AP-HA-derived cell pools demonstrated a higher antibody titer compared to the pB-HA pools, consistent with observations from the shake flask experiments (Fig. 5g). This increase in volumetric productivity was also reflected in the calculated cell specific productivity values, which were found to be higher for the AP-HA pools relative to the pB-HA pools achieving (Fig. 5h).

Overall, the AP-HA transposase system demonstrated superior performance compared to the commercially established pB-HA system in the context of antibody production in CHO cell lines. Across both shake flask and bioreactor cultivations, higher antibody titers and cell specific productivities were consistently observed in AP-HA-derived cell pools. Taken together, the AP-HA system represents a promising alternative for efficient and scalable transgene integration in biopharmaceutical manufacturing.

### Transposition of increased cargos allows expression of large heterologous proteins using the AP transposase

To further demonstrate the versatility and broad applicability of the AP transposase system, its use was extended to the expression of large molecules in a heterologous system. This approach aimed to evaluate the system’s capability to transpose large vectors, thereby enabling the expression of significantly larger proteins.

First, we focused on evaluating the impact of increasing transposon cargo size on transposition efficiency. During the initial development and engineering of the system, the cargo load between the left and right ITR sequences did not exceed 9 kb in size. Therefore, we created expression vectors with larger cargo capacities. Cargo sizes of 13.9 kb and 18.8 kb were tested against a standard antibody transposon with a size of 8.6 kb. The increase in cargo size was achieved by doubling or tripling the antibody expression cassette, while keeping the same metabolic selection marker set-up (Fig. 6a). During the selection process, cell growth and viability in course of the selection process were closely monitored (Fig. 6b) as direct indicators of transposition efficiency. The reference construct demonstrated the most robust outgrowth, reaching approximately 3 million cells after 17 days, compared to about 2 million cells for the extended cargo samples. Although the reference construct displayed a slightly better viability profile, the titer at the end of the shake flask passage showed comparable levels of recombinant protein across all constructs. This indicated that the overall transposition efficiency of the system was maintained, even with cargo sizes up to 18.8 kb.

**Figure 6:**
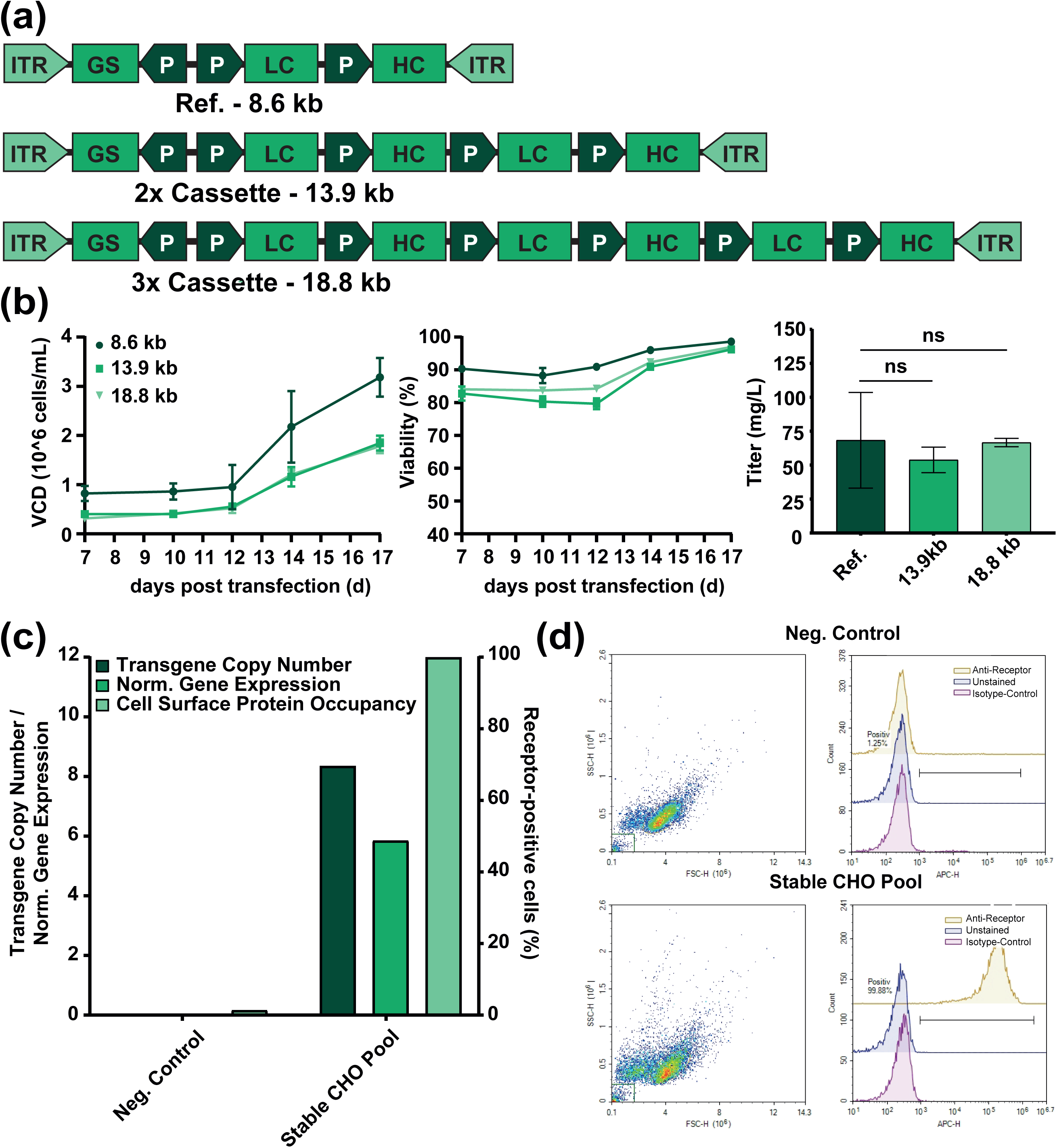
*Acyrthosiphon pisum* (AP) transposase system demonstrates high efficiency and versatility in transposing large vectors and maintaining transposition efficiency with increased cargo sizes. (a) Transposons with increased cargo sizes (13.9 kb and 18.8 kb) were compared against a standard antibody transposon with an 8.6 kb cargo. ITR = Inverted Terminal Repeat, GS = Glutamine Synthetase, P = Promoter, LC = Light Chain and HC = Heavy Chain. (b) Monitoring of cell growth and viability during the selection process after transfection of plasmids with different cargo sizes. Data for antibody production was generated after shake flask cultivation for 3 days on batch mode. Error bars indicate standard deviation from two biological replicates. (c) Genetic characterization and cell surface staining of stable CHO pool No. 12 (Table S2) co-transfected with a transposon vector encoding for a 100 kDa human surface receptor and the AP transposase. Gene expression values were normalized to endogenous *Eif3i* expression. (d) Fluorescence-Activated Cell Sorting (FACS) of the transfected pool (bottom) and the host control (top). For both samples cells were either unstained or stained with an anti-receptor antibody or an isotype control.

To extend the versatility of the novel AP transposase system to further biotechnological applications such as the development of cell lines for bioassays, stable CHO pools overexpressing a 100 kDa human cell surface receptor were established via transposition. Following the selection process, genetic characterization of stable pools confirmed overexpression of the heterologous gene of the transgene compared to the endogenous housekeeping gene and allowed for the determination of the number of integrated vectors (Table S2). For further investigation, one of the pools with relatively medium expression levels and copy number (Fig. 6c) was selected for cell surface staining of the expressed receptor in compared to the non-transfected host cells using flow cytometry (Fig. 6d). This analysis revealed that nearly 100% of the transfected cells showed a positive and strong cell surface receptor expression, further confirming the high efficiency of the engineered transposase system and its broad applicability. This clear distinction between the controls and the cells stained for the recombinant cell surface receptor provides robust validation of the efficiency of the novel AP transposase system and confirms its capacity to also facilitate the generation of recombinant cell lines for bioassay applications.

In conclusion, the AP transposase system demonstrated functional efficiency and comparable productivity, even with increased cargo sizes and for expression of large heterologous cell surface receptor proteins. This suggests the potential for inserting even larger genes into mammalian cell lines.

### Expanding the application of the novel AP transposase system to Adeno-Associated Virus production in human cells

After investigating the AP transposase system in the context of CHO cells and antibody production, we expanded our research to its application in Human Embryonic Kidney 293 cells (HEK293) cells, a human cell culture system widely used in viral vector production. This system is crucial for the development of advanced therapy medicinal products (ATMPs) such as Adeno-Associated Viruses (AAV). Our goal was to develop a HEK293-based AAV assay cell line, specifically for quantifying infectious AAV particles rather than AAV production, thereby showcasing the versatility of the novel AP transposase system across different host cell species. Traditional assays for determining infectious AAV particles typically use AAV2 rep and cap-expressing HeLa cells, which are transduced with an AAV vector in the presence of human adenovirus type 5. Since AAV vectors do not induce a cytopathic effect (CPE), co-infection with a helper virus is necessary to induce AAV genome replication and measure infectious events. Lee et al. previously reported the generation of an inducible cell line capable of replicating the AAV genome without the need for a helper virus, significantly simplifying the assay setup ^44^.

Building on this concept, we generated HEK293 cells expressing all the genes necessary for AAV genome replication (e2a, e4orf6, and rep68) using the AP transposase system. HEK293 cells were an ideal candidate for this work due to their widespread use and adaptability in the viral vector field. For cell line engineering, we followed a sequential approach as shown in Fig. 7a: To mitigate toxicity of viral genes, all genes were controlled by an inducible promoter. First, an inducible activator was introduced into HEK293 cells, creating HEK293-Ind cells. After antibiotic selection, the e2a gene was inserted (resulting in the intermediate cell line “HEK293-Ind-E2A”), followed by insertion of e4orf6 and rep68, resulting in the final cell line: “HEK293-Ind-E2A-E4-Rep68”. In all three transfections, transposase-encoding pDNA was co-transfected with the respective transposons.

**Figure 7:**
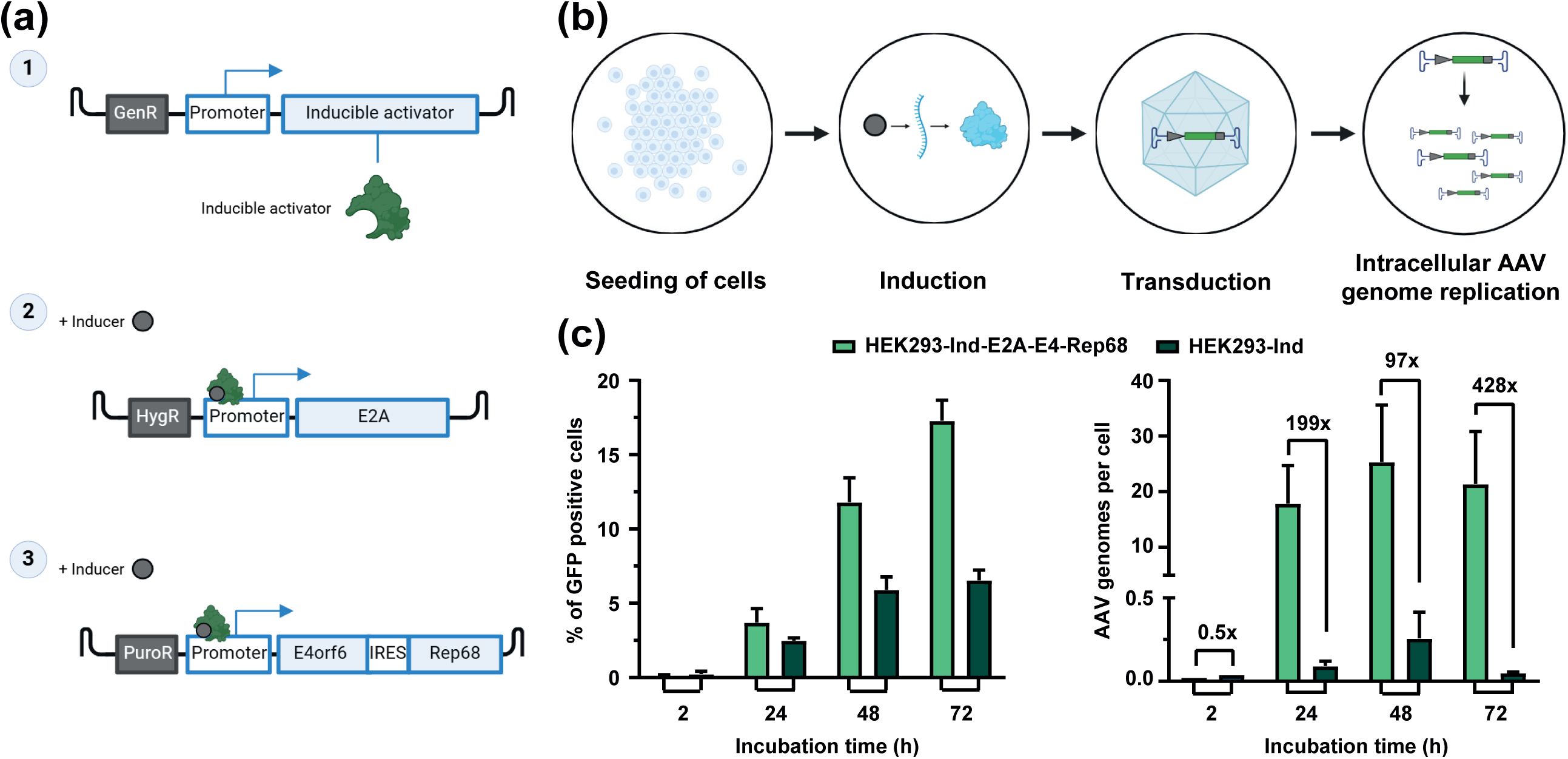
Genetic engineering of HEK293 cells using the *Acyrthosiphon pisum* transposase system. (a) In order to generate Adeno-associated virus (AAV) genome replicating HEK293-based assay cells, different Gene of Interests (GoI) were integrated into the host cell genome in a sequential approach: (1) a gene encoding an inducible activator including a Geneticin resistance gene (GenR), (2) e2a in combination with a hygromycin resistance gene (HygR) and (3) e4orf6 linked to rep68 via an IRES in combination with a Puromycin (PuroR) resistance gene. Upon three individual selection rounds, the resulting inducible HEK293-Ind-E2A-E4-Rep68 cells were used in an AAV genome replication assay (b). Therefore, expression of the integrated genes was induced by adding the respective inducer 4 hours after seeding into 6-well plates. 20 h later, the cells were transduced with an rAAV2-GFP vector (10 vector genomes / cell). (c) Cells were harvested at the indicated time points and analyzed for Green Fluorescent Protein (GFP) expression using flow cytometry and for AAV genome replication by digital PCR. Error bars indicate standard deviation from three biological replicates.

Finally, we validated the engineered cell line in an AAV genome replication assay. The cells were induced to express all genes of interest and subsequently transduced with a GFP-encoding AAV2 vector (Fig. 7b). Green Fluorescent Protein (GFP) expression and the number of AAV genomes per cell were monitored over 72 hours, demonstrating effective AAV genome replication in the engineered cell line compared to a control cell line (“HEK293-Ind”; Fig. 7c). This experiment showcased the adaptability of the novel AP transposase system and its species-agnostic application, indicating numerous potential future biotechnological applications

## Discussion

Functionally active transposase/transposon systems, although rarely found in nature, hold immense value for biomedicine and biotechnology. Especially the field of biopharmaceutical cell line development has greatly benefited from the discovery and implementation of these ancient molecular machineries ^5–7^. By introducing a multi-layered, combining a metagenomic screen with structural prediction, we introduced a novel and highly active chimeric transposase/transposon system, constructed *de-novo* from two different aphid species: *Acyrthosiphon pisum* and *Aphis craccivora*. This system was demonstrated to be suitable for cell line development and holds potential for broader applications, such as stable AAV producer cell lines due to its activity in human cell lines ^45^.

The search for novel and functional transposases is exceptionally challenging. Despite employing a comprehensive metagenomic screen and a stringent selection process that included both sequence assessments and structural analysis of the predicted dimeric scaffold, only two of the screened transposases were confirmed to be functionally active. Incorporating structural prediction tools in the selection process can effectively eliminate non-optimal candidates, as evidenced by our findings that the functional transposases in our study consistently adopt highly similar structures. Interestingly, although we aimed to enforce species diversity in our selection process, the only two functional transposases identified were derived from closely related insect species. These findings emphasize that most transposable elements found in genomes across species have likely lost their activity through various mechanisms and remain genetic fossils that are difficult to resurrect. Furthermore, consistent with other functional transposases, insects appear to be particularly valuable sources for identifying active systems, suggesting that insect genomes may have a higher tolerance for mobile genetic elements and overall genome plasticity. These findings align with other reports where *piggyBac*-like elements, identifiable in most species, were screened ^27,46^. However, such screens often yield a limited number of active transposases, with a best-case scenario of approximately 40%, as reported by Zhang et al. in 2024. This was echoed in our study, where only two out of seven evaluated systems were functional in mammalian cell lines. Other instances have employed an orthogonal approach to resurrect fossil transposases, resulting in the *Sleeping Beauty* Tc1/mariner-type transposase ^47^.

Although the wildtype transposases from *Acyrthosiphon pisum* and *Aphis craccivora* exhibit initial activity, we demonstrated several optimization strategies to enhance overall transposition activity. Our data suggest that internal natural NLS sequences might be sufficient to drive low levels of activity. However, engineering a more efficient nuclear import is crucial for increasing overall transposition efficiency. Furthermore, for optimizing recombinant protein expression, combining a transposase with a transposon containing an appropriate and stringent selection marker can significantly enhance efficiency. For instance, in cell line development approaches, the use of attenuated metabolic selection markers significantly increases overall protein expression levels ^41^. Through mutational protein engineering, we developed a hyperactive transposase variant and provided structural rationale for the impact of the mutations on its conformational flexibility. While the hyperactivity-enhancing mutations appear to increase flexibility, particularly in the catalytic domain of the transposase, the overall structure remains largely unaffected. Thus, the observed hyperactivity seems to be driven more by changes in kinetics and conformational flexibility rather than by adopting an overall altered dimeric scaffold.

For newly discovered transposases, the exact mechanisms of DNA-transposition often may remain elusive. While the mechanisms from the *piggyBac* system are well-understood ^9,36^, new transposases from the same family can still exhibit differing characteristics with regards to ITR or DNA-binding, necessitating individual optimization ^48^. This may also apply to our chimeric AP-HA/AC system from *Acyrthosiphon pisum* and *Aphis craccivora,* as we have not elucidated the exact mechanism behind the functionality of a chimeric combination. Nevertheless, the data clearly showed that this chimeric strategy could yield improved performance in biotechnological applications. Given the high sequence homology observed between the transposase/transposon pairs from *Acyrthosiphon pisum* and *Aphis craccivora*, it is plausible that these elements share a common evolutionary origin – either through vertical inheritance from a shared ancestor or via horizontal transfer between species. Over evolutionary time, species-specific host defense mechanisms may have driven divergent trajectories ^2^, potentially leading to the accumulation of impairing mutations within the transposase coding regions or the ITR regions. Such mutations could differentially impact transposition efficiency, depending on each host’s regulatory landscape governing genome plasticity and transposon activity. Notably, codon usage between the two elements is nearly identical, which may support vertical transmission, although convergence through amelioration cannot be excluded. By combining components from both species, the chimeric system may circumvent species-specific limitations, leveraging the functional integrity retained in each element to restore or enhance transposition capability.

Moreover, we applied a metabolic selection procedure to dissect variations in the system, resulting in many improvements. From a mechanistic point of view, the chimeric combination had the most enhancing effects on selection behavior. One potential explanation for this could be that the availability and kinetics of transposition occur earlier and/or in more cells, thus being more effective. For instance, the levels of recombinant protein expression remained consistent, likely due to minimal differences in copy number or no preference in integration loci. Increasing transgene copy numbers and protein expression levels seems to be more effectively driven by modulation of the selection marker ^40^.

The presented study thoroughly characterized the distribution of integration sites in clonal cell lines, providing valuable insights into the integration preference of the system. The chimeric AP-HA/AC system behaves similarly to other well-characterized systems: integration preferably occurs at the TTAA-motif ^43^, and integration sites are predominantly located within intronic sequences ^42^ and spread across the genome ^16^. Transposases are also known for selecting transcriptionally active genes ^18^, a characteristic that may need further verification through transcriptomic analysis.

Further characterization involved the evaluation of a potential limit in cargo size of the transposase/transposon system. Other transposases tend to show decreased efficiencies with increased cargos ^46^ or even reported diminished activity for cargo sizes >8 kilobases ^49^. With maintaining a high transposition efficiency, indicating by evenly fast recovery of cell pools during selection, while cargo size was increased up to approx. 18 kilobases the AP-HA/AC system offers high cargo transposition capability in mammalian cells. Although currently not proven to reach >100 kilobases, already proven for the the *piggyBac* transposase ^50^. Besides the unknown limitations of the presented technology regarding cargo size, the application of this system as a gene-delivery vehicle and its potential use in germline transgenesis warrants further investigation.

## Supporting information

Supplemental Sequence Information

Integration Site Analysis

## Acknowledgements

The authors gratefully acknowledge the support by team members of Cell Line Development, Upstream Development and Downstream Development from Boehringer Ingelheim Pharma GmbH & Co. KG.

This research did not receive any specific grant from funding agencies in the public, commercial, or not-for-profit sectors. All this research was funded by Boehringer Ingelheim Pharma GmbH & Co.KG.

## Author Contributions

D.H., F.R., M.S. and R.N. performed study design. F.R. executed most of the CHO experiments. R.N. and E.W. performed HEK293 cell culture experiments. M.S. executed metagenomic screening and structural modelling. D.H., R.N. and B.L. performed data analysis. B.L. executed the integration site analysis. All authors wrote and approved of the final article. Figures were partially created with BioRender.com.

During the preparation of this work the authors used CHATGPT-4 in order to check for grammar and spelling. After using this tool/service, the authors reviewed and edited the content as needed and take full responsibility for the content of the published article.

## Methods

### Metagenomic Screening and Transposon Identification

To identify piggyBac-like transposons, we conducted a comprehensive *in-silico* screen using the NCBI BLAST database. Genomes from a broad range of species, including bacterial, archaeal, and eukaryotic organisms, were queried for transposase-like sequences using known piggyBac transposase proteins as search templates. Candidate genomic regions were analyzed for the presence of intact open reading frames (ORFs) encoding transposase-like proteins. Sequences were retained if they contained the conserved DDD catalytic motif and had an approximate length of 600 amino acids, consistent with canonical piggyBac transposases. In a second screening step, we manually inspected the genomic regions flanking each candidate transposase gene for the presence of TTAA target site duplications and repetitive DNA motifs at both the 5′ and 3′ ends. Only candidates exhibiting both TTAA motifs and flanking repetitive sequences were retained for further analysis. Protein sequences of the identified transposases were aligned to assess sequence conservation and guide downstream comparative analyses. Structural predictions were performed using AlphaFold to assess the presence of a dimeric fold based on predicted tertiary structure features.

### Structural Prediction via AlphaFold2

Protein complex structures were predicted using AlphaFold-Multimer via ColabFold (v1.5.5). Input sequences were provided in multi-chain FASTA format. Amber relaxation was applied to all models post-prediction. Structural comparisons were performed against experimentally determined complexes, with particular reference to PDB entry 6X68.

### Plasmid Construction

Codon optimized DNA sequences encoding for the transposase genes were synthetized (Twist Bioscience) and directly cloned into Boehringer Ingelheim proprietary expression plasmids. Transposase expression was driven by a CMV promoter, and all transposases harbored an N-terminal Flag tag. Transposon plasmids, including ITR sequences for transposase recognition, were generated internally using restriction enzymes (NEB) according to the manufacturer. Vectors for antibody expression were designed to include both light chain and heavy chain on the same plasmid, with expression of both chains also driven by CMV promoters. Expression of the metabolic selection marker was driven by a SV40 promoter. Transposon plasmids for HEK293 engineering were generated accordingly and are depicted in Figure 6a. All plasmid preparations were performed using the EndoFree Plasmid Maxi Kit (Qiagen) according to the manufacturer’s instructions, and plasmid concentration and purity were assessed by spectrophotometry using a NanoPhotometer N120 (Implen) and agarose gel electrophoresis. All plasmid constructs were sequence-verified by Sanger sequencing prior to use.

### Protein Engineering

Transposase mutants were synthesized by Twist Bioscience and directly cloned into a transposase expression plasmid under the control of a CMV promoter. Mutational design was guided by structural insights from an AlphaFold model, enabling rational targeting of key functional regions. Specific modifications aimed to enhance DNA-binding affinity, introduce additional disulfide bridges within the catalytic domain, and increase hydrophobicity at the C-terminus to promote dimerization.

In addition, family shuffling was employed to generate further diversity. Mutational designs were informed by conserved regions identified within the discovered transposase cluster, allowing recombination of homologous sequences to explore functional variation.

### Cell Culture

CHO-K1 cells were purchased from ATCC. Suspension adaption and generation of bi-allelic GS knock out was performed at Boehringer Ingelheim. Resulting proprietary glutamine synthetase deficient (GS^-/-^) CHO-K1 host cell system (BI-HEX^®^ CHO-K1GS^-/-^) was cultured in suspension in proprietary animal-component free and chemically defined cell culture medium. Cell culture maintenance was performed every 2–3 days with seeding densities between 0.3×10^6^ to 0.7×10^6^ cells per mL. The Vi-CELL BLU (Beckman Coulter) or Cedex Analyzer^TM^ (Roche Diagnostics) were used for cell counting. Cultures were incubated at 36.5 °C, 5% CO^2^ and with agitation at 120 rpm (50 mm orbit) in an orbital shaker incubator (Infors).

HEK293 cells were purchased from ATCC (CRL-1573). Cells were cultured adherently in DMEM (1x) + GlutaMAX^TM^-I – Dulbeccós Modified Eagle Medium with 4.5 g/L D-Glucose and Pyruvate (Gibco) and 10 % Fetal Bovine Serum (Gibco) in Nunc^TM^ EasYFlask^TM^ (Thermo Fisher Scientifi). Sub-cultivation was performed every 2-3 days. Therefore, cells were washed with PBS, detached from the flasks using TrypLE^TM^ Select (Gibco) and counted with aNucleoCounter^®^ NC-202^TM^ (ChemoMetec). Subsequently, cells were seeded to new Flasks with seeding density of 3×10^4^ cells/cm^2^.

### Generation of Stable CHO Cell Pools

BI-HEX®CHO-K1GS^-/-^ host cells were co-transfected with either 4 µg transposase-encoding mRNA (GenScript) or 1.5 µg transposase-encoding plasmid in combination with 5 µg transposon, carrying the genetic information for the transgene sequences (e.g. antibody) as well as glutamine synthetase gene serving as a selection marker. For transfection, the Neon™ transfection system in combination with the 100 µL Neon™ Kit was used for electroporation (Thermo Fisher Scientific) of 5×10^6^ cells. After the transfection, stable CHO cell pools were generated via metabolic selection in Boehringer Ingelheim proprietary selection medium devoid of L-glutamine and methionine sulfoximine. Selection pressure was applied one day post-transfection by complete media exchange and resuspension of cells in 40 mL medium. After reaching a viability of >70% and a VCC of >1.5×105 cells/mL, cells were transferred in 25 mL medium in a shaking flask and cultivated until reaching viabilities >95%.

### Generation of Clonal Cell Lines

Single cell cloning (SCC) of stable mAb producing cell pools was performed using the CellCelector® (Sartorius) device in combination with the nanowell-based cloning technology for the deposition of single cells. Cells were plated out in nanowells and the verification of single cell deposition was performed by microscopic imaging. On day 4 following deposition, additional images were taken to assess cell confluence, and the best-growing cells were transferred into 96-well plates for further cultivation. Cell productivity was subsequently determined by bio-layer interferometry using an Octet® HTX system (FortéBIO). Best growing and producing clones were selected for expansion into shake flasks.

### Generation of AAV genome replicating HEK293-based assay cell line

The generation of the HEK293 cell line involved a series of co-transfections using a Polyethylinimine (PEI)-based protocol. All steps involved the co-transfection of a plasmid of interest with a helper plasmid, which encoded the AP transposase.

In a sequential approach, three plasmids were introduced into the cells. All transfections were performed using the transfection Reagent FectoVIR (Polyplus Transfection) with a pDNA:FectoVIR ratio of 1:3 and 2 pg of pDNA per cell. Transposase-encoding pDNA was co-transfected in all transfections (2/3 Transposon and 1/3 transposase-encoding pDNA).

Firstly, a plasmid encoding an inducible activator was transfected to enable the use of an inducible promoter. The transfected cells were then selected using 100 µg/mL of Geneticin.

Subsequently, *e2a*, controlled by an inducible promoter, was introduced alongside a hygromycin resistance gene. The selection of these cells was achieved using a combination of 100 µg/mL Geneticin and 125 µg/mL Hygromycin B.

In the final step, *e4orf6* and *rep68* (linked via an IRES) were introduced together with a puromycin resistance gene. For selection the cells were cultured in a medium containing 100 µg/mL Geneticin, 125 µg/mL Hygromycin B, and 0.8 µg/mL Puromycin.

The resulting cell pool is referred to as HEK-293-Ind-E2A-E4-Rep68.

### AAV genome replication assayes

The functionality of the generated HEK293-based assay cell line was verified by an AAV genome replication assay. To this end, HEK293-Ind-E2A-E4-Rep68 cells were seeded in a 6-well plate (5×10^5^ cells/well) and incubated at 37°C for 4 h. Subsequently, the inducer was added to the cell culture and further incubated for 20 h. Then, cells were transduced with an rAAV2-GFP vector (10 vector genomes / cells). Upon AAV transduction, the cells were harvested after 2 h, 24 h, 48 h and 72 h and analyzed for GFP expression by flow cytometry using a NovoCyte Flow Cytometer System (Agilent) or for AAV genome replication using dPCR. For dPCR analysis, cells were centrifuged at 1000 × *g* for 5 minutes and the DNA was isolated with the DNeasy® Blood & Tissue Kit according to the manufactureŕs instruction (Qiagen). Subsequently, the number of AAV genomes was determined using dPCR with the QIAcuity One instrument (Qiagen) in combination with the dPCR CGT ITR (FAM) Assay (Qiagen) following the manufacturer’s instructions.

### Surface Staining for Receptor Expression Analysis

CHO pools prepared for FACS analysis by staining with APC-labelled anti-receptor antibody. Prior to staining, cells were incubated with an Fc-block solution. The Fc-block solution was prepared and used to block the cells, which were then incubated at room temperature for 10 minutes. The cells were then prepared for antibody staining. A total of 1.5×10^6^ cells were stained in a FACS buffer with antibodies. Unstained and isotype controls were included for each pool. The cells were incubated on a shaker at 4°C for 45 minutes in the dark. After incubation, the cells were washed three times with cold FACS buffer without EDTA and prepared for subsequent FACS sorting using the Novocyte 3005 Instrument (OLS OMNI Life Science).

### Determination of Antibody Concentration

Antibody concentration was determined by bio-layer interferometry using an Octet® HTX system (FortéBIO) with protein A sensors.

### Antibody purification and product quality analysis

Antibody purification from harvested, cell-free CCF was performed according to Boehringer Ingelheim standard downstream small-scale sample preparation applying a miniature protein A capture step (MabSelect SuRe, Cytiva). Product quality attributes were determined via size exclusion chromatography (SEC, 1290 Infinity II/ AgilentTechnologies; H-Class / Waters) and capillary gel electrophoresis (LabChip^®^/PerkinElmer). Charge heterogenicity was evaluated via a strong cation-exchange column (CEX,1260 Infinity II/ Agilent Technologies; Alliance/Waters) to identify main, acidic (APGs) and basic (BPGs) antibody fractions. N-glycosylation of the Fc part was analyzed by LabChip^®^ glycan profiling assay (PerkinElmer) and hydrophilic interaction liquid chromatography (OligoMap/ Shimadzu Nexera).

### Fed-Batch Bioprocess

Bioprocess performance of stable pools and clones was assessed in a small-scale bioreactor system (Sartorius). At the start of the process, cells were seeded with a target value of 0.7×10^6^ viable cells/mL. Starting at day 2 of the process, automatic feeding of 40 mL/L/day of nutrient feeds was started and glucose concentration was adapted manually to constantly exceed a concentration of 4 g/L. The pH was maintained at 6.95, and the partial pressure of oxygen was consistently controlled at 50%. Cell growth and metabolites were measured daily. Fed-batch runs were cultivated for up to 14 days and harvested via centrifugation and 0.2µm sterile filtration.

### Isolation of Genomic DNA and Total RNA

Nucleic acid material was isolated from a total of 1×10^7^ cells. The QIAsymphony DSP DNA Midi Kit was used to isolate gDNA on the QIAsymphony SP (both Qiagen) according to the manufacturer’s protocol. During the extraction procedure, an RNase treatment was performed to remove RNA. The QIAsymphony RNA Kit (Qiagen) was used to isolate total RNA according to the manufacturer’s protocol. During the extraction procedure, a DNase treatment was performed to remove genomic DNA. The concentration of DNA and RNA samples was determined by UV scan with a NanoPhotometer N120 (Implen) at A260nm.

### Hybridization-based Target Enrichment and Sequencing

To enrich the genomic regions of the cell lines in which vector integration has occurred, SureSelectXT HS2 DNA Kit (Agilent Technologies) was used according to the manufacturer’s protocol. Briefly, 200 ng of gDNA was diluted and enzymatically fragmented, resulting in a target fragment size of 180 to 250 bp. Subsequently, whole genome NGS libraries were prepared and hybridized with custom probes targeting the expression vector. The concentrations and size of final libraries were measured using the high sensitivity Quant iT™ dsDNA Assay Kit (Invitrogen) and 5200 Fragment Analyzer System (Agilent), respectively. For sequencing, alle samples were pooled and 5 µL library pool was denatured with 5 µL 0.2 M NaOH for 5 min at RT. Afterwards, the denatured library pool was neutralized with 5 µL 0.2 M Tris HCl and diluted to a final loading concentration of the library of 1.2 pM with HT1 hybridization buffer (NextSeq Accessory Box, Illumina). Furthermore, 1 % denatured PhiX-control 20 pM was spiked. 1300 µL of the final library pool were loaded into the NextSeq reagent cartridge and sequencing was performed with the NextSeq550 system (Illumina). Run parameters were set as followed: paired end sequencing mode with 150 cycles per read (2 x 150 bp) and 8 cycles per index read. Sequencing quality was rated based on the Q30 and the clusters passing filtering. The Q30 value implicates a base call accuracy of 99.9 %. The accepted quality specifications for sequencing runs required more than 80 % of the bases above Q30 and more than 70 % of the clusters pass filtering.

### Integration Site Analysis

Sequencing data was converted into FASTQ files using bcl2fastq (v2.20.0.422, Illumina) and analyzed with an in-house developed bash script. This script performs the following analytical steps:

Initial quality control of raw sequencing reads was performed using FastQC (v0.11.9) ^52^. Adapter sequences and low-quality bases were trimmed using Trimmomatic (v0.39), followed by a second round of quality control on the trimmed reads to ensure data integrity. Unique molecular indices (UMIs), incorporated during library preparation, were extracted using fgbio (v2.3.0), and a BAM file combining reads and UMIs was generated. Reads were then extracted from the BAM file using samtools (v1.9) and aligned to the Chinese hamster (Cricetulus griseus) reference genome CriGri-PICRH-1.0 ^53^ using bwa-mem2 (v2.2.1). Reads mapping to the same genomic position and sharing identical UMIs were grouped, and consensus sequences were generated using fgbio. These consensus reads were subsequently re-mapped to the CriGri-PICRH-1.0 genome, and low-quality mappings were filtered out using samtools. Reads aligning to both the Cricetulus griseus genome and the vector reference were extracted for downstream analysis. Structural variants were identified using lumpy (v0.2.13), and the resulting variants were analyzed for vector integration events, which were exported as a tabulated output.

Further examination of vector integration sites was performed using RStudio (v2022.02.3 Build 492). The analysis pipeline classified vector breakpoints into three categories: “left” ITR, “right” ITR, or “unknown” based on their genomic context and orientation. To improve specificity, artifacts located in close proximity to true integration events but exhibiting significantly lower read counts were removed. Breakpoints corresponding to the same vector-genome junction were paired by assessing their spatial proximity and matching breakpoint classifications. Gene annotations and sequence motifs at the integration sites were extracted from the reference genome to provide functional context. To ensure robustness, hits were filtered based on normalized read counts, retaining only those with at least 10% of the maximum read count for a given breakpoint. For breakpoint pairs, at least one member of the pair was required to meet this threshold. All visualizations were generated using RStudio to support interpretation and reporting of integration patterns.

### Droplet Digital PCR

Gene expression analysis and transgene copy number determination was performed via droplet digital Polymerase Chain Reaction (ddPCR) on a QX200 system (BioRad). Primer-probe sets (Table S3) were designed to specifically prime to an endogenous housekeeper control (e.g. Eif3i) and the target genes (e.g. LC and HC). Each hydrolysis probe was labelled with a reporter fluorescence dye (FAM for target genes and HEX or VIC for endogenous control).

For the separation of potential tandem copies of the expression vector, a restriction digest of the 750 ng gDNA was performed for copy number determination according to the manufactureŕs instructions (NEB). After formation of droplets using a QX200 AutoDG (BioRad) according to the manufacturer’s instructions, cycling was carried out in a standard PCR cycler under the following conditions for ddPCR: 95 °C for 10 min followed by 50 cycles at 94 °C for 30 s and 59 °C for 60 s and a final step of 98 °C for 10 min. For One-Step RT-ddPCR to analyze gene expression, the protocol was initiated by a single step of 50 °C for 10 min followed by the previously described cycling program with adaption of the annealing temperature to 56 °C. For gene expression analysis 500 pg of total RNA was used as template for the ddPCR reaction. Amplified samples were analyzed using a QX200 droplet reader (BioRad, Germany) for the determination of total fluorescent droplet counts.

Based on the absolute quantification of target molecules per reaction, the gene copy number per cell was determined as the ratio between transgenic and endogenous copy number per reaction (corresponding to the copy number per haploid genome) and then multiplied by 2 to express the copy number per cell. Normalized gene expression values were determined as the ratio between transgenic and endogenous target molecule concentration per reaction.

### Southern Blot Analysis

Prior to Southern Blot analysis, 1000 ng of gDNA and diluted amounts of vector DNA were digested with restriction enzymes according to the manufactureŕs instructions (NEB, Germany). Digested gDNA samples were separated on TAE-agarose gel by electrophoresis. For size determination, a DIG-labelled DNA Molecular Weight Marker VII (Roche Diagnostics) was loaded as well. After depurination and a neutralization, transfer of DNA onto positively charged nylon membrane (Roche Diagnostics) was carried out by capillary forces during overnight incubation.

For Southern blot hybridization, the hybridization solution from the DIG Easy Hyb Kit (Roche Diagnostics) was used according to protocol. Labelling of DNA probes with Digoxigenin-Deoxyuridine Triphosphate (DIG-dUTP) was performed utilizing the PCR DIG Probe Synthesis Kit (Roche Diagnostics). The PCR probes were generated by using the respective expression vector as template in the PCR reaction. Hybridization with the DIG-labelled probe was then performed over night at 42 °C. Washing and blocking of membranes was done using the DIG Wash and Block Buffer Set (Roche Diagnostics) according to protocol. For immunological detection the membrane was incubated with an anti-digoxigenin alkaline phosphatase conjugate (Roche Diagnostics). The CDP-Star Detection Reagent (GE Healthcare) was used for detection of the hybridization signals according to protocol. Subsequently, images of the blot membranes were acquired on a ChemiDoc Touch system (Bio-Rad) using the optimal auto-exposure function. Processing and labelling of the resulting images was performed in the Image Lab Software (Bio-Rad, Germany) Version 6.1.0 (build7).

### Data Analysis

Data visualization was performed using GraphPad Prism (GraphPad) and data are presented as mean ±□standard deviation. Number of replicates are indicated in the respective figure legends. Unpaired t-test was used to assess significant differences between results and *P*-values of *□*P□*<□0.05, **□*P□*<□0.01, ***□*P□*<□0.001, and ****□*P□*<□0.0001 were considered significant.

