## Supplemental Information for "Discovery of a Novel Chimeric Transposase-Transposon System for Advanced Genome Engineering"

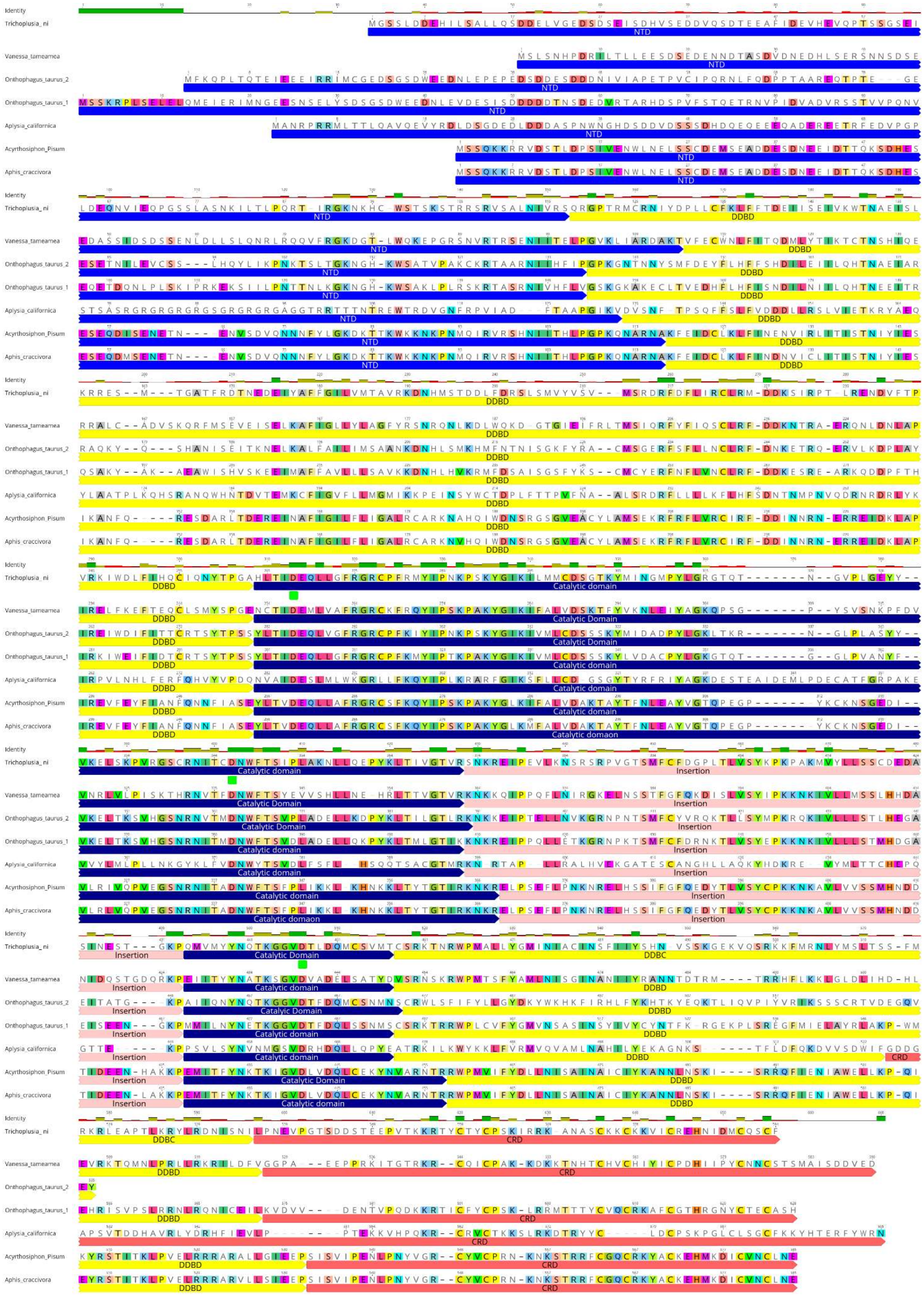

Fig S1: Protein amino acid sequence alignment of identified transposases compared to the *piggyBac* transposase from *Trichoplusia ni*. Numbers above each sequence refer to the alignment position and also each individual transposase sequence is numbered. Homologous amino acid between all transposases are highlighted in colors. Catalytic DDD triad motif is highlighted in green in the *piggyBac* sequence. DDBD = dimerization and DNA-binding domain; CRD = cysteine-rich domain.

| | Trichoplusia_ ni | Vanessa_tameamea | Onthophagus_taurus_2 | Onthophagus_taurus_1 | Aplysia_californica | Acyrthosiphon_ Pisum | Aphis_craccivora |
|---|---|---|---|---|---|---|---|
| Trichoplusia_ ni | | 31.37 | 39.58 | 43.10 | 20.39 | 33.28 | 33.45 |
| Vanessa_tameamea | 31.37 | | 30.55 | 31.86 | 22.37 | 39.43 | 39.26 |
| Onthophagus_taurus_2 | 39.58 | 30.55 | | 50.09 | 19.64 | 30.17 | 30.17 |
| Onthophagus_taurus_1 | 43.10 | 31.86 | 50.09 | | 20.10 | 32.89 | 33.22 |
| Aplysia_californica | 20.39 | 22.37 | 19.64 | 20.10 | | 22.08 | 22.24 |
| Acyrthosiphon_pisum | 33.28 | 39.43 | 30.17 | 32.89 | 22.08 | | 98.29 |
| Aphis_craccivora | 33.45 | 39.26 | 30.17 | 33.22 | 22.24 | 98.29 | |

Fig S2: Sequence identity (%) of selected transposase full-length amino acid sequences.

| Structure | RMSD | Dimeric Scaffold |
|---|---|---|
| 6x68 (PDB)[1] | Ref. | 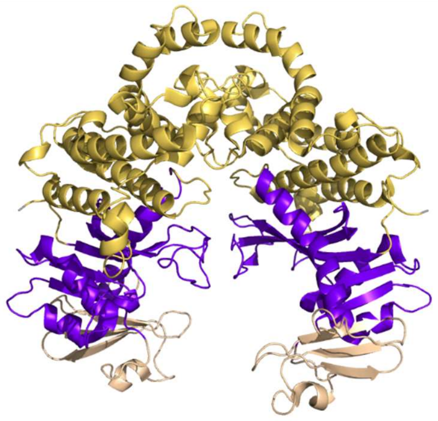 |
| *Trichoplusia ni* Wildtype | 1.22 | 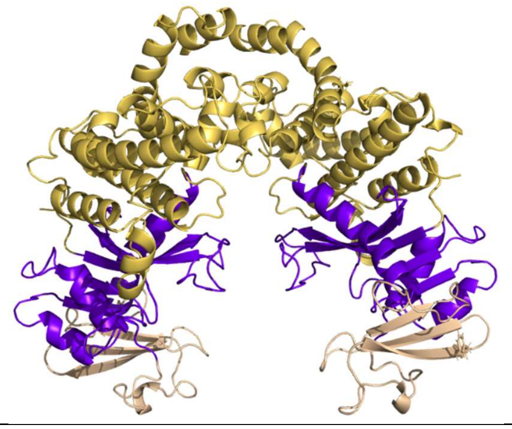 |

| | | |
|---|---|---|
| *Acyrthosiphon pisum* Wildtyp | 1.28 | 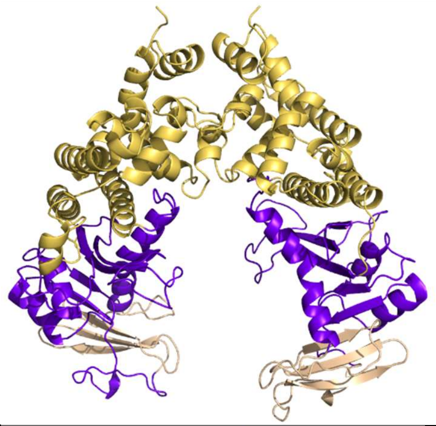 |
| *Acyrthosiphon pisum* Hyperactive Variant | 1.30 | 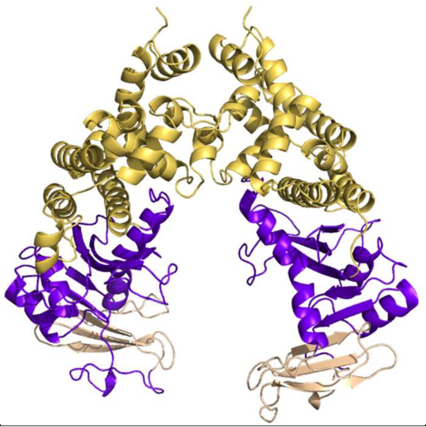 |
| *Aphis craccivora* | 1.32 | 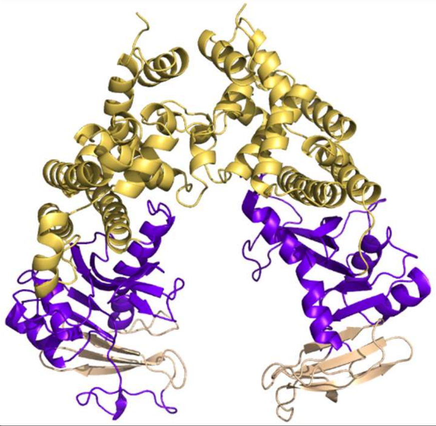 |
| *Vanessa tameamea* | 1.24 | 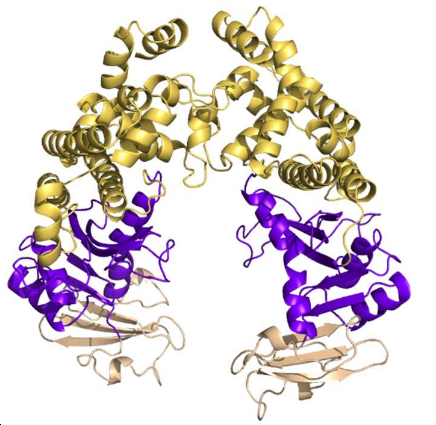 |

| | | |
|---|---|---|
| *Aplysia californica* | 1.45 | 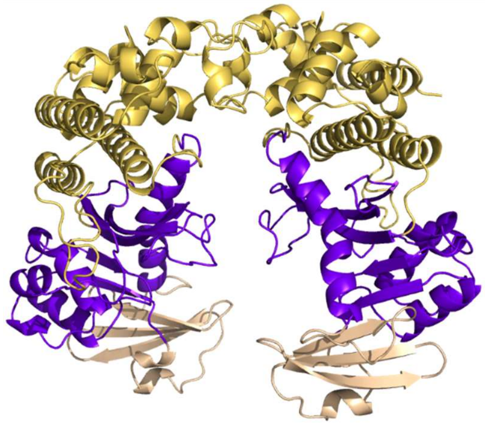 |
| *Onthophagus taurus* 1 | 1.10 | 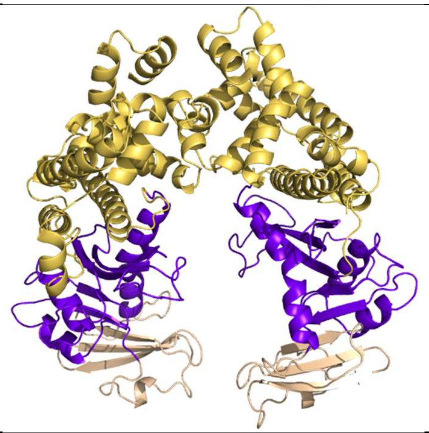 |
| *Onthophagus taurus* 2 | 1.36 | 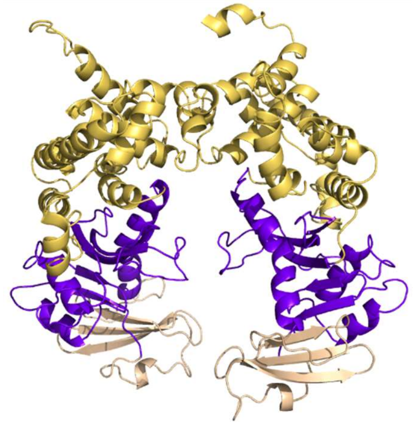 - No predicted CRD |

Fig S3: Structural Analysis of Dimeric Transposase Scaffolds. The Alphafold-based prediction, excluding the unstructured N-terminal Domain (NTD) and Cysteine-rich Domain (CRD), was aligned to the cryo-EM structure of *T.ni* (PDB ID: 6x68). The depicted structure illustrates the dimeric configuration.

| Structure | Monomeric Overlay | Monomeric Overlay 180° turn |
|---|---|---|
| *Trichoplusia ni* Wildtype | 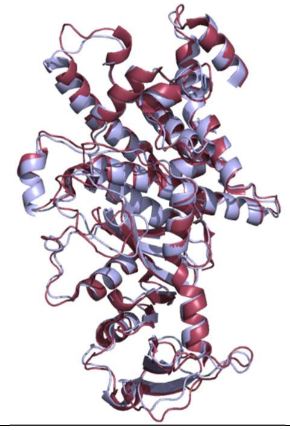 | 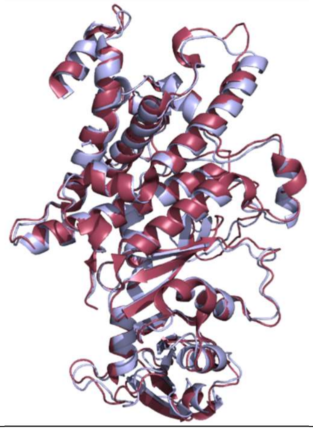 |
| *Acyrthosiphon pisum* Wildtyp | 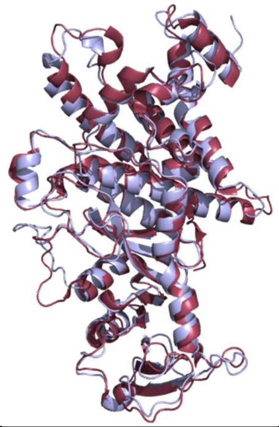 | 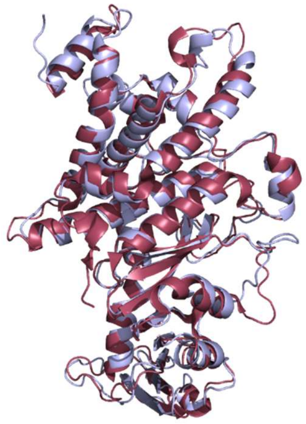 |
| *Aphis craccivora* | 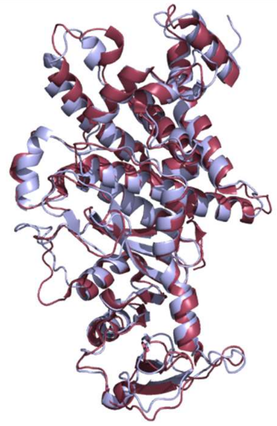 | 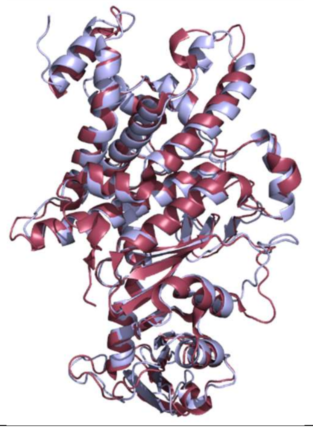 |

| | | |
|---|---|---|
| *Vanessa tameamea* | 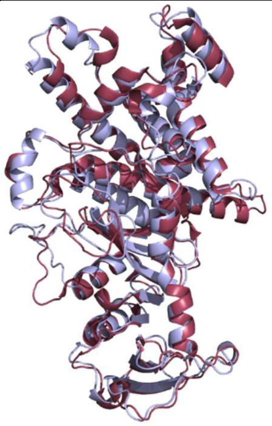 | 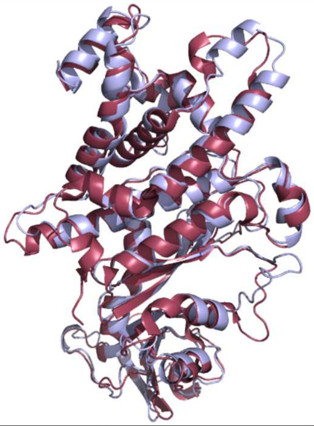 |
| *Aplysia californica* | 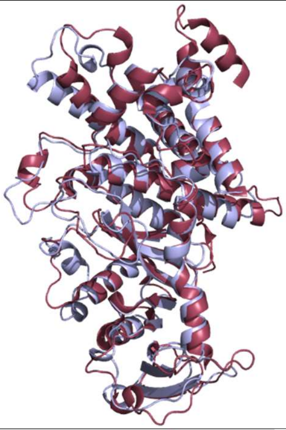 | 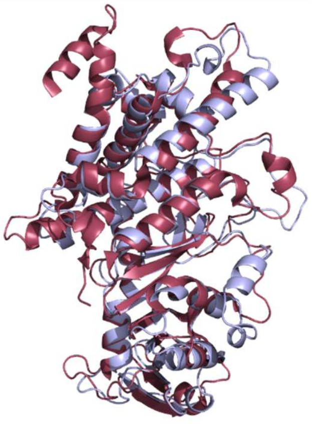 |
| *Onthophagus taurus* 1 | 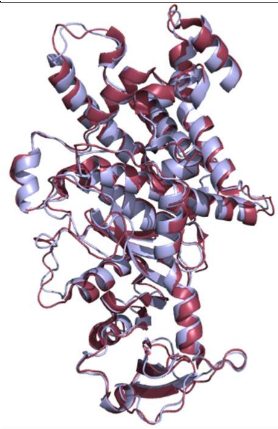 | 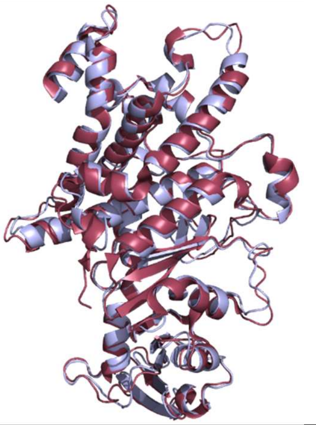 |

| | | |
|---|---|---|
| *Onthophagus taurus* 2 | 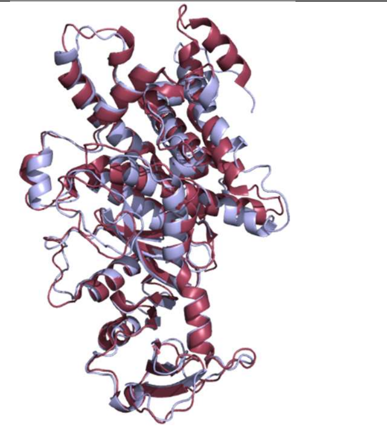 | 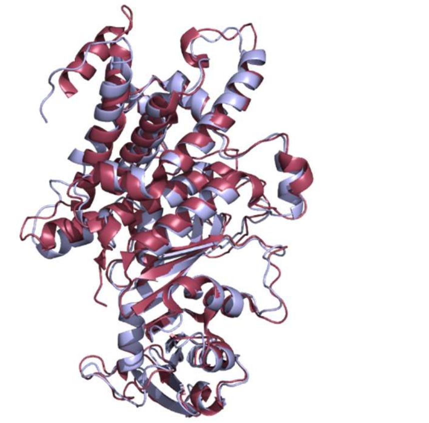 |

Fig S4: Structural Analysis of Monomeric Transposase Scaffolds. The Alphafold-based prediction, excluding the unstructured N-terminal Domain (NtD) and C-terminal Regulatory Domain (CRD), was aligned to the cryo-EM structure of *T.ni* (PDB ID: 6x68). The depicted overlays show the reference structure (6x68) in raspberry red and the investigated transposases in light blue.

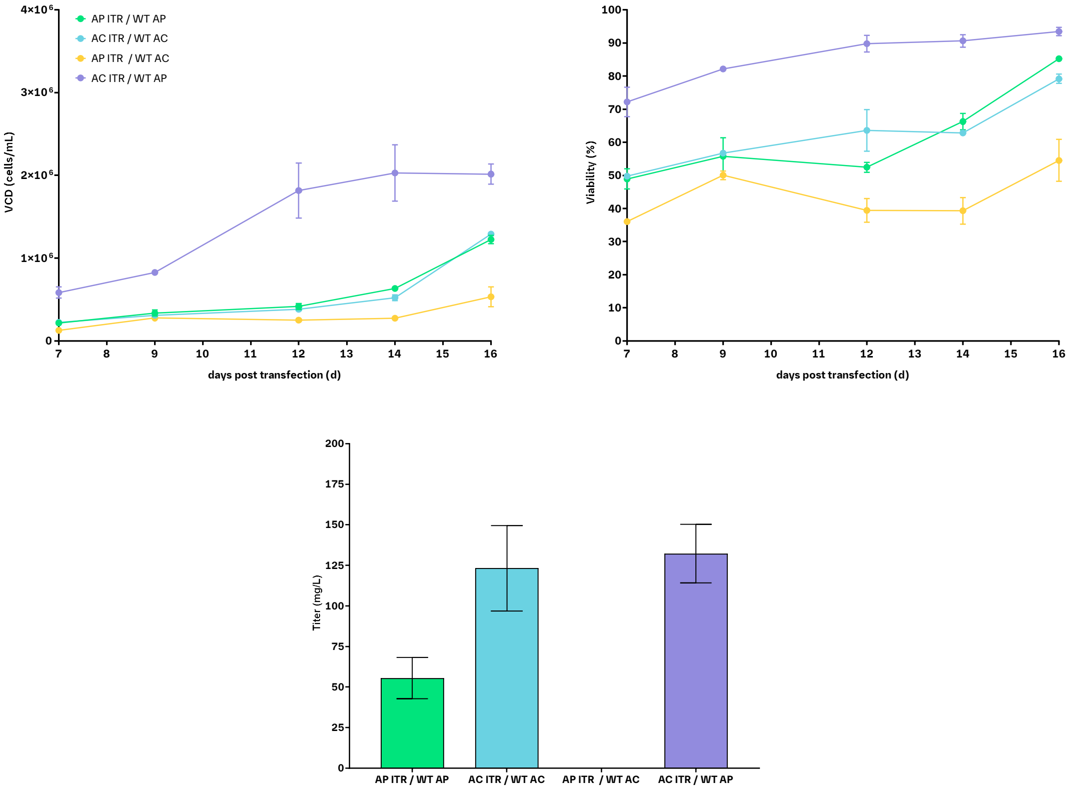

Fig S5: Monitoring of selection-marker based survival and recombinant protein expression following the transfection of GS-deficient CHO cells following cross combination of transposase protein from wildtype (WT) *Acyrtosiphon pisum* (AP) with Inverted Terminal Repeat (ITR) sequences from *Aphis craccivora* (AC) and *vice versa*. Top: Viable cell density (VCD) and cell viability in course of the metabolic selection process in L-glutamine-free medium. Bottom: Antibody concentration at the end of a 3 day batch cultivation. Error bars indicate standard deviation from two biological replicates.

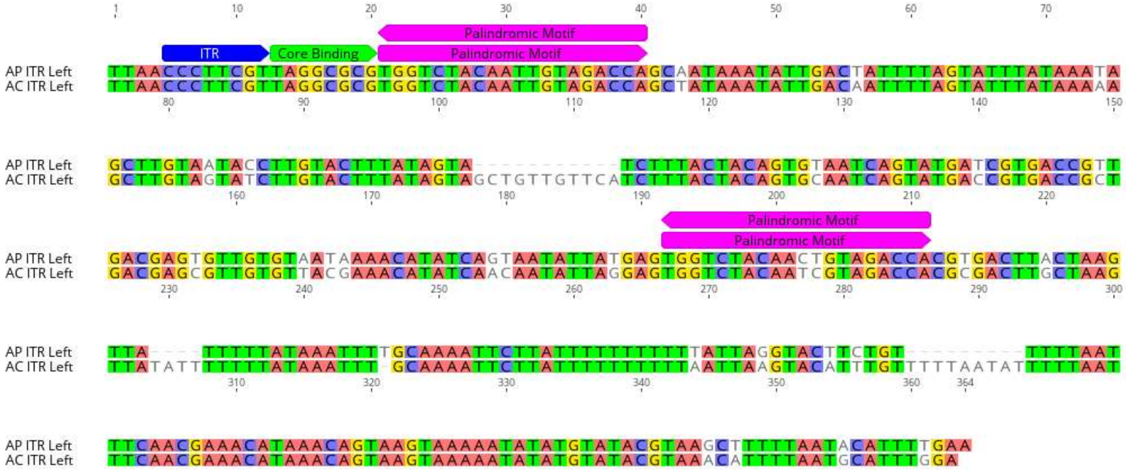

Fig S6: Sequence alignment of left Inverted Terminal Repeat (ITR) sequences of functionally active transposase/transposon system discovered in *Acyrthosiphon pisum* (AP) and *Aphis craccivora* (AC). Sequence similarities between two species are highlighted.

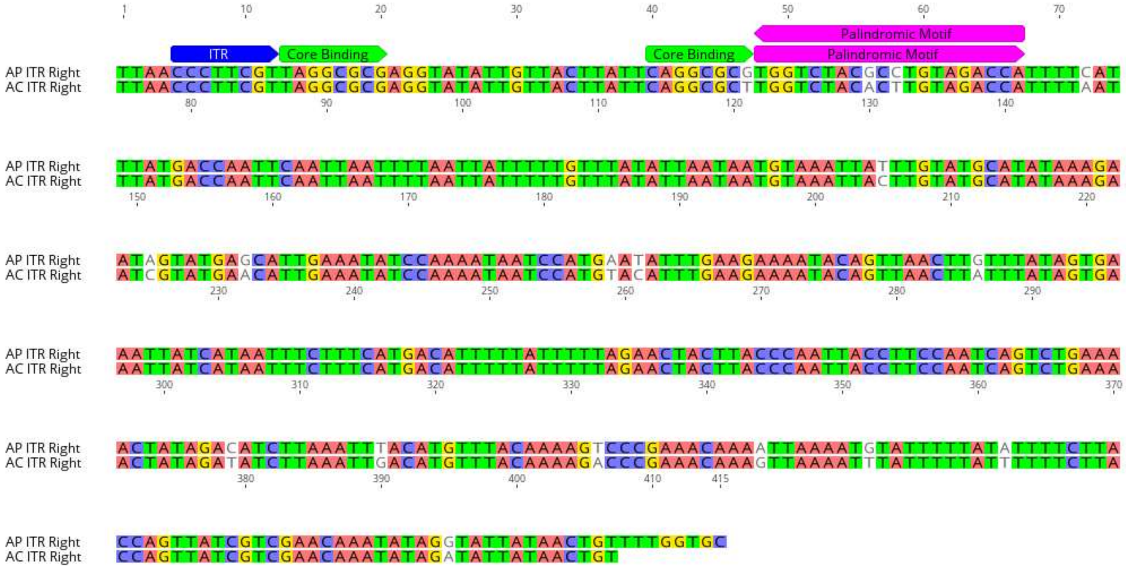

Fig S7: Sequence Alignment of right Inverted Terminal Repeat (ITR) sequences of functionally active transposase/transposon system discovered in *Acyrthosiphon pisum* (AP) and *Aphis craccivora* (AC). Sequence similarities between two species are highlighted.

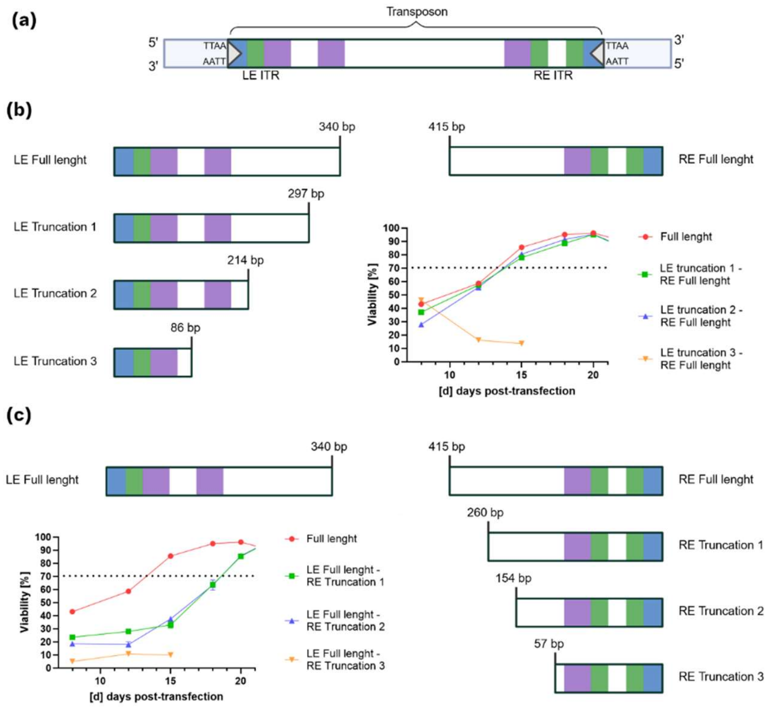

Figure S8: Truncation of the *Acyrthosiphon pisum* (AP) transposon flanking regions outside the core domains of left end (LE) and right (RE) Inverted Terminal Repeats (ITR) sequences consistently reduces transposition activity. (a) Schematic representation of the LE and RE ITR sequences of the *Acyrthosiphon pisum* AP and *Aphis craccivora* AC transposon sequences. Predicted core transposase binding sites are highlighted in green, palindromic motifs predicted to be recognized by the cysteine-rich domain dimer are shown in purple, and the conserved terminal motif is marked in blue. Lowercase letters indicate sequence differences between the AC and AP variants. (b) Systematic truncation of LE sequence (while keeping RE at full length) and impact on transposase activity as measured from selection behavior. (c) Systematic truncation of RE sequence (while keeping LE at full length) and impact on transposase activity as measured from selection behavior.

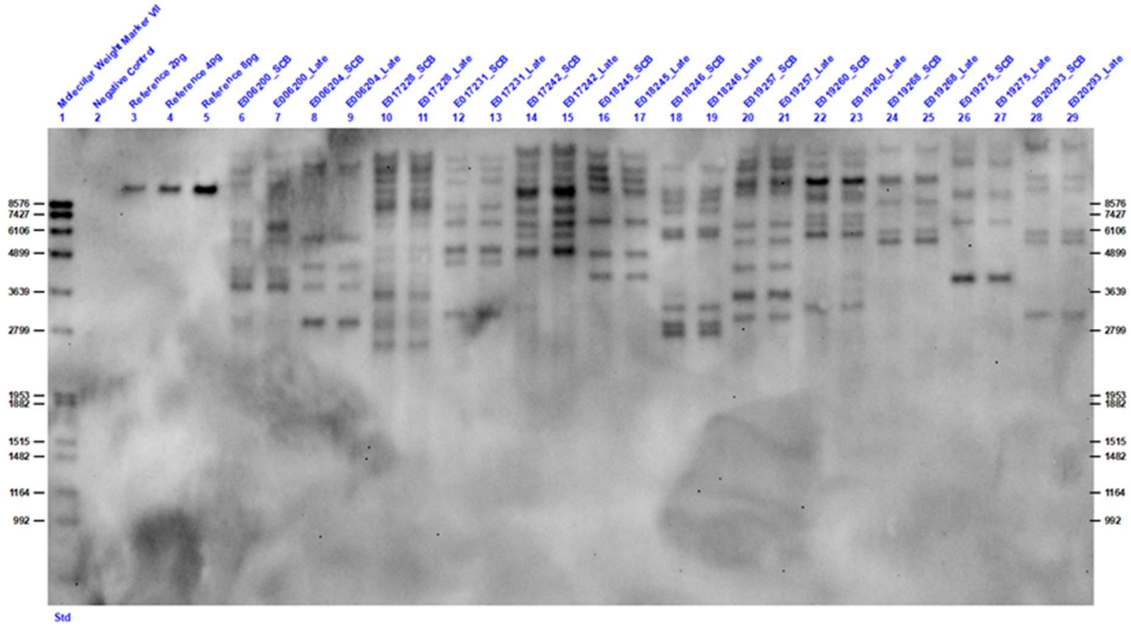

Fig S9: Genetic stability analysis of transposase-derived clonal cell lines via Southern blot analysis using hybridization probes to specifically target the LC. Genomic DNA was digested prior to analysis with restriction enzyme *HindIII*, cutting once within the expression vector. Cryopreserved cells (SCB) were compared to cells harvested after long term cultivation for 52 days (Late), clone names are indicated by individual numbers.

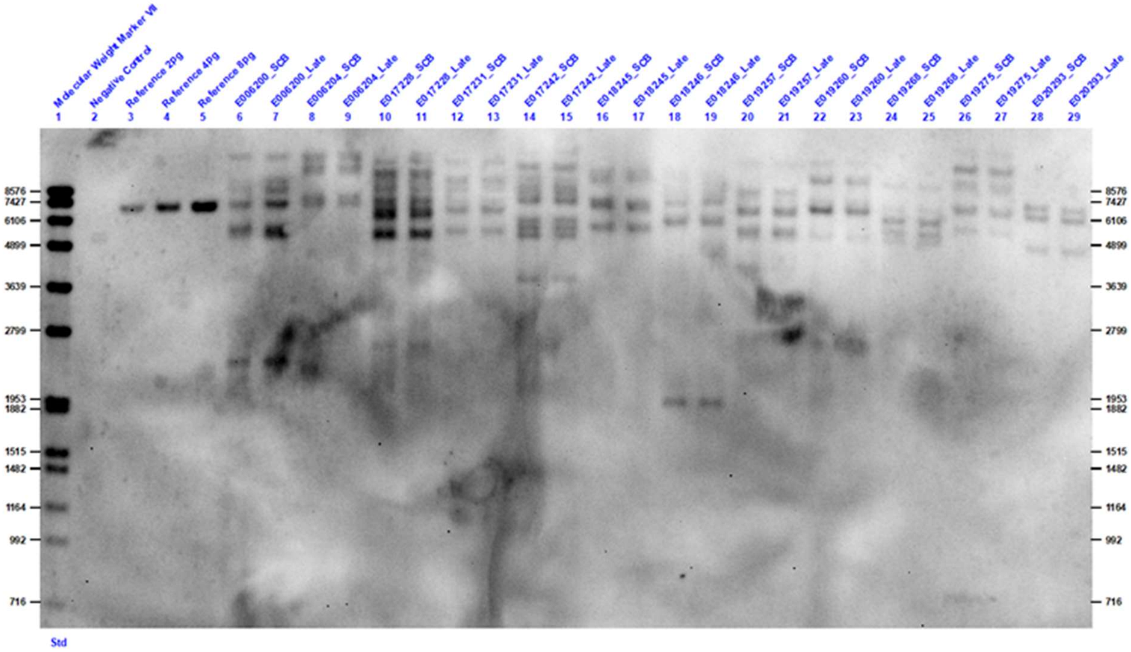

Fig. S10: Genetic stability analysis of transposase-derived clonal cell lines via Southern blot analysis using hybridization probes to specifically target the HC. Genomic DNA was digested prior to analysis with restriction enzyme *HindIII*, cutting once within the expression vector. Cryopreserved cells (SCB) were compared to cells harvested after long term cultivation for 52 days (Late), clone names are indicated by individual numbers.

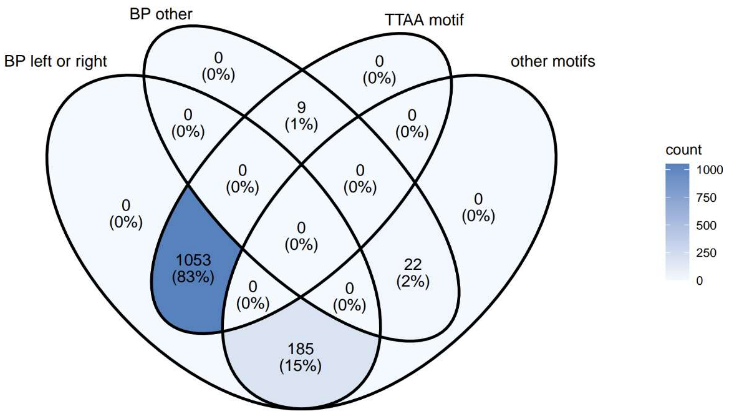

Fig. S11: Venn diagram visualizing the identified motif and vector breakpoints from 648 vector integration sites identified in clonal CHO cell lines following transfection using the *Acyrtosiphon pisum* transposase. Each circle represents the motif distribution at the identified vector breakpoint (BP), with the overlapping region indicating motifs common to both. Left and right BP correspond to the transposon ends.

Table S1: Individual and combinatorial screening of mutations to improve the selection behaviour and productivity of CHO cell pools generated using the *Acyrtosiphon pisum* (AP) transposase. "N/A": variant dysfunctional. "++": variant with strongly improved characteristics as compared to AP wild type. "+": variant with improved characteristics as compared to AP wild type. "0": comparable to AP transposase wild type. "-": variant with decreased characteristics compared to AP wild type "—": variant with strongly decreased characteristics compared to AP wild type

| Variant | Selection behavior | Productivity |
|---|---|---|
| E428Q | N/A | N/A |
| K87Y | ++ | + |
| H100E | N/A | N/A |
| A264S | 0 | 0 |
| F287L | N/A | N/A |
| F238K | N/A | N/A |
| Q445E | N/A | N/A |
| L25V | N/A | N/A |
| N40T | N/A | N/A |
| L123F | N/A | N/A |
| I168F | N/A | N/A |
| V258I | N/A | N/A |
| A264G | N/A | N/A |

| | | |
|---|---|---|
| S270P | -- | 0 |
| Q273V | + | 0 |
| V212I + I215L | + | + |
| S277N + L284I | 0 | 0 |
| F345I + I348A | N/A | N/A |
| I363V + K365S | ++ | + |
| F76H + L78T + G79S | N/A | N/A |
| Q261M | - | -- |
| I386V | N/A | N/A |
| E391K + Y393I | N/A | N/A |
| I438A | -- | -- |
| K87Y + Q273V | + | + |
| K87Y + A264S + Q273V | ++ | 0 |
| A264S + Q273V | ++ | + |
| K87Y + A264S + S270P + Q273V | - | - |
| A264S + S270P + Q273V | 0 | 0 |
| S270P + Q273V | ++ | 0 |
| K87Y + A264S | ++ | + |
| L583F | + | + |
| K576I | + | 0 |
| Delete N584 + E585 | ++ | + |
| I578F | 0 | - |
| K206D + R209D | N/A | N/A |
| N333C + A338C | N/A | N/A |
| S372E | 0 | + |
| Q492K | 0 | - |
| S277N | 0 | + |
| T306K | 0 | - |

| | | |
|---|---|---|
| A338F | N/A | N/A |
| D577F | - | 0 |

Table S2: Copy number and norm. gene expression values from stable CHO pools overexpressing a a 100 kDa human surface receptor. Gene expression values were normalized to endogenous Eif3i expression. Pool 12 was used for cell surface staining.

| Pool No. | Copies / cell | Norm. Expression |
|---|---|---|
| 1 | 5.88 | 4.13 |
| 2 | 4.95 | 3.77 |
| 3 | 1.48 | 1.11 |
| 4 | 8.08 | 1.47 |
| 5 | 1.48 | 0.73 |
| 6 | 7.63 | 8.57 |
| 7 | 10.52 | 19.53 |
| 8 | 5.68 | 7.67 |
| 9 | 6.63 | 9.48 |
| 10 | 4.44 | 2.45 |
| 11 | 3.66 | 2.65 |
| **12** | **8.09** | **5.84** |
| 13 | 8.34 | 5.89 |

Table S3: Primer and probe sequences used for ddPCR

| Oligo | Sequence 5' -> 3' | Target |
|---|---|---|
| Forward | ACGTCCGCATCCATTACTTC | Eif3i |
| Reverse | AAGAGTCTAAAACCCTCTTGAGCA | |
| Probe | TCCCCGGTGGATGGAGATCCAGCT | |
| Forward | CATCTTCCCACCTTCCGACG | IgG LC |
| Reverse | CTGCAGAGCATTGTCCACCT | |
| Probe | AGTCCGGCACAGCTTCTGTCGTGT | |
| Forward | CACACCTGTCCTCCATGTCC | IgG HC |
| Reverse | CCTCGTGAGACACATCCACC | |
| Probe | GCCCTTCCGTGTTTCTGTTCCCTCCA | |
| Forward | ACAGCCTGGAATGCACCTAC | Human cell surface receptor |
| Reverse | AAGCCGTCCCAGATTTCCAG | |

| | |
|---|---|
| Probe | CGACCTGGAACCTGACAGCAACCCT |

1. Chen, Q. *et al.* Structural basis of seamless excision and specific targeting by piggyBac transposase. *Nat Commun* **11**, 3446 (2020).
