## Supplemental Sequence Information for "Discovery of a Novel Chimeric Transposase-Transposon System for Advanced Genome Engineering"

**Supplemental Information – Sequence Listing**

>N_Terminal_NLS
MAPKKKRKVGIHGVPAA

>C_Terminal_NLS
KRPAATKKAGQAKKKK

>AP_Transposase_WT
MSSQKKRRVDSTLDPSIVENWLNELSSCDEMSEADDESDNEEIDTTQKSDHESESEQDISENETNEN
VSDVQNNNFYLGKDKTTKWKKNKPNMQIRVRSHNIITHLPGPKQNARNAKFEIDCLKLFINENVIRLITI
STNIYIESIKANFQRESDARLTDEREINAFIGILFLIGALRCARKNAHQIWDNSRGSGVEACYLAMSEKR
FRFLVRCIRFDDINNRNERREIDKLAPIREVFEYFIANFQNNFIASEYLTVDEQLLAFRGRCSFKQYIPSK
PAKYGLKIFALVDAKTAYTFNLEAYVGTQPEGPYKCKNSGEDIVLRIVQPVEGSNRNITADNWFTSFPL
IKKLKHNKKLTYTGTIRKNKRELPSEFLPNKNRELHSSIFGFQEDYTLVSYCPKKNKAVLVVSSMHNDD
TIDEENHAKKPEMITFYNKTKIGVDLVDQLCEKYNVARNTRRWPMVIFYDLLNISAINAICIYKANNLNS
KISRRQFIENIAWELLKPQIKYRSTITKLPVELRRRARALLGIEEPSISVIPENLPNYVGRCYVCPRNKNK
STRRFCGQCRKYACKEHMKDICVNCLNE

>AC_Transposase_WT
MSSQKKRRVDSTLDPSIVENWLNELSSCDEMSEADDESDNEEIDTTQKSDHESESEQDMSENETNE
NVSDVQNNNFYLGKDKTTKWKKNKPNMQIRVRSHNIITHLPGPKQNARNAKFEIDCLKLFINDNVICLI
TISTNIYIESIKANFQRESDARLTDEREINAFIGILFLIGALRCARKNVHQIWDNSRGSGVEACYLAMSEK
RFRFLVRCIRFDDINNRNERREIDKLAPIREVFEYFIANFQNNFIASEYLTVDEQLLAFRGRCSFKQYIP
SKPAKYGLKMFALVDAKTAYTFNLEAYVGTQPEGPYKCKNSGEDIVLRLVQPVEGSNRNITADNWFT
SFPLIKKLKHNKKLTYTGTIRKNKRELPSEFLPNKNRELHSSIFGFQEDYTLVSYCPKKNKAVLVVSSM
HNDDTIDEENLAKKPEMITFYNKTKIGVDLVDQLCEKYNVARNTRRWPMVIFYDLLNISAINAICIYKAN
NLNSKISRRQFIENIAWELLKPQIEYRSTITKLPVELRRRARVLLSIEEPSISVIPENLPNYVGRCYVCPR
NKNKSTRRFCGQCRKYACKEHMKDICVNCLNE

>AP_Transposase_HA
MSSQKKRRVDSTLDPSIVENWLNELSSCDEMSEADDESDNEEIDTTQKSDHESESEQDISENETNEN
VSDVQNNNFYLGKDKTTKWKKNKPNMQIRVRSHNIITHLPGPKQNARNAKFEIDCLKLFINENVIRLITI
STNIYIESIKANFQRESDARLTDEREINAFIGILFLIGALRCARKNAHQIWDNSRGSGVEACYLAMSEKR
FRFLIRCLRFDDINNRNERREIDKLAPIREVFEYFIANFQNNFIASEYLTVDEQLLAFRGRCSFKVYIPSK
PAKYGLKIFALVDAKTAYTFNLEAYVGTQPEGPYKCKNSGEDIVLRIVQPVEGSNRNITADNWFTSFPL
IKKLKHNKKLTYTGTVRSNKRELPSEFLPNKNRELHSSIFGFQEDYTLVSYCPKKNKAVLVVSSMHND
DTIDEENHAKKPEMITFYNKTKIGVDLVDQLCEKYNVARNTRRWPMVIFYDLLNISAINAICIYKANNLN
SKISRRQFIENIAWELLKPQIKYRSTITKLPVELRRRARALLGIEEPSISVIPENLPNYVGRCYVCPRNKN
KSTRRFCGQCRKYACKEHMKDICVNCLNE

>AC_ITR_Left
TTAACCCTTCGTTAGGCGCGTGGTCTACAATTGTAGACCAGCTATAAATATTGACAATTTTAGTAT
TTATAAAAAGCTTGTAGTATCTTGTACTTTATAGTAGCTGTTGTTCATCTTTACTACAGTGCAATCA
GTATGACCGTGACCGCTGACGAGCGTTGTGTTACGAAACATATCAACAATATTAGGAGTGGTCTA
CAATCGTAGACCACGCGACTTGCTAAGTTATATTTTTTTATAAATTTGCAAAATTCTTATTTTTTTTT
TAATTAAGTACATTTGTTTTTAATATTTTTAATTTCAACGAAACATAAACAGTAAGTAAAAATATATG
TATACGTAAACATTTTAATGCATTTGGA

>AC_ITR_Right
TTAACCCTTCGTTAGGCGCGAGGTATATTGTTACTTATTCAGGCGCTTGGTCTACACTTGTAGAC
CATTTTAATTTATGACCAATTCAATTAATTTTAATTATTTTTGTTTATATTAATAATGTAAATTACTTG
TATGCATATAAAGAATCGTATGAACATTGAAATATCCAAAATAATCCATGTACATTTGAAGAAAATA
CAGTTAACTTATTTATAGTGAAATTATCATAATTTCTTTCATGACATTTTTATTTTTAGAACTACTTA
CCCAATTACCTTCCAATCAGTCTGAAAACTATAGATATCTTAAATTGACATGTTTACAAAAGACCC

GAAACAAAGTTAAAATTTATTTTTATTTTTTCTTACCAGTTATCGTCGAACAAATATAGATATTATAA
CTGT

>AP_ITR_Left
TTAACCCTTCGTTAGGCGCGTGGTCTACAATTGTAGACCAGCAATAAATATTGACTATTTTAGTAT
TTATAAATAGCTTGTAATACCTTGTACTTTATAGTATCTTTACTACAGTGTAATCAGTATGATCGTG
ACCGTTGACGAGTGTTGTGTAATAAAACATATCAGTAATATTATGAGTGGTCTACAACTGTAGACC
ACGTGACTTACTAAGTTATTTTTATAAATTTTGCAAAATTCTTATTTTTTTTTTTATTAGGTACTTCTG
TTTTTAATTTCAACGAAACATAAACAGTAAGTAAAAATATATGTATACGTAAGCTTTTTAATACATTT
TGAA

>AP_ITR_Right
TTAACCCTTCGTTAGGCGCGAGGTATATTGTTACTTATTCAGGCGCGTGGTCTACGCCTGTAGAC
CATTTTCATTTATGACCAATTCAATTAATTTTAATTATTTTTGTTTATATTAATAATGTAAATTATTTG
TATGCATATAAAGAATAGTATGAGCATTGAAATATCCAAAATAATCCATGAATATTTGAAGAAAATA
CAGTTAACTTGTTTATAGTGAAATTATCATAATTTCTTTCATGACATTTTTATTTTTAGAACTACTTA
CCCAATTACCTTCCAATCAGTCTGAAAACTATAGACATCTTAAATTTACATGTTTACAAAAGTCCC
GAAACAAAATTAAAATGTATTTTTATATTTTCTTACCAGTTATCGTCGAACAAATATAGGTATTATAA
CTGTTTTGGTGC

>piggyBac_HA
MGSSLDDEHILSALLQSDDELVGEDSDSEVSDHVSEDDVQSDTEEAFIDEVHEVQPTSSGSEILDEQ
NVIEQPGSSLASNRILTLPQRTIRGKNKHCWSTSKPTRRSRVSALNIVRSQRGPTRMCRNIYDPLLCF
KLFFTDEIISEIVKWTNAEISLKRRESMTSATFRDTNEDEIYAFFGILVMTAVRKDNHMSTDDLFDRSLS
MVYVSVMSRDRFDFLIRCLRMDDKSIRPTLRENDVFTPVRKIWDLFIHQCIQNYTPGAHLTIDEQLLGF
RGRCPFRVYIPNKPSKYGIKILMMCDSGTKYMINGMPYLGRGTQTNGVPLGEYYVKELSKPVHGSCR
NITCDNWFTSIPLAKNLLQEPYKLTIVGTVRSNKREIPEVLKNSRSRPVGTSMFCFDGPLTLVSYKPKP
AKMVYLLSSCDEDASINESTGKPQMVMYYNQTKGGVDTLDQMCSVMTCSRKTNRWPMALLYGMINI
ACINSFIIYSHNVSSKGEKVQSRKKFMRNLYMGLTSSFMRKRLEAPTLKRYLRDNISNILPKEVPGTSD
DSTEEPVMKKRTYCTYCPSKIRRKASASCKKCKKVICREHNIDMCQSCF

>piggyBac_ITR_Left
TTAACCCTAGAAAGATAGTCTGCGTAAAATTGACGCATGCATTCTTGAAATATTGCTCTCTCTTTC
TAAATAGCGCGAATCCGTCGCTGTGCATTTAGGACATCTCAGTCGCCGCTTGGAGCTCCCGTGA
GGCGTGCTTGTCAATGCGGTAAGTGTCACTGATTTTGAACTATAACGACCGCGTGAGTCAAAATG
ACGCATGATTATCTTTTACGTGACTTTTAAGATTTAACTCATACGATAATTATATTGTTATTTCATGT
TCTACTTACGTGATAACTTATTATATATATATATTTTCTTGTTATAGATATC

>piggyBac_ITR_Right
TTAACCCTAGAAAGATAATCATATTGTGACGTACGTTAAAGATAATCATGCGTAAAATTGACGCAT
GTGTTTTATCGGTCTGTATATCGAGGTTTATTTATTAATTTGAATAGATATTAAGTTTTATTATATTT
ACACTTACATACTAATAATAAATTCAACAAACAATTTATTTATGTTTATTTATTTATTAAAAAAAAACA
AAAACTCAAAATTTCTTCTATAAAGTAACAAA

>CHO_GS
MATSASSHLNKNIKQMYLCLPQGEKVQAMYIWVDGTGEGLRCKTRTLDCEPKCVEELPEWNFDGSS
TFQSEGSNSDMYLSPVAMFRDPFRRDPNKLVFCEVFKYNRKPAETNLRHSCKRIMDMVSNQHPWF
GMEQEYTLMGTDGHPFGWPSNGFPGPQGPYYCGVGADKAYGRDIVEAHYRACLYAGVKITGTNAE
VMPAQWEFQIGPCEGIRMGDHLWVARFILHRVCEDFGVIATFDPKPIPGNWNGAGCHTNFSTKAMR
EENGLKHIEEAIEKLSKRHRYHIRAYDPKGGLDNARRLTGFHETSNINDFSAGVANRSASIRIPRTVGQ
EKKGYFEDRRPSANCDPFAVTEAIVRTCLLNETGDEPFQYKN

>Prov_GS
MSAEHILSLIEEHNVRFIDLRFTDTKGKEQHITIPAHQVDEDFFEEGQMFDGSSFGGWKGINESDMVL
MPDPSTAMIDPFFADITLIIRCDILEPGTMQGYGRDPRSISKRAEDFLRSSGIADVVLFGPEPEFFVFDD
IRFGNSMHSSYYHIDDIEGAWNTGTKYEGGNKGHRPSVKGGYFPVPPVDSSQDLRSAMCTTMEDLG
LVVEAHHHEVATAGQNEIATRFNTMTKKADETQVYKYVVHNVAHAFGKTATFMPKPLVGDNGSGMH
CHMSLSKDGVNLFAGDKYGGLSEMALYYIGGIIKHARALNAFTNPTTNSYKRLVPGFEAPVMLAYSAR
NRSASIRIPVVASTKARRIEVRFPDPAANPYLAFAAQLMAGLDGIINKIHPGDAMDKNLYDLPPEEAKEI
PTVAGSLEEALAELDKNREFLTRGGVFTDDAIDGYIELLRSDIQRVRMTPHPLEFEMYYSV
