## Supplementary material for "Discovery of a Novel Chimeric Transposase-Transposon System for Advanced Genome Engineering": Integration Site Analysis

| Integration Site | Chromosome | Position | Breakpoint Vector | clone no. | Breakpoint Type | Motif | Gene | Context |
|---|---|---|---|---|---|---|---|---|
| chr1-0.1:13519771 | chr1-0.1 | 13519771 | 8970 | E003102 | right | TTAA | LOC118237663 | intron |
| chr1-0.1:13519774 | chr1-0.1 | 13519774 | 1 | E003102 | left | TTAA | LOC118237663 | intron |
| chr1-0.1:136553348 | chr1-0.1 | 136553348 | 8970 | E003102 | right | TTAA | NA | NA |
| chr1-0.1:136553351 | chr1-0.1 | 136553351 | 1 | E003102 | left | TTAA | NA | NA |
| chr1-0.1:152699576 | chr1-0.1 | 152699576 | 1 | E003102 | left | GCTGTTA | NA | NA |
| chr1-0.1:152699580 | chr1-0.1 | 152699580 | 8971 | E003102 | right | TTAA | NA | NA |
| chr1-0.1:161950008 | chr1-0.1 | 161950008 | 1 | E003102 | left | TTAA | LOC100758601 | intron |
| chr1-0.1:161950011 | chr1-0.1 | 161950011 | 8970 | E003102 | right | TTAA | LOC100758601 | intron |
| chr1-0.1:220444322 | chr1-0.1 | 220444322 | 7 | E003102 | left | TACTGAT | NA | NA |
| chr1-0.1:220444329 | chr1-0.1 | 220444329 | 8970 | E003102 | right | TTAA | NA | NA |
| chr1-0.1:247557119 | chr1-0.1 | 247557119 | 1 | E003102 | left | TTAA | Myo9b | intron |
| chr1-0.1:247557122 | chr1-0.1 | 247557122 | 8970 | E003102 | right | TTAA | Myo9b | intron |
| chr1-1.1:127886625 | chr1-1.1 | 127886625 | 8970 | E003102 | right | TTAA | NA | NA |
| chr1-1.1:127886628 | chr1-1.1 | 127886628 | 1 | E003102 | left | TTAA | NA | NA |
| chr1-1.1:165966637 | chr1-1.1 | 165966637 | 8969 | E003102 | right | AATTTAG | NA | NA |
| chr1-1.1:165966639 | chr1-1.1 | 165966639 | 1 | E003102 | left | TTTAGTA | NA | NA |
| chr1-1.1:188690318 | chr1-1.1 | 188690318 | 8970 | E003102 | right | TTAA | Gnl3 | intron |
| chr1-1.1:188690321 | chr1-1.1 | 188690321 | 1 | E003102 | left | TTAA | Gnl3 | intron |
| chr1-1.1:235739501 | chr1-1.1 | 235739501 | 8970 | E003102 | right | TTAA | NA | NA |
| chr1-1.1:235739504 | chr1-1.1 | 235739504 | 1 | E003102 | left | TTAA | NA | NA |
| chr1-1.1:247790276 | chr1-1.1 | 247790276 | 8970 | E003102 | right | ACAGTTA | NA | NA |
| chr1-1.1:247790280 | chr1-1.1 | 247790280 | 1 | E003102 | left | TTAA | NA | NA |
| chr1-1.1:260052567 | chr1-1.1 | 260052567 | 1 | E003102 | left | TTAA | NA | NA |
| chr1-1.1:260052570 | chr1-1.1 | 260052570 | 8970 | E003102 | right | TTAA | NA | NA |
| chr1-1.1:265732720 | chr1-1.1 | 265732720 | 1 | E003102 | left | TTAA | LOC103161398 | intron |
| chr1-1.1:265732723 | chr1-1.1 | 265732723 | 8970 | E003102 | right | TTAA | LOC103161398 | intron |
| chr10.1:24818680 | chr10.1 | 24818680 | 3 | E003102 | left | CCTTTTA | NA | NA |
| chr10.1:24818683 | chr10.1 | 24818683 | 8967 | E003102 | right | TTTATGT | NA | NA |
| chr2.1:32589956 | chr2.1 | 32589956 | 8970 | E003102 | right | TTAA | Foxj3 | intron |
| chr2.1:32589959 | chr2.1 | 32589959 | 1 | E003102 | left | TTAA | Foxj3 | intron |
| chr2.1:34349138 | chr2.1 | 34349138 | 1 | E003102 | left | TTAA | St3gal3 | intron |
| chr2.1:34349141 | chr2.1 | 34349141 | 8970 | E003102 | right | TTAA | St3gal3 | intron |
| chr2.1:68244054 | chr2.1 | 68244054 | 1 | E003102 | left | TTAA | NA | NA |
| chr2.1:121187014 | chr2.1 | 121187014 | 1 | E003102 | left | TTAA | NA | NA |
| chr2.1:121187017 | chr2.1 | 121187017 | 8970 | E003102 | right | TTAA | NA | NA |
| chr2.1:275285247 | chr2.1 | 275285247 | 1 | E003102 | left | TTAA | NA | NA |
| chr2.1:275285250 | chr2.1 | 275285250 | 8970 | E003102 | right | TTAA | NA | NA |
| chr2.1:342681989 | chr2.1 | 342681989 | 1 | E003102 | left | TTAA | NA | NA |
| chr2.1:342681992 | chr2.1 | 342681992 | 8970 | E003102 | right | TTAA | NA | NA |
| chr2.1:343743586 | chr2.1 | 343743586 | 1 | E003102 | left | TTAA | NA | NA |
| chr2.1:343743589 | chr2.1 | 343743589 | 8970 | E003102 | right | TTAA | NA | NA |
| chr2.1:349313311 | chr2.1 | 349313311 | 8968 | E003102 | right | TTATTTA | NA | NA |
| chr2.1:349313314 | chr2.1 | 349313314 | 1 | E003102 | left | TTTATAG | NA | NA |
| chr2.1:351414737 | chr2.1 | 351414737 | 1 | E003102 | left | TTAA | NA | NA |
| chr2.1:351414740 | chr2.1 | 351414740 | 8970 | E003102 | right | TTAA | NA | NA |
| chr2.1:377966691 | chr2.1 | 377966691 | 8971 | E003102 | right | TTAA | NA | NA |
| chr2.1:377966695 | chr2.1 | 377966695 | 1 | E003102 | left | TAACAAA | NA | NA |
| chr2.1:389874513 | chr2.1 | 389874513 | 1 | E003102 | left | TTAA | LOC100760251 | intron |
| chr2.1:389874516 | chr2.1 | 389874516 | 8970 | E003102 | right | TTAA | LOC100760251 | intron |
| chr2.1:393220047 | chr2.1 | 393220047 | 1 | E003102 | left | TTAA | Slc38a2 | intron |
| chr2.1:393220051 | chr2.1 | 393220051 | 8970 | E003102 | right | TAACTGC | Slc38a2 | intron |
| chr2.1:442816754 | chr2.1 | 442816754 | 1 | E003102 | left | TTAA | Scly | intron |
| chr2.1:442816757 | chr2.1 | 442816757 | 8970 | E003102 | right | TTAA | Scly | intron |
| chr3.1:25324603 | chr3.1 | 25324603 | 1 | E003102 | left | TTAA | Pcgf5 | intron |
| chr3.1:25324606 | chr3.1 | 25324606 | 8970 | E003102 | right | TTAA | Pcgf5 | intron |
| chr3.1:43699075 | chr3.1 | 43699075 | 1 | E003102 | left | TTAA | NA | NA |
| chr3.1:102069190 | chr3.1 | 102069190 | 8970 | E003102 | right | TTAA | Thrsp | intron |
| chr3.1:102069193 | chr3.1 | 102069193 | 1 | E003102 | left | TTAA | Thrsp | intron |
| chr3.1:200993967 | chr3.1 | 200993967 | 8970 | E003102 | right | TTAA | Galnt2 | intron |
| chr3.1:200993970 | chr3.1 | 200993970 | 1 | E003102 | left | TTAA | Galnt2 | intron |
| chr3.1:204971518 | chr3.1 | 204971518 | 8970 | E003102 | right | TTAA | LOC103161335 | intron |
| chr3.1:204971521 | chr3.1 | 204971521 | 8967 | E003102 | right | TTAA | LOC103161335 | intron |
| chr3.1:221636261 | chr3.1 | 221636261 | 8966 | E003102 | right | CCACTAG | Frmd4a | intron |
| chr3.1:221636264 | chr3.1 | 221636264 | 5 | E003102 | left | CTAGGGA | Frmd4a | intron |
| chr3.1:268347276 | chr3.1 | 268347276 | 8967 | E003102 | right | TTAA | NA | NA |
| chr4.1:12483318 | chr4.1 | 12483318 | 8970 | E003102 | right | TTAA | NA | NA |
| chr4.1:12483321 | chr4.1 | 12483321 | 1 | E003102 | left | TTAA | NA | NA |
| chr4.1:15006628 | chr4.1 | 15006628 | 1 | E003102 | left | TATGTTA | NA | NA |
| chr4.1:15006632 | chr4.1 | 15006632 | 8971 | E003102 | right | TTAA | NA | NA |
| chr4.1:23646107 | chr4.1 | 23646107 | 5 | E003102 | left | CAACTAA | NA | NA |
| chr4.1:23646110 | chr4.1 | 23646110 | 8969 | E003102 | right | CTAATGA | NA | NA |
| chr4.1:39887230 | chr4.1 | 39887230 | 8966 | E003102 | right | GGTTACT | NA | NA |
| chr4.1:39887237 | chr4.1 | 39887237 | 8 | E003102 | left | AGTGAAT | NA | NA |
| chr4.1:42076719 | chr4.1 | 42076719 | 8971 | E003102 | right | TTAA | Cip2a | intron |
| chr4.1:42076723 | chr4.1 | 42076723 | 1 | E003102 | left | TAACTTG | Cip2a | intron |
| chr4.1:59884258 | chr4.1 | 59884258 | 8970 | E003102 | right | ATTGGTT | Opa1 | intron |
| chr4.1:59884263 | chr4.1 | 59884263 | 1 | E003102 | left | TTAA | Opa1 | intron |
| chr4.1:82416471 | chr4.1 | 82416471 | 1 | E003102 | left | TTAA | NA | NA |

| | | | | | | | | |
|---|---|---|---|---|---|---|---|---|
| chr4.1:82416474 | chr4.1 | 82416474 | 8970 | E003102 | right | TTAA | NA | NA |
| chr4.1:102211551 | chr4.1 | 102211551 | 8970 | E003102 | right | TTAA | Fam216a | intron |
| chr4.1:102211554 | chr4.1 | 102211554 | 1 | E003102 | left | TTAA | Fam216a | intron |
| chr4.1:167473788 | chr4.1 | 167473788 | 1 | E003102 | left | TTAA | Scaper | intron |
| chr4.1:167473792 | chr4.1 | 167473792 | 8970 | E003102 | right | TAACTAT | Scaper | intron |
| chr4.1:209102946 | chr4.1 | 209102946 | 8970 | E003102 | right | TTAA | Stag1 | intron |
| chr4.1:209102949 | chr4.1 | 209102949 | 1 | E003102 | left | TTAA | Stag1 | intron |
| chr4.1:223323754 | chr4.1 | 223323754 | 1 | E003102 | left | TTAA | NA | NA |
| chr4.1:223323757 | chr4.1 | 223323757 | 8970 | E003102 | right | TTAA | NA | NA |
| chr5.1:50694515 | chr5.1 | 50694515 | 1 | E003102 | left | TTAA | Nek7 | intron |
| chr5.1:50694518 | chr5.1 | 50694518 | 8970 | E003102 | right | TTAA | Nek7 | intron |
| chr6.1:5006022 | chr6.1 | 5006022 | 1 | E003102 | left | TTAA | Stx16 | intron |
| chr6.1:5006025 | chr6.1 | 5006025 | 8970 | E003102 | right | TTAA | Stx16 | intron |
| chr6.1:36016810 | chr6.1 | 36016810 | 1 | E003102 | left | GATGTTA | Kif16b | intron |
| chr6.1:36016814 | chr6.1 | 36016814 | 8971 | E003102 | right | TTAA | Kif16b | intron |
| chr6.1:59967196 | chr6.1 | 59967196 | 8970 | E003102 | right | TTAA | NA | NA |
| chr7.1:10565074 | chr7.1 | 10565074 | 8971 | E003102 | right | TTAA | NA | NA |
| chr7.1:10565075 | chr7.1 | 10565075 | 8 | E003102 | left | TAAGAAT | NA | NA |
| chr7.1:35418345 | chr7.1 | 35418345 | 539 | E003102 | unknown | TTAA | NA | NA |
| chr7.1:130557864 | chr7.1 | 130557864 | 1 | E003102 | left | TTAA | Pgs1 | intron |
| chr7.1:130557867 | chr7.1 | 130557867 | 8970 | E003102 | right | TTAA | Pgs1 | intron |
| chr8.1:1752980 | chr8.1 | 1752980 | 8971 | E003102 | right | TTAA | NA | NA |
| chr8.1:1752986 | chr8.1 | 1752986 | 1 | E003102 | left | ACCCATT | NA | NA |
| chr8.1:11503132 | chr8.1 | 11503132 | 8970 | E003102 | right | TTAA | Eps8 | intron |
| chr8.1:11503135 | chr8.1 | 11503135 | 1 | E003102 | left | TTAA | Eps8 | intron |
| chr8.1:13970663 | chr8.1 | 13970663 | 8970 | E003102 | right | TTAA | NA | NA |
| chr8.1:13970666 | chr8.1 | 13970666 | 1 | E003102 | left | TTAA | NA | NA |
| chr9.1:8219030 | chr9.1 | 8219030 | 1 | E003102 | left | TTAA | NA | NA |
| chr9.1:8219033 | chr9.1 | 8219033 | 8970 | E003102 | right | TTAA | NA | NA |
| chr1-0.1:51936736 | chr1-0.1 | 51936736 | 1 | E003104 | left | TTAA | NA | exon |
| chr1-0.1:61829844 | chr1-0.1 | 61829844 | 8970 | E003104 | right | TTAA | Snx27 | intron |
| chr1-0.1:61829847 | chr1-0.1 | 61829847 | 1 | E003104 | left | TTAA | Snx27 | intron |
| chr1-1.1:10398120 | chr1-1.1 | 10398120 | 1 | E003104 | left | TTAA | Tsga10 | intron |
| chr1-1.1:10398123 | chr1-1.1 | 10398123 | 8970 | E003104 | right | TTAA | Tsga10 | intron |
| chr2.1:454399533 | chr2.1 | 454399533 | 8971 | E003104 | right | TTAA | Tmem232 | intron |
| chr2.1:454399538 | chr2.1 | 454399538 | 1 | E003104 | left | AACCATC | Tmem232 | intron |
| chr3.1:171175818 | chr3.1 | 171175818 | 1 | E003104 | left | TTAA | NA | NA |
| chr3.1:171175821 | chr3.1 | 171175821 | 8970 | E003104 | right | TTAA | NA | NA |
| chr6.1:24039135 | chr6.1 | 24039135 | 8970 | E003104 | right | TTAA | LOC118239095 | intron |
| chr6.1:24039138 | chr6.1 | 24039138 | 1 | E003104 | left | TTAA | LOC118239095 | intron |
| chr6.1:129708907 | chr6.1 | 129708907 | 1 | E003104 | left | TTAA | Arhgap15 | intron |
| chr6.1:129708910 | chr6.1 | 129708910 | 8970 | E003104 | right | TTAA | Arhgap15 | intron |
| chrX.1:26972663 | chrX.1 | 26972663 | 1 | E003104 | left | TTAA | NA | NA |
| chrX.1:26972666 | chrX.1 | 26972666 | 8970 | E003104 | right | TTAA | NA | NA |
| chrX.1:63878050 | chrX.1 | 63878050 | 8970 | E003104 | right | CATGTTA | NA | NA |
| chrX.1:63878054 | chrX.1 | 63878054 | 1 | E003104 | left | TTAA | NA | NA |
| chr1-1.1:158064609 | chr1-1.1 | 158064609 | 1 | E003115 | left | TTAA | NA | NA |
| chr1-1.1:158064611 | chr1-1.1 | 158064611 | 8970 | E003115 | right | TTAA | NA | NA |
| chr1-1.1:212036185 | chr1-1.1 | 212036185 | 1 | E003115 | left | TTAA | NA | NA |
| chr1-1.1:212036188 | chr1-1.1 | 212036188 | 8970 | E003115 | right | TTAA | NA | NA |
| chr3.1:145135067 | chr3.1 | 145135067 | 10052 | E003115 | unknown | TTAA | NA | NA |
| chr3.1:145135070 | chr3.1 | 145135070 | 1 | E003115 | left | TTAA | NA | NA |
| chr4.1:173488559 | chr4.1 | 173488559 | 8970 | E003115 | right | GCTGTTA | NA | NA |
| chr4.1:173488563 | chr4.1 | 173488563 | 1 | E003115 | left | TTAA | NA | NA |
| chr5.1:42529167 | chr5.1 | 42529167 | 8970 | E003115 | right | TTAA | NA | NA |
| chr5.1:42529170 | chr5.1 | 42529170 | 1 | E003115 | left | TTAA | NA | NA |
| chr8.1:38708518 | chr8.1 | 38708518 | 8970 | E003115 | right | AAGAAGG | NA | NA |
| chrX.1:27562262 | chrX.1 | 27562262 | 1 | E003115 | left | TTAA | NA | NA |
| chrX.1:27562265 | chrX.1 | 27562265 | 8970 | E003115 | right | TTAA | NA | NA |
| unplaced_scaffold_151.1:9279 | unplaced_scaffold_151.1 | 9279 | 1 | E003115 | left | TTAA | NA | NA |
| unplaced_scaffold_151.1:9282 | unplaced_scaffold_151.1 | 9282 | 8970 | E003115 | right | TTAA | NA | NA |
| chr1-0.1:189873478 | chr1-0.1 | 189873478 | 8970 | E004130 | right | TTAA | NA | NA |
| chr1-0.1:189873481 | chr1-0.1 | 189873481 | 1 | E004130 | left | TTAA | NA | NA |
| chr1-1.1:150606615 | chr1-1.1 | 150606615 | 1 | E004130 | left | TTAA | NA | NA |
| chr1-1.1:150606618 | chr1-1.1 | 150606618 | 8970 | E004130 | right | TTAA | NA | NA |
| chr1-1.1:175018575 | chr1-1.1 | 175018575 | 1 | E004130 | left | TTAA | Thrb | intron |
| chr1-1.1:175018578 | chr1-1.1 | 175018578 | 8970 | E004130 | right | TTAA | Thrb | intron |
| chr2.1:438971848 | chr2.1 | 438971848 | 8970 | E004130 | right | TTAA | Gigyf2 | intron |
| chr2.1:438971851 | chr2.1 | 438971851 | 1 | E004130 | left | TTAA | Gigyf2 | intron |
| chr3.1:22710516 | chr3.1 | 22710516 | 8970 | E004130 | right | TTAA | LOC113830663 | intron |
| chr3.1:22710519 | chr3.1 | 22710519 | 1 | E004130 | left | TTAA | LOC113830663 | intron |
| chr4.1:46367211 | chr4.1 | 46367211 | 1 | E004130 | left | TTAA | Gramd1c | intron |
| chr4.1:46367214 | chr4.1 | 46367214 | 8970 | E004130 | right | TTAA | Gramd1c | intron |
| chr4.1:157678312 | chr4.1 | 157678312 | 8970 | E004130 | right | TTAA | NA | NA |
| chr4.1:157678315 | chr4.1 | 157678315 | 1 | E004130 | left | TTAA | NA | NA |
| chr7.1:12864444 | chr7.1 | 12864444 | 1 | E004130 | left | TTAA | NA | NA |
| chr7.1:12864447 | chr7.1 | 12864447 | 8970 | E004130 | right | TTAA | NA | NA |
| chr7.1:109479256 | chr7.1 | 109479256 | 1 | E004130 | left | TTAA | NA | exon |
| chr7.1:109479259 | chr7.1 | 109479259 | 8970 | E004130 | right | TTAA | NA | exon |

| | | | | | | | | |
|---|---|---|---|---|---|---|---|---|
| chr1-0.1:211328471 | chr1-0.1 | 211328471 | 1 | E004134 | left | TTAA | Foxp2 | intron |
| chr1-0.1:211328474 | chr1-0.1 | 211328474 | 8970 | E004134 | right | TTAA | Foxp2 | intron |
| chr2.1:255441637 | chr2.1 | 255441637 | 1 | E004134 | left | TTAA | Asf1a | intron |
| chr2.1:255441640 | chr2.1 | 255441640 | 8970 | E004134 | right | TTAA | Asf1a | intron |
| chr2.1:393562300 | chr2.1 | 393562300 | 1 | E004134 | left | TTAA | NA | NA |
| chr2.1:393562303 | chr2.1 | 393562303 | 8970 | E004134 | right | TTAA | NA | NA |
| chr3.1:223243876 | chr3.1 | 223243876 | 8951 | E004134 | right | GGGGTGG | NA | NA |
| chr3.1:223243889 | chr3.1 | 223243889 | 1 | E004134 | left | TTAA | NA | NA |
| chr5.1:169215041 | chr5.1 | 169215041 | 1 | E004134 | left | ACTGTTA | Cog5 | intron |
| chr5.1:169215045 | chr5.1 | 169215045 | 8971 | E004134 | right | TTAA | Cog5 | intron |
| chr1-0.1:115639386 | chr1-0.1 | 115639386 | 1 | E006200 | left | TTAA | Ccdc39 | intron |
| chr1-1.1:141298556 | chr1-1.1 | 141298556 | 1 | E006200 | left | TTAA | NA | NA |
| chr1-1.1:141298559 | chr1-1.1 | 141298559 | 8984 | E006200 | right | TTAA | NA | NA |
| chr2.1:398634546 | chr2.1 | 398634546 | 1 | E006200 | left | TTAA | Rarg | intron |
| chr2.1:398634549 | chr2.1 | 398634549 | 8984 | E006200 | right | TTAA | Rarg | intron |
| chr3.1:247024731 | chr3.1 | 247024731 | 1 | E006200 | left | TTAA | NA | NA |
| chr3.1:247024734 | chr3.1 | 247024734 | 8984 | E006200 | right | TTAA | NA | NA |
| chr4.1:30713171 | chr4.1 | 30713171 | 1 | E006200 | left | TTAA | NA | NA |
| chr4.1:30713174 | chr4.1 | 30713174 | 8984 | E006200 | right | TTAA | NA | NA |
| chr5.1:109239504 | chr5.1 | 109239504 | 8984 | E006200 | right | TTAA | NA | NA |
| chr5.1:109239507 | chr5.1 | 109239507 | 1 | E006200 | left | TTAA | NA | NA |
| chr6.1:41561113 | chr6.1 | 41561113 | 1 | E006200 | left | TTAA | Jag1 | intron |
| chr6.1:41561116 | chr6.1 | 41561116 | 8984 | E006200 | right | TTAA | Jag1 | intron |
| chr7.1:114839585 | chr7.1 | 114839585 | 1 | E006200 | left | TTAA | LOC118237813 | intron |
| chr7.1:114839588 | chr7.1 | 114839588 | 8984 | E006200 | right | TTAA | LOC118237813 | intron |
| chr8.1:23833524 | chr8.1 | 23833524 | 1 | E006200 | left | CTTGTTA | Mical3 | intron |
| chrX.1:21425619 | chrX.1 | 21425619 | 8984 | E006200 | right | TTAA | NA | NA |
| chrX.1:21425622 | chrX.1 | 21425622 | 1 | E006200 | left | TTAA | NA | NA |
| chrX.1:103595713 | chrX.1 | 103595713 | 1 | E006200 | left | TTAA | Arhgap6 | intron |
| chrX.1:103595716 | chrX.1 | 103595716 | 8984 | E006200 | right | TTAA | Arhgap6 | intron |
| unplaced_scaffold_2.1:3907197 | unplaced_scaffold_2.1 | 3907197 | 2 | E006200 | left | TTAA | NA | NA |
| unplaced_scaffold_2.1:3907202 | unplaced_scaffold_2.1 | 3907202 | 8984 | E006200 | right | AACCATG | NA | NA |
| chr2.1:398676740 | chr2.1 | 398676740 | 2 | E006204 | left | ATCTATA | NA | NA |
| chr2.1:398676742 | chr2.1 | 398676742 | 8980 | E006204 | right | CTATAAT | NA | NA |
| chr3.1:211283955 | chr3.1 | 211283955 | 8984 | E006204 | right | TTAA | Pdss1 | intron |
| chr3.1:211283958 | chr3.1 | 211283958 | 1 | E006204 | left | TTAA | Pdss1 | intron |
| chr3.1:213645749 | chr3.1 | 213645749 | 8984 | E006204 | right | TTAA | NA | NA |
| chr3.1:213645752 | chr3.1 | 213645752 | 1 | E006204 | left | TTAA | NA | NA |
| chr4.1:27722033 | chr4.1 | 27722033 | 8984 | E006204 | right | TTAA | NA | NA |
| chr4.1:27722036 | chr4.1 | 27722036 | 1 | E006204 | left | TTAA | NA | NA |
| chr5.1:3835870 | chr5.1 | 3835870 | 8984 | E006204 | right | TTAA | NA | NA |
| chr7.1:5129292 | chr7.1 | 5129292 | 8984 | E006204 | right | TTAA | Cebpzos | intron |
| chr7.1:5129295 | chr7.1 | 5129295 | 1 | E006204 | left | TTAA | Cebpzos | intron |
| unplaced_scaffold_44.1:134119 | unplaced_scaffold_44.1 | 134119 | 1 | E006204 | left | TTAA | NA | NA |
| chr1-1.1:69352269 | chr1-1.1 | 69352269 | 1 | E006206 | left | TTAA | Col4a1 | intron |
| chr1-1.1:69352272 | chr1-1.1 | 69352272 | 8984 | E006206 | right | TTAA | Col4a1 | intron |
| chr1-1.1:162147059 | chr1-1.1 | 162147059 | 1 | E006206 | left | TTAA | NA | CDS |
| chr1-1.1:162147062 | chr1-1.1 | 162147062 | 8984 | E006206 | right | TTAA | NA | CDS |
| chr2.1:244430423 | chr2.1 | 244430423 | 8981 | E006206 | right | ATATAAA | Akap7 | intron |
| chr2.1:244430426 | chr2.1 | 244430426 | 3 | E006206 | left | TAAATGT | Akap7 | intron |
| chr3.1:199677166 | chr3.1 | 199677166 | 1 | E006206 | left | TTAA | Ankrd11 | intron |
| chr3.1:199677169 | chr3.1 | 199677169 | 8984 | E006206 | right | TTAA | Ankrd11 | intron |
| chr4.1:125082951 | chr4.1 | 125082951 | 6 | E006206 | left | TTAA | NA | NA |
| chr4.1:125082952 | chr4.1 | 125082952 | 8984 | E006206 | right | TTAA | NA | NA |
| chr4.1:156022667 | chr4.1 | 156022667 | 1 | E006206 | left | TTAA | NA | NA |
| chr4.1:156022670 | chr4.1 | 156022670 | 8984 | E006206 | right | TTAA | NA | NA |
| chr5.1:154933796 | chr5.1 | 154933796 | 1 | E006206 | left | TTAA | NA | NA |
| chr5.1:154933799 | chr5.1 | 154933799 | 8984 | E006206 | right | TTAA | NA | NA |
| chr8.1:2288888 | chr8.1 | 2288888 | 1 | E006206 | left | TTAA | LOC103163695 | intron |
| chr8.1:2288891 | chr8.1 | 2288891 | 8984 | E006206 | right | TTAA | LOC103163695 | intron |
| chr8.1:63371783 | chr8.1 | 63371783 | 8984 | E006206 | right | TTAA | NA | NA |
| chr8.1:63399550 | chr8.1 | 63399550 | 5 | E006206 | left | AGCCTGA | NA | NA |
| chr1-0.1:222134167 | chr1-0.1 | 222134167 | 1 | E007215 | left | TTAA | Ankib1 | intron |
| chr1-0.1:222134170 | chr1-0.1 | 222134170 | 8984 | E007215 | right | TTAA | Ankib1 | intron |
| chr1-0.1:255089739 | chr1-0.1 | 255089739 | 8984 | E007215 | right | TTAA | NA | NA |
| chr1-0.1:255089742 | chr1-0.1 | 255089742 | 1 | E007215 | left | TTAA | NA | NA |
| chr1-1.1:75182846 | chr1-1.1 | 75182846 | 1 | E007215 | left | TTAA | LOC107980006 | intron |
| chr1-1.1:75182849 | chr1-1.1 | 75182849 | 8984 | E007215 | right | TTAA | LOC107980006 | intron |
| chr1-1.1:124925741 | chr1-1.1 | 124925741 | 7982 | E007215 | unknown | CTTTTCC | NA | NA |
| chr1-1.1:124926026 | chr1-1.1 | 124926026 | 5126 | E007215 | unknown | TTAA | NA | NA |
| chr1-1.1:124926527 | chr1-1.1 | 124926527 | 7683 | E007215 | unknown | GGCTGGC | NA | NA |
| chr1-1.1:124927384 | chr1-1.1 | 124927384 | 1 | E007215 | left | TTAA | NA | NA |
| chr10.1:25530645 | chr10.1 | 25530645 | 1 | E007215 | left | TTAA | NA | NA |
| chr10.1:25530648 | chr10.1 | 25530648 | 8984 | E007215 | right | TTAA | NA | NA |
| chr2.1:371147110 | chr2.1 | 371147110 | 1 | E007215 | left | TTAA | NA | NA |
| chr2.1:371147113 | chr2.1 | 371147113 | 8984 | E007215 | right | TTAA | NA | NA |
| chr2.1:439021524 | chr2.1 | 439021524 | 8984 | E007215 | right | TTAA | Gigyf2 | intron |
| chr2.1:439021527 | chr2.1 | 439021527 | 1 | E007215 | left | TTAA | Gigyf2 | intron |
| chr5.1:167385031 | chr5.1 | 167385031 | 1 | E007215 | left | TTAA | NA | NA |

| | | | | | | | | |
|---|---|---|---|---|---|---|---|---|
| chr5.1:167385034 | chr5.1 | 167385034 | 8984 | E007215 | right | TTAA | NA | NA |
| chr6.1:26645927 | chr6.1 | 26645927 | 1 | E007215 | left | TTAA | Asxl1 | intron |
| chr6.1:26645930 | chr6.1 | 26645930 | 8984 | E007215 | right | TTAA | Asxl1 | intron |
| chr1-0.1:267709538 | chr1-0.1 | 267709538 | 1 | E013141 | left | TTAA | Ank3 | intron |
| chr1-0.1:267709541 | chr1-0.1 | 267709541 | 8970 | E013141 | right | TTAA | Ank3 | intron |
| chr4.1:86953565 | chr4.1 | 86953565 | 1 | E013141 | left | TTAA | Stard13 | intron |
| chr4.1:86953568 | chr4.1 | 86953568 | 8970 | E013141 | right | TTAA | Stard13 | intron |
| chr5.1:20359141 | chr5.1 | 20359141 | 8973 | E013141 | right | TTAA | NA | NA |
| chr5.1:20361650 | chr5.1 | 20361650 | 13 | E013141 | left | AGGCCCT | NA | NA |
| chr6.1:65845936 | chr6.1 | 65845936 | 1 | E013141 | left | TTAA | Aven | intron |
| chr6.1:65845939 | chr6.1 | 65845939 | 8970 | E013141 | right | TTAA | Aven | intron |
| chr6.1:137399729 | chr6.1 | 137399729 | 1 | E013141 | left | TTAA | NA | NA |
| chr6.1:137399733 | chr6.1 | 137399733 | 8970 | E013141 | right | TAACAGA | NA | NA |
| chrX.1:25204041 | chrX.1 | 25204041 | 8970 | E013141 | right | TTAA | Tenm1 | intron |
| chrX.1:25204044 | chrX.1 | 25204044 | 1 | E013141 | left | TTAA | Tenm1 | intron |
| chr1-0.1:65851087 | chr1-0.1 | 65851087 | 8970 | E013145 | right | TTAA | NA | NA |
| chr1-0.1:65851090 | chr1-0.1 | 65851090 | 1 | E013145 | left | TTAA | NA | NA |
| chr1-0.1:209441783 | chr1-0.1 | 209441783 | 1 | E013145 | left | TTAA | NA | NA |
| chr1-0.1:209441786 | chr1-0.1 | 209441786 | 8970 | E013145 | right | TTAA | NA | NA |
| chr1-1.1:5336989 | chr1-1.1 | 5336989 | 8970 | E013145 | right | TTAA | NA | NA |
| chr1-1.1:5343063 | chr1-1.1 | 5343063 | 4 | E013145 | left | ACCCATC | Fhl2 | intron |
| chr10.1:13263754 | chr10.1 | 13263754 | 1 | E013145 | left | TTAA | NA | NA |
| chr10.1:13264281 | chr10.1 | 13264281 | 8163 | E013145 | unknown | TATTCAA | NA | NA |
| chr2.1:252157596 | chr2.1 | 252157596 | 8970 | E013145 | right | TTAA | LOC100752741 | intron |
| chr2.1:252157599 | chr2.1 | 252157599 | 1 | E013145 | left | TTAA | LOC100752741 | intron |
| chr2.1:317064399 | chr2.1 | 317064399 | 1 | E013145 | left | CGAAATT | NA | exon |
| chr2.1:317064401 | chr2.1 | 317064401 | 8970 | E013145 | right | TTAA | NA | exon |
| chr3.1:21086107 | chr3.1 | 21086107 | 16 | E013145 | left | GTGTGTT | Sorbs1 | intron |
| chr3.1:21086111 | chr3.1 | 21086111 | 8970 | E013145 | right | GTTGTTA | Sorbs1 | intron |
| chr3.1:113358554 | chr3.1 | 113358554 | 1 | E013145 | left | TTAA | Arnt2 | intron |
| chr3.1:113358558 | chr3.1 | 113358558 | 8970 | E013145 | right | TAACAAG | Arnt2 | intron |
| chr4.1:209099878 | chr4.1 | 209099878 | 1 | E013145 | left | AAAGTTA | Stag1 | intron |
| chr4.1:209099882 | chr4.1 | 209099882 | 8971 | E013145 | right | TTAA | Stag1 | intron |
| chr6.1:58468226 | chr6.1 | 58468226 | 8971 | E013145 | right | CCAGTTA | NA | NA |
| chr6.1:58468231 | chr6.1 | 58468231 | 1 | E013145 | left | TAACAAA | NA | exon |
| chr1-0.1:270365636 | chr1-0.1 | 270365636 | 1 | E013146 | left | TTAA | Reep3 | intron |
| chr1-0.1:270365639 | chr1-0.1 | 270365639 | 8970 | E013146 | right | TTAA | Reep3 | intron |
| chr1-1.1:85926757 | chr1-1.1 | 85926757 | 1 | E013146 | left | TTAA | Sertad2 | intron |
| chr1-1.1:85926760 | chr1-1.1 | 85926760 | 8970 | E013146 | right | TTAA | Sertad2 | intron |
| chr1-1.1:200076356 | chr1-1.1 | 200076356 | 8969 | E013146 | right | ATTTTAG | NA | NA |
| chr1-1.1:200076359 | chr1-1.1 | 200076359 | 5 | E013146 | left | TTAGTAC | NA | NA |
| chr1-1.1:209819671 | chr1-1.1 | 209819671 | 8970 | E013146 | right | TTAA | LOC103159539 | intron |
| chr1-1.1:209819674 | chr1-1.1 | 209819674 | 1 | E013146 | left | TTAA | LOC103159539 | intron |
| chr3.1:246284669 | chr3.1 | 246284669 | 8970 | E013146 | right | TTTGTTA | LOC100770382 | intron |
| chr3.1:246284673 | chr3.1 | 246284673 | 1 | E013146 | left | TTAA | LOC100770382 | intron |
| chr6.1:35021463 | chr6.1 | 35021463 | 1 | E013146 | left | TTAA | Rrbp1 | intron |
| chr6.1:35021466 | chr6.1 | 35021466 | 8970 | E013146 | right | TTAA | Rrbp1 | intron |
| chr7.1:88915785 | chr7.1 | 88915785 | 8970 | E013146 | right | TTAA | NA | NA |
| chr7.1:117615645 | chr7.1 | 117615645 | 1 | E013146 | left | TTAA | NA | NA |
| chr7.1:117615648 | chr7.1 | 117615648 | 8970 | E013146 | right | TTAA | NA | NA |
| chr8.1:29024010 | chr8.1 | 29024010 | 8970 | E013146 | right | TTAA | Efcab12 | intron |
| chr8.1:29024013 | chr8.1 | 29024013 | 1 | E013146 | left | TTAA | Efcab12 | intron |
| chr2.1:39990612 | chr2.1 | 39990612 | 8970 | E014154 | right | TTAA | Faf1 | intron |
| chr2.1:39990615 | chr2.1 | 39990615 | 1 | E014154 | left | TTAA | Faf1 | intron |
| chr2.1:139501242 | chr2.1 | 139501242 | 1 | E014154 | left | TTAA | NA | NA |
| chr2.1:139501245 | chr2.1 | 139501245 | 8970 | E014154 | right | TTAA | NA | NA |
| chr3.1:183422478 | chr3.1 | 183422478 | 8970 | E014154 | right | TTAA | NA | NA |
| chr8.1:69510713 | chr8.1 | 69510713 | 10 | E014154 | left | CAGATTA | Ctnna2 | intron |
| chr8.1:69510714 | chr8.1 | 69510714 | 8970 | E014154 | right | TTAA | Ctnna2 | intron |
| unplaced_scaffold_532.1:10084 | unplaced_scaffold_532.1 | 10084 | 2209 | E014154 | unknown | TTGTATA | NA | NA |
| unplaced_scaffold_532.1:10114 | unplaced_scaffold_532.1 | 10114 | 1 | E014154 | left | TTAA | NA | NA |
| chr1-0.1:108847374 | chr1-0.1 | 108847374 | 8 | E014156 | left | AAAAATG | NA | NA |
| chr1-0.1:108868904 | chr1-0.1 | 108868904 | 8970 | E014156 | right | TTAA | NA | NA |
| chr1-0.1:165759839 | chr1-0.1 | 165759839 | 1 | E014156 | left | TTAA | NA | NA |
| chr1-0.1:165759842 | chr1-0.1 | 165759842 | 8970 | E014156 | right | TTAA | NA | NA |
| chr2.1:108738010 | chr2.1 | 108738010 | 8970 | E014156 | right | TTAA | NA | NA |
| chr2.1:108738013 | chr2.1 | 108738013 | 1 | E014156 | left | TTAA | NA | NA |
| chr2.1:128920035 | chr2.1 | 128920035 | 8970 | E014156 | right | TTAA | LOC100751097 | intron |
| chr2.1:128920038 | chr2.1 | 128920038 | 1 | E014156 | left | TTAA | LOC100751097 | intron |
| chr2.1:290435715 | chr2.1 | 290435715 | 1 | E014156 | left | TTAA | NA | NA |
| chr2.1:290435718 | chr2.1 | 290435718 | 8970 | E014156 | right | TTAA | NA | NA |
| chr2.1:356190440 | chr2.1 | 356190440 | 1 | E014156 | left | TGTGGTT | LOC113833947 | intron |
| chr2.1:356190445 | chr2.1 | 356190445 | 8971 | E014156 | right | TTAA | LOC113833947 | intron |
| chr3.1:27368055 | chr3.1 | 27368055 | 8970 | E014156 | right | AAAGTTA | NA | NA |
| chr3.1:27368059 | chr3.1 | 27368059 | 1 | E014156 | left | TTAA | NA | NA |
| chr3.1:40883016 | chr3.1 | 40883016 | 1 | E014156 | left | TTAA | Anxa1 | intron |
| chr3.1:40883019 | chr3.1 | 40883019 | 8970 | E014156 | right | TTAA | Anxa1 | intron |
| chr3.1:202736632 | chr3.1 | 202736632 | 8970 | E014156 | right | TTAA | Map3k21 | intron |
| chr3.1:202736635 | chr3.1 | 202736635 | 1 | E014156 | left | TTAA | Map3k21 | intron |

| | | | | | | | | |
|---|---|---|---|---|---|---|---|---|
| chr3.1:208821074 | chr3.1 | 208821074 | 8970 | E014156 | right | TTAA | NA | NA |
| chr3.1:208821077 | chr3.1 | 208821077 | 1 | E014156 | left | TTAA | NA | NA |
| chr4.1:30218186 | chr4.1 | 30218186 | 1 | E014156 | left | TTAA | NA | NA |
| chr4.1:30218189 | chr4.1 | 30218189 | 8970 | E014156 | right | TTAA | NA | NA |
| chr4.1:126889131 | chr4.1 | 126889131 | 8971 | E014156 | right | TGTGTTA | Maml2 | intron |
| chr4.1:126889137 | chr4.1 | 126889137 | 1 | E014156 | left | AACCTAG | Maml2 | intron |
| chr4.1:227651977 | chr4.1 | 227651977 | 8970 | E014156 | right | TTAA | NA | NA |
| chr4.1:227651980 | chr4.1 | 227651980 | 1 | E014156 | left | TTAA | NA | NA |
| chr5.1:12042460 | chr5.1 | 12042460 | 1 | E014156 | left | TTAA | NA | NA |
| chr5.1:12042463 | chr5.1 | 12042463 | 8970 | E014156 | right | TTAA | NA | NA |
| chr6.1:9188149 | chr6.1 | 9188149 | 8970 | E014156 | right | TTAA | NA | NA |
| chr6.1:9188152 | chr6.1 | 9188152 | 1 | E014156 | left | TTAA | NA | NA |
| chr6.1:29020248 | chr6.1 | 29020248 | 8970 | E014156 | right | TTAA | NA | NA |
| chr6.1:29020914 | chr6.1 | 29020914 | 10 | E014156 | left | AAACAAA | NA | NA |
| chr6.1:55892861 | chr6.1 | 55892861 | 8970 | E014156 | right | TTAA | NA | CDS |
| chr6.1:55892864 | chr6.1 | 55892864 | 1 | E014156 | left | TTAA | NA | CDS |
| chr6.1:71370369 | chr6.1 | 71370369 | 8970 | E014156 | right | TTAA | Dcdc1 | intron |
| chr6.1:71370372 | chr6.1 | 71370372 | 1 | E014156 | left | TTAA | Dcdc1 | intron |
| chr6.1:136413295 | chr6.1 | 136413295 | 3138 | E014156 | unknown | TCACCCC | NA | NA |
| chr6.1:136413298 | chr6.1 | 136413298 | 131 | E014156 | unknown | CCCCAAT | NA | NA |
| chrX.1:57892175 | chrX.1 | 57892175 | 8970 | E014156 | right | TTAA | Klhl4 | intron |
| chrX.1:57892178 | chrX.1 | 57892178 | 1 | E014156 | left | TTAA | Klhl4 | intron |
| chr1-0.1:94033283 | chr1-0.1 | 94033283 | 8971 | E014157 | right | GTTGTTA | NA | NA |
| chr1-0.1:94033289 | chr1-0.1 | 94033289 | 1 | E014157 | left | AACCTCT | NA | NA |
| chr2.1:90547971 | chr2.1 | 90547971 | 8970 | E014157 | right | TTAA | LOC118238315 | intron |
| chr2.1:90547974 | chr2.1 | 90547974 | 1 | E014157 | left | TTAA | LOC118238315 | intron |
| chr2.1:130153153 | chr2.1 | 130153153 | 8970 | E014157 | right | TTAGTTA | NA | NA |
| chr2.1:130153157 | chr2.1 | 130153157 | 1 | E014157 | left | TTAA | NA | NA |
| chr2.1:251251966 | chr2.1 | 251251966 | 8970 | E014157 | right | TTAA | NA | NA |
| chr2.1:251251969 | chr2.1 | 251251969 | 1 | E014157 | left | TTAA | NA | NA |
| chr4.1:30496808 | chr4.1 | 30496808 | 8970 | E014157 | right | TTAA | NA | NA |
| chr4.1:30496811 | chr4.1 | 30496811 | 1 | E014157 | left | TTAA | NA | NA |
| chr4.1:89727525 | chr4.1 | 89727525 | 1 | E014157 | left | TTAA | NA | NA |
| chr4.1:89727529 | chr4.1 | 89727529 | 8970 | E014157 | right | TAACAAA | NA | NA |
| chr5.1:43679803 | chr5.1 | 43679803 | 2164 | E014157 | unknown | TTAA | NA | NA |
| chr6.1:93177947 | chr6.1 | 93177947 | 1 | E014157 | left | TTAA | Znf804a | intron |
| chr6.1:93177951 | chr6.1 | 93177951 | 8970 | E014157 | right | TAACTGC | Znf804a | intron |
| chr1-0.1:249138897 | chr1-0.1 | 249138897 | 1 | E014158 | left | TTAA | Pbx4 | intron |
| chr1-0.1:249138900 | chr1-0.1 | 249138900 | 638 | E014158 | unknown | TTAA | Pbx4 | intron |
| chr1-1.1:225067144 | chr1-1.1 | 225067144 | 8970 | E014158 | right | TTAA | Piwil2 | intron |
| chr1-1.1:225067147 | chr1-1.1 | 225067147 | 1 | E014158 | left | TTAA | Piwil2 | intron |
| chr10.1:28122831 | chr10.1 | 28122831 | 1 | E014158 | left | TTAA | LOC103159000 | intron |
| chr10.1:28122834 | chr10.1 | 28122834 | 8970 | E014158 | right | TTAA | LOC103159000 | intron |
| chr2.1:87197347 | chr2.1 | 87197347 | 1 | E014158 | left | TTAA | NA | NA |
| chr2.1:87197350 | chr2.1 | 87197350 | 8970 | E014158 | right | TTAA | NA | NA |
| chr2.1:100273216 | chr2.1 | 100273216 | 8970 | E014158 | right | TTAA | Unc13b | intron |
| chr2.1:100273219 | chr2.1 | 100273219 | 1 | E014158 | left | TTAA | Unc13b | intron |
| chr3.1:243781427 | chr3.1 | 243781427 | 8970 | E014158 | right | TTAA | Zscan26 | intron |
| chr3.1:243781430 | chr3.1 | 243781430 | 1 | E014158 | left | TTAA | Zscan26 | intron |
| chr4.1:12100016 | chr4.1 | 12100016 | 8970 | E014158 | right | TTAA | Jam2 | intron |
| chr4.1:12100019 | chr4.1 | 12100019 | 1 | E014158 | left | TTAA | Jam2 | intron |
| chr4.1:209659983 | chr4.1 | 209659983 | 1 | E014158 | left | TTAA | NA | NA |
| chr4.1:209659986 | chr4.1 | 209659986 | 8970 | E014158 | right | TTAA | NA | NA |
| chr5.1:622628 | chr5.1 | 622628 | 1 | E014158 | left | TTAA | NA | exon |
| chr5.1:143562413 | chr5.1 | 143562413 | 8970 | E014158 | right | TTAA | Sec23a | intron |
| chr5.1:175795280 | chr5.1 | 175795280 | 8971 | E014158 | right | TTAA | Esyt2 | intron |
| chr6.1:129139048 | chr6.1 | 129139048 | 8971 | E014158 | right | TTAA | Gtdc1 | intron |
| chr6.1:129139052 | chr6.1 | 129139052 | 1 | E014158 | left | TAACAAC | Gtdc1 | intron |
| chr8.1:48173887 | chr8.1 | 48173887 | 1 | E014158 | left | TTAA | Suclg2 | intron |
| chr8.1:48173890 | chr8.1 | 48173890 | 8970 | E014158 | right | TTAA | Suclg2 | intron |
| chr8.1:90949901 | chr8.1 | 90949901 | 8970 | E014158 | right | TCAGTTA | Osbpl3 | intron |
| chr8.1:90949905 | chr8.1 | 90949905 | 1 | E014158 | left | TTAA | Osbpl3 | intron |
| chr8.1:97727720 | chr8.1 | 97727720 | 8970 | E014158 | right | TTAA | Rheb | intron |
| chr8.1:97727723 | chr8.1 | 97727723 | 1 | E014158 | left | TTAA | Rheb | intron |
| chr1-0.1:208438578 | chr1-0.1 | 208438578 | 8970 | E014159 | right | TTAA | NA | NA |
| chr1-0.1:208438581 | chr1-0.1 | 208438581 | 1 | E014159 | left | TTAA | NA | NA |
| chr1-0.1:237266475 | chr1-0.1 | 237266475 | 8970 | E014159 | right | TTAA | NA | NA |
| chr1-0.1:237266478 | chr1-0.1 | 237266478 | 1 | E014159 | left | TTAA | NA | NA |
| chr1-1.1:200201714 | chr1-1.1 | 200201714 | 1 | E014159 | left | TTAA | Camk2g | intron |
| chr1-1.1:200201717 | chr1-1.1 | 200201717 | 8970 | E014159 | right | TTAA | Camk2g | intron |
| chr1-1.1:255004608 | chr1-1.1 | 255004608 | 8970 | E014159 | right | TTAA | LOC113839263 | intron |
| chr1-1.1:255004611 | chr1-1.1 | 255004611 | 1 | E014159 | left | TTAA | LOC113839263 | intron |
| chr2.1:19688910 | chr2.1 | 19688910 | 8970 | E014159 | right | TTAA | Gpn2 | intron |
| chr2.1:19695278 | chr2.1 | 19695278 | 1706 | E014159 | unknown | AAGAAAA | Gpatch3 | intron |
| chr2.1:46573115 | chr2.1 | 46573115 | 1 | E014159 | left | TTAA | Slc35d1 | intron |
| chr2.1:46573118 | chr2.1 | 46573118 | 8970 | E014159 | right | TTAA | Slc35d1 | intron |
| chr2.1:276645246 | chr2.1 | 276645246 | 1 | E014159 | left | TTAA | NA | NA |
| chr2.1:276645249 | chr2.1 | 276645249 | 8970 | E014159 | right | TTAA | NA | NA |
| chr2.1:336085537 | chr2.1 | 336085537 | 8970 | E014159 | right | TTAA | NA | NA |

| | | | | | | | | |
|---|---|---|---|---|---|---|---|---|
| chr2.1:336085540 | chr2.1 | 336085540 | 1 | E014159 | left | TTAA | NA | NA |
| chr2.1:343858115 | chr2.1 | 343858115 | 1 | E014159 | left | TTAA | Rai14 | intron |
| chr2.1:343858118 | chr2.1 | 343858118 | 8970 | E014159 | right | TTAA | Rai14 | intron |
| chr2.1:428349017 | chr2.1 | 428349017 | 8970 | E014159 | right | TTAA | Atg9a | intron |
| chr2.1:428349020 | chr2.1 | 428349020 | 1 | E014159 | left | TTAA | Atg9a | intron |
| chr2.1:446580060 | chr2.1 | 446580060 | 8970 | E014159 | right | TTAA | St8sia4 | intron |
| chr2.1:446580063 | chr2.1 | 446580063 | 1 | E014159 | left | TTAA | St8sia4 | intron |
| chr3.1:111605865 | chr3.1 | 111605865 | 8970 | E014159 | right | TTAA | Nox4 | intron |
| chr3.1:111605868 | chr3.1 | 111605868 | 1 | E014159 | left | TTAA | Nox4 | intron |
| chr3.1:131130218 | chr3.1 | 131130218 | 1 | E014159 | left | TTAA | Chsy1 | intron |
| chr3.1:131130221 | chr3.1 | 131130221 | 8970 | E014159 | right | TTAA | Chsy1 | intron |
| chr3.1:138741977 | chr3.1 | 138741977 | 8970 | E014159 | right | TTAA | NA | NA |
| chr3.1:138741980 | chr3.1 | 138741980 | 1 | E014159 | left | TTAA | NA | NA |
| chr4.1:51739705 | chr4.1 | 51739705 | 1 | E014159 | left | TTAA | Pla1a | intron |
| chr4.1:51739708 | chr4.1 | 51739708 | 8970 | E014159 | right | TTAA | Pla1a | intron |
| chr4.1:85277401 | chr4.1 | 85277401 | 8970 | E014159 | right | TTAA | Tex26 | intron |
| chr4.1:85277404 | chr4.1 | 85277404 | 1 | E014159 | left | TTAA | Tex26 | intron |
| chr4.1:198457674 | chr4.1 | 198457674 | 1 | E014159 | left | GATGGTT | NA | NA |
| chr4.1:198457679 | chr4.1 | 198457679 | 8971 | E014159 | right | TTAA | NA | NA |
| chr5.1:57199103 | chr5.1 | 57199103 | 1 | E014159 | left | TTAA | NA | NA |
| chr5.1:57199106 | chr5.1 | 57199106 | 8970 | E014159 | right | TTAA | NA | NA |
| chr5.1:132778837 | chr5.1 | 132778837 | 8970 | E014159 | right | TTAA | NA | NA |
| chr5.1:132784941 | chr5.1 | 132784941 | 10726 | E014159 | unknown | TCTCTCT | NA | NA |
| chr7.1:130974853 | chr7.1 | 130974853 | 1 | E014159 | left | TTAA | LOC118239162 | intron |
| chr7.1:130974856 | chr7.1 | 130974856 | 8970 | E014159 | right | TTAA | LOC118239162 | intron |
| chr8.1:86666032 | chr8.1 | 86666032 | 8970 | E014159 | right | TTAA | NA | NA |
| chr8.1:86666035 | chr8.1 | 86666035 | 1 | E014159 | left | TTAA | NA | NA |
| chrX.1:121575897 | chrX.1 | 121575897 | 5 | E014159 | left | AATGGGC | NA | NA |
| chrX.1:121590144 | chrX.1 | 121590144 | 8970 | E014159 | right | TTAA | NA | exon |
| unplaced_scaffold_125.1:29066 | unplaced_scaffold_125.1 | 29066 | 1 | E014159 | left | CTTAGCC | NA | NA |
| chr1-0.1:151775357 | chr1-0.1 | 151775357 | 8970 | E014160 | right | TTAA | NA | NA |
| chr1-0.1:151775360 | chr1-0.1 | 151775360 | 1 | E014160 | left | TTAA | NA | NA |
| chr1-1.1:76674504 | chr1-1.1 | 76674504 | 9475 | E014160 | unknown | CAGCATG | NA | NA |
| chr1-1.1:88743985 | chr1-1.1 | 88743985 | 1017 | E014160 | unknown | GGCTCTC | NA | NA |
| chr1-1.1:259802257 | chr1-1.1 | 259802257 | 8971 | E014160 | right | TTAA | NA | NA |
| chr1-1.1:259802261 | chr1-1.1 | 259802261 | 1 | E014160 | left | TAACACC | NA | NA |
| chr3.1:44475647 | chr3.1 | 44475647 | 8970 | E014160 | right | TTAA | NA | exon |
| chr3.1:44475650 | chr3.1 | 44475650 | 1 | E014160 | left | TTAA | NA | exon |
| chr3.1:59647014 | chr3.1 | 59647014 | 13 | E014160 | left | AGTATTG | NA | NA |
| chr3.1:59647021 | chr3.1 | 59647021 | 8970 | E014160 | right | TTAA | NA | NA |
| chr4.1:102435789 | chr4.1 | 102435789 | 8970 | E014160 | right | GCTGTTA | Ift81 | intron |
| chr4.1:102435793 | chr4.1 | 102435793 | 8966 | E014160 | right | TTAA | Ift81 | intron |
| chr4.1:102818134 | chr4.1 | 102818134 | 1 | E014160 | left | TTAA | NA | NA |
| chr4.1:102818137 | chr4.1 | 102818137 | 8970 | E014160 | right | TTAA | NA | NA |
| chr5.1:8863758 | chr5.1 | 8863758 | 8970 | E014160 | right | TTAA | NA | NA |
| chr5.1:8863761 | chr5.1 | 8863761 | 1 | E014160 | left | TTAA | NA | NA |
| chr6.1:49240220 | chr6.1 | 49240220 | 1 | E014160 | left | TTAA | NA | NA |
| chr6.1:49240223 | chr6.1 | 49240223 | 8970 | E014160 | right | TTAA | NA | NA |
| chr6.1:126701175 | chr6.1 | 126701175 | 1 | E014160 | left | AACCAGA | NA | NA |
| chr8.1:19283157 | chr8.1 | 19283157 | 1 | E014160 | left | TTAA | NA | NA |
| chrX.1:201402 | chrX.1 | 201402 | 8970 | E014160 | right | TTAA | NA | NA |
| chrX.1:201405 | chrX.1 | 201405 | 1 | E014160 | left | TTAA | NA | NA |
| chrX.1:124033799 | chrX.1 | 124033799 | 1 | E014160 | left | TTAA | LOC103160033 | intron |
| chrX.1:124033802 | chrX.1 | 124033802 | 8970 | E014160 | right | TTAA | LOC103160033 | intron |
| chr1-0.1:93308965 | chr1-0.1 | 93308965 | 1 | E014162 | left | TTAA | NA | NA |
| chr1-0.1:93308968 | chr1-0.1 | 93308968 | 8970 | E014162 | right | TTAA | NA | NA |
| chr1-0.1:94033283 | chr1-0.1 | 94033283 | 8971 | E014162 | right | GTTGTTA | NA | NA |
| chr1-0.1:94033289 | chr1-0.1 | 94033289 | 1 | E014162 | left | AACCTCT | NA | NA |
| chr2.1:90547971 | chr2.1 | 90547971 | 8970 | E014162 | right | TTAA | LOC118238315 | intron |
| chr2.1:90547974 | chr2.1 | 90547974 | 1 | E014162 | left | TTAA | LOC118238315 | intron |
| chr2.1:130153153 | chr2.1 | 130153153 | 8970 | E014162 | right | TTAGTTA | NA | NA |
| chr2.1:130153157 | chr2.1 | 130153157 | 1 | E014162 | left | TTAA | NA | NA |
| chr2.1:251251966 | chr2.1 | 251251966 | 8970 | E014162 | right | TTAA | NA | NA |
| chr2.1:251251969 | chr2.1 | 251251969 | 1 | E014162 | left | TTAA | NA | NA |
| chr4.1:30496808 | chr4.1 | 30496808 | 8970 | E014162 | right | TTAA | NA | NA |
| chr4.1:30496811 | chr4.1 | 30496811 | 1 | E014162 | left | TTAA | NA | NA |
| chr4.1:89727525 | chr4.1 | 89727525 | 1 | E014162 | left | TTAA | NA | NA |
| chr4.1:89727529 | chr4.1 | 89727529 | 8970 | E014162 | right | TAACAAA | NA | NA |
| chr5.1:43679800 | chr5.1 | 43679800 | 2163 | E014162 | unknown | TCATTTA | NA | NA |
| chr6.1:93177947 | chr6.1 | 93177947 | 1 | E014162 | left | TTAA | Znf804a | intron |
| chr6.1:93177951 | chr6.1 | 93177951 | 8970 | E014162 | right | TAACTGC | Znf804a | intron |
| chr2.1:39990612 | chr2.1 | 39990612 | 8970 | E014163 | right | TTAA | Faf1 | intron |
| chr2.1:39990615 | chr2.1 | 39990615 | 1 | E014163 | left | TTAA | Faf1 | intron |
| chr2.1:139501242 | chr2.1 | 139501242 | 1 | E014163 | left | TTAA | NA | NA |
| chr2.1:139501245 | chr2.1 | 139501245 | 8970 | E014163 | right | TTAA | NA | NA |
| chr2.1:232796337 | chr2.1 | 232796337 | 1 | E014163 | left | TTAA | NA | NA |
| chr2.1:232796340 | chr2.1 | 232796340 | 8970 | E014163 | right | TTAA | NA | NA |
| chr3.1:183422478 | chr3.1 | 183422478 | 8970 | E014163 | right | TTAA | NA | NA |
| chr8.1:69510713 | chr8.1 | 69510713 | 10 | E014163 | left | CAGATTA | Ctnna2 | intron |

| | | | | | | | | |
|---|---|---|---|---|---|---|---|---|
| chr8.1:69510714 | chr8.1 | 69510714 | 8970 | E014163 | right | TTAA | Ctnna2 | intron |
| unplaced_scaffold_532.1:10084 | unplaced_scaffold_532.1 | 10084 | 2209 | E014163 | unknown | TTGTATA | NA | NA |
| unplaced_scaffold_532.1:10114 | unplaced_scaffold_532.1 | 10114 | 1 | E014163 | left | TTAA | NA | NA |
| chr1-0.1:178306545 | chr1-0.1 | 178306545 | 1 | E014167 | left | TATGTTA | Pclo | intron |
| chr1-0.1:178306549 | chr1-0.1 | 178306549 | 8971 | E014167 | right | TTAA | Pclo | intron |
| chr1-0.1:181667406 | chr1-0.1 | 181667406 | 8970 | E014167 | right | TCTGTTA | NA | NA |
| chr1-0.1:181667410 | chr1-0.1 | 181667410 | 1 | E014167 | left | TTAA | NA | NA |
| chr1-1.1:210747449 | chr1-1.1 | 210747449 | 8970 | E014167 | right | TTAA | Pck2 | intron |
| chr1-1.1:210747452 | chr1-1.1 | 210747452 | 1 | E014167 | left | TTAA | Pck2 | intron |
| chr1-1.1:266526916 | chr1-1.1 | 266526916 | 8970 | E014167 | right | TTAA | NA | NA |
| chr1-1.1:266526919 | chr1-1.1 | 266526919 | 1 | E014167 | left | TTAA | NA | NA |
| chr2.1:47475996 | chr2.1 | 47475996 | 17 | E014167 | left | TCATTCA | NA | NA |
| chr2.1:47476011 | chr2.1 | 47476011 | 8970 | E014167 | right | TTAA | NA | NA |
| chr4.1:38679681 | chr4.1 | 38679681 | 8970 | E014167 | right | TTAA | NA | NA |
| chr4.1:38679684 | chr4.1 | 38679684 | 1 | E014167 | left | TTAA | NA | NA |
| chr4.1:154924458 | chr4.1 | 154924458 | 1 | E014167 | left | TTAA | Oaf | intron |
| chr4.1:154924464 | chr4.1 | 154924464 | 8970 | E014167 | right | ACCCAGG | Oaf | intron |
| chr8.1:46072718 | chr8.1 | 46072718 | 8 | E014167 | left | TCCATAA | Mitf | intron |
| chrX.1:53995674 | chrX.1 | 53995674 | 1 | E014167 | left | TTAA | LOC100754537 | intron |
| chrX.1:53995677 | chrX.1 | 53995677 | 8970 | E014167 | right | TTAA | LOC100754537 | intron |
| chr1-0.1:59926464 | chr1-0.1 | 59926464 | 8970 | E014168 | right | TTAA | LOC118239199 | intron |
| chr1-0.1:59926467 | chr1-0.1 | 59926467 | 1 | E014168 | left | TTAA | LOC118239199 | intron |
| chr1-0.1:91757605 | chr1-0.1 | 91757605 | 1 | E014168 | left | TTAA | Sucnr1 | intron |
| chr1-0.1:91757608 | chr1-0.1 | 91757608 | 8970 | E014168 | right | TTAA | Sucnr1 | intron |
| chr1-0.1:236957174 | chr1-0.1 | 236957174 | 1 | E014168 | left | TTAA | Galntl6 | intron |
| chr1-0.1:236957177 | chr1-0.1 | 236957177 | 8970 | E014168 | right | TTAA | Galntl6 | intron |
| chr1-1.1:2047267 | chr1-1.1 | 2047267 | 8970 | E014168 | right | TTAA | Septin10 | intron |
| chr1-1.1:2047270 | chr1-1.1 | 2047270 | 1 | E014168 | left | TTAA | Septin10 | intron |
| chr1-1.1:25198760 | chr1-1.1 | 25198760 | 8970 | E014168 | right | TTAA | Il17f | intron |
| chr1-1.1:25198763 | chr1-1.1 | 25198763 | 1 | E014168 | left | TTAA | Il17f | intron |
| chr1-1.1:87490759 | chr1-1.1 | 87490759 | 1 | E014168 | left | TTAA | Meis1 | intron |
| chr1-1.1:87490762 | chr1-1.1 | 87490762 | 8970 | E014168 | right | TTAA | Meis1 | intron |
| chr1-1.1:147930084 | chr1-1.1 | 147930084 | 1 | E014168 | left | TTAA | NA | NA |
| chr1-1.1:147930087 | chr1-1.1 | 147930087 | 8970 | E014168 | right | TTAA | NA | NA |
| chr1-1.1:201332269 | chr1-1.1 | 201332269 | 1 | E014168 | left | TTAA | Psmc6 | intron |
| chr1-1.1:201332272 | chr1-1.1 | 201332272 | 8970 | E014168 | right | TTAA | Psmc6 | intron |
| chr1-1.1:218455749 | chr1-1.1 | 218455749 | 1 | E014168 | left | TTAA | NA | NA |
| chr1-1.1:218455752 | chr1-1.1 | 218455752 | 8970 | E014168 | right | TTAA | NA | NA |
| chr1-1.1:257294865 | chr1-1.1 | 257294865 | 1 | E014168 | left | TTAA | NA | NA |
| chr1-1.1:257294868 | chr1-1.1 | 257294868 | 8970 | E014168 | right | TTAA | NA | NA |
| chr1-1.1:262524019 | chr1-1.1 | 262524019 | 1 | E014168 | left | ATATGTC | NA | NA |
| chr1-1.1:262524046 | chr1-1.1 | 262524046 | 5092 | E014168 | unknown | AATTAGT | NA | NA |
| chr1-1.1:262527394 | chr1-1.1 | 262527394 | 8970 | E014168 | right | TTAA | NA | NA |
| chr10.1:27857906 | chr10.1 | 27857906 | 8970 | E014168 | right | TTAA | NA | NA |
| chr10.1:27857909 | chr10.1 | 27857909 | 1 | E014168 | left | TTAA | NA | NA |
| chr2.1:114584100 | chr2.1 | 114584100 | 8970 | E014168 | right | CCTGTTA | NA | NA |
| chr2.1:114584104 | chr2.1 | 114584104 | 1 | E014168 | left | TTAA | NA | NA |
| chr2.1:364735875 | chr2.1 | 364735875 | 8970 | E014168 | right | TTAA | Nsun2 | intron |
| chr2.1:364735878 | chr2.1 | 364735878 | 1 | E014168 | left | TTAA | Nsun2 | intron |
| chr2.1:439337327 | chr2.1 | 439337327 | 8970 | E014168 | right | TTAA | Inpp5d | intron |
| chr2.1:439337330 | chr2.1 | 439337330 | 1 | E014168 | left | TTAA | Inpp5d | intron |
| chr4.1:16678433 | chr4.1 | 16678433 | 1 | E014168 | left | TTAA | NA | NA |
| chr4.1:16678436 | chr4.1 | 16678436 | 8970 | E014168 | right | TTAA | NA | NA |
| chr4.1:23327395 | chr4.1 | 23327395 | 8970 | E014168 | right | TTAA | NA | NA |
| chr4.1:23327398 | chr4.1 | 23327398 | 1 | E014168 | left | TTAA | NA | NA |
| chr4.1:67225979 | chr4.1 | 67225979 | 1 | E014168 | left | TTAA | NA | NA |
| chr4.1:67225982 | chr4.1 | 67225982 | 8970 | E014168 | right | TTAA | NA | NA |
| chr5.1:66895855 | chr5.1 | 66895855 | 8970 | E014168 | right | TTAA | LOC103159433 | intron |
| chr5.1:66895858 | chr5.1 | 66895858 | 1 | E014168 | left | TTAA | LOC103159433 | intron |
| chr5.1:162991450 | chr5.1 | 162991450 | 8970 | E014168 | right | TCTGTTA | NA | CDS |
| chr5.1:162991454 | chr5.1 | 162991454 | 1 | E014168 | left | TTAA | NA | CDS |
| chr6.1:74899454 | chr6.1 | 74899454 | 5 | E014168 | left | AGTGTTT | Slc1a2 | intron |
| chr6.1:74899456 | chr6.1 | 74899456 | 8970 | E014168 | right | TTAA | Slc1a2 | intron |
| chr6.1:101690852 | chr6.1 | 101690852 | 8970 | E014168 | right | TTAA | Wipf1 | intron |
| chr6.1:101690855 | chr6.1 | 101690855 | 1 | E014168 | left | TTAA | Wipf1 | intron |
| chr6.1:144366829 | chr6.1 | 144366829 | 8970 | E014168 | right | TTAA | Eng | intron |
| chr6.1:144366832 | chr6.1 | 144366832 | 1 | E014168 | left | TTAA | Eng | intron |
| chr7.1:19219861 | chr7.1 | 19219861 | 1 | E014168 | left | TTAA | LOC118239176 | intron |
| chr7.1:19219864 | chr7.1 | 19219864 | 8970 | E014168 | right | TTAA | LOC118239176 | intron |
| chr7.1:112229180 | chr7.1 | 112229180 | 1 | E014168 | left | TTAA | NA | NA |
| chr7.1:112229183 | chr7.1 | 112229183 | 8970 | E014168 | right | TTAA | NA | NA |
| chr7.1:119898117 | chr7.1 | 119898117 | 1 | E014168 | left | TTAA | LOC100773415 | NA |
| chr7.1:119898120 | chr7.1 | 119898120 | 8970 | E014168 | right | TTAA | LOC100773415 | NA |
| chr1-0.1:61596691 | chr1-0.1 | 61596691 | 8970 | E015170 | right | TTAA | NA | NA |
| chr1-0.1:61596694 | chr1-0.1 | 61596694 | 1 | E015170 | left | TTAA | NA | NA |
| chr1-0.1:161322444 | chr1-0.1 | 161322444 | 1 | E015170 | left | TTAA | Cradd | intron |
| chr1-1.1:236620970 | chr1-1.1 | 236620970 | 8967 | E015170 | right | TATATAT | NA | NA |
| chr1-1.1:236620988 | chr1-1.1 | 236620988 | 1 | E015170 | left | TTAA | NA | NA |
| chr2.1:516362 | chr2.1 | 516362 | 1 | E015170 | left | TTAA | NA | NA |

| | | | | | | | | |
|---|---|---|---|---|---|---|---|---|
| chr2.1:516365 | chr2.1 | 516365 | 8970 | E015170 | right | TTAA | NA | NA |
| chr2.1:221567558 | chr2.1 | 221567558 | 8970 | E015170 | right | TTAA | Taf4b | intron |
| chr2.1:221567561 | chr2.1 | 221567561 | 1 | E015170 | left | TTAA | Taf4b | intron |
| chr2.1:281351108 | chr2.1 | 281351108 | 8970 | E015170 | right | TTAA | NA | NA |
| chr2.1:281351111 | chr2.1 | 281351111 | 1 | E015170 | left | TTAA | NA | NA |
| chr4.1:44703029 | chr4.1 | 44703029 | 1 | E015170 | left | TTAA | NA | NA |
| chr4.1:44703032 | chr4.1 | 44703032 | 8970 | E015170 | right | TTAA | NA | NA |
| chr7.1:98503752 | chr7.1 | 98503752 | 8970 | E015170 | right | TTAA | NA | NA |
| chr7.1:98503755 | chr7.1 | 98503755 | 1 | E015170 | left | TTAA | NA | NA |
| chr1-0.1:13699120 | chr1-0.1 | 13699120 | 8970 | E016186 | right | CTTGTTA | LOC103158657 | intron |
| chr1-0.1:13699124 | chr1-0.1 | 13699124 | 1 | E016186 | left | TTAA | LOC103158657 | intron |
| chr1-0.1:59928848 | chr1-0.1 | 59928848 | 8970 | E016186 | right | TTAA | LOC118239199 | intron |
| chr1-0.1:59928851 | chr1-0.1 | 59928851 | 1 | E016186 | left | TTAA | LOC118239199 | intron |
| chr1-0.1:87297918 | chr1-0.1 | 87297918 | 8970 | E016186 | right | TTAA | NA | NA |
| chr1-0.1:87297921 | chr1-0.1 | 87297921 | 1 | E016186 | left | TTAA | NA | NA |
| chr1-0.1:238982503 | chr1-0.1 | 238982503 | 8970 | E016186 | right | TTAA | Sh3rf1 | intron |
| chr1-0.1:238982506 | chr1-0.1 | 238982506 | 1 | E016186 | left | TTAA | Sh3rf1 | intron |
| chr2.1:198255337 | chr2.1 | 198255337 | 8970 | E016186 | right | TTAA | NA | NA |
| chr2.1:198255340 | chr2.1 | 198255340 | 1 | E016186 | left | TTAA | NA | NA |
| chr2.1:334757093 | chr2.1 | 334757093 | 1 | E016186 | left | TCAGTTA | Parp8 | intron |
| chr2.1:334757097 | chr2.1 | 334757097 | 8971 | E016186 | right | TTAA | Parp8 | intron |
| chr2.1:453744431 | chr2.1 | 453744431 | 1 | E016186 | left | TTAA | Man2a1 | intron |
| chr2.1:453744434 | chr2.1 | 453744434 | 8970 | E016186 | right | TTAA | Man2a1 | intron |
| chr3.1:89563180 | chr3.1 | 89563180 | 8970 | E016186 | right | TTAA | NA | NA |
| chr3.1:89563183 | chr3.1 | 89563183 | 1 | E016186 | left | TTAA | NA | NA |
| chr3.1:221557380 | chr3.1 | 221557380 | 8970 | E016186 | right | TTAA | Frmd4a | intron |
| chr3.1:221557383 | chr3.1 | 221557383 | 1 | E016186 | left | TTAA | Frmd4a | intron |
| chr4.1:51733290 | chr4.1 | 51733290 | 8970 | E016186 | right | TTAA | Pla1a | intron |
| chr4.1:51733293 | chr4.1 | 51733293 | 1 | E016186 | left | TTAA | Pla1a | intron |
| chr5.1:49021660 | chr5.1 | 49021660 | 1 | E016186 | left | TTAA | NA | NA |
| chr5.1:49021663 | chr5.1 | 49021663 | 8970 | E016186 | right | TTAA | NA | NA |
| chr5.1:127037054 | chr5.1 | 127037054 | 1 | E016186 | left | TTAA | NA | NA |
| chr5.1:127037056 | chr5.1 | 127037056 | 8966 | E016186 | right | AAAGGCC | NA | NA |
| chr7.1:94548479 | chr7.1 | 94548479 | 8969 | E016186 | right | CCATTAG | Asic2 | intron |
| chr8.1:49125187 | chr8.1 | 49125187 | 1 | E016186 | left | TTAA | NA | NA |
| chr8.1:49125896 | chr8.1 | 49125896 | 435 | E016186 | unknown | GCGGAGC | NA | NA |
| chr1-0.1:428177 | chr1-0.1 | 428177 | 8970 | E016188 | right | TTAA | Depdc1 | intron |
| chr1-0.1:428180 | chr1-0.1 | 428180 | 1 | E016188 | left | TTAA | Depdc1 | intron |
| chr1-0.1:72936021 | chr1-0.1 | 72936021 | 1 | E016188 | left | TTAA | NA | NA |
| chr1-0.1:72936024 | chr1-0.1 | 72936024 | 8970 | E016188 | right | TTAA | NA | NA |
| chr1-0.1:86480009 | chr1-0.1 | 86480009 | 8973 | E016188 | right | TTAA | LOC103159251 | intron |
| chr1-0.1:86480013 | chr1-0.1 | 86480013 | 1 | E016188 | left | TAACGAG | LOC103159251 | intron |
| chr1-1.1:45552333 | chr1-1.1 | 45552333 | 1 | E016188 | left | TTAA | NA | NA |
| chr1-1.1:45552336 | chr1-1.1 | 45552336 | 8970 | E016188 | right | TTAA | NA | NA |
| chr1-1.1:69692894 | chr1-1.1 | 69692894 | 8970 | E016188 | right | TTAA | NA | NA |
| chr1-1.1:69692897 | chr1-1.1 | 69692897 | 1 | E016188 | left | TTAA | NA | NA |
| chr1-1.1:217791061 | chr1-1.1 | 217791061 | 8970 | E016188 | right | TTAA | NA | NA |
| chr1-1.1:217791070 | chr1-1.1 | 217791070 | 1 | E016188 | left | CTTCTGT | NA | NA |
| chr2.1:243826451 | chr2.1 | 243826451 | 8970 | E016188 | right | TTAA | NA | NA |
| chr2.1:243826454 | chr2.1 | 243826454 | 1 | E016188 | left | TTAA | NA | NA |
| chr2.1:246798232 | chr2.1 | 246798232 | 8970 | E016188 | right | TTAA | NA | NA |
| chr2.1:246798235 | chr2.1 | 246798235 | 1 | E016188 | left | TTAA | NA | NA |
| chr2.1:340426202 | chr2.1 | 340426202 | 8970 | E016188 | right | TTAA | NA | NA |
| chr2.1:340426205 | chr2.1 | 340426205 | 1 | E016188 | left | TTAA | NA | NA |
| chr3.1:70280144 | chr3.1 | 70280144 | 8970 | E016188 | right | TTAA | NA | NA |
| chr3.1:70280147 | chr3.1 | 70280147 | 1 | E016188 | left | TTAA | NA | NA |
| chr3.1:77733403 | chr3.1 | 77733403 | 1 | E016188 | left | TTAA | Rbbp6 | intron |
| chr3.1:77733406 | chr3.1 | 77733406 | 8970 | E016188 | right | TTAA | Rbbp6 | intron |
| chr3.1:271842549 | chr3.1 | 271842549 | 1 | E016188 | left | TTAA | Auh | intron |
| chr3.1:271842552 | chr3.1 | 271842552 | 8970 | E016188 | right | TTAA | Auh | intron |
| chr3.1:281899965 | chr3.1 | 281899965 | 1 | E016188 | left | TTAA | Prxl2c | intron |
| chr3.1:281899968 | chr3.1 | 281899968 | 8970 | E016188 | right | TTAA | Prxl2c | intron |
| chr4.1:28078428 | chr4.1 | 28078428 | 8970 | E016188 | right | TTAA | Chmp2b | intron |
| chr4.1:28078431 | chr4.1 | 28078431 | 1 | E016188 | left | TTAA | Chmp2b | intron |
| chr6.1:101690115 | chr6.1 | 101690115 | 8967 | E016188 | right | TTAA | Wipf1 | intron |
| chr6.1:101690118 | chr6.1 | 101690118 | 1 | E016188 | left | TTAA | Wipf1 | intron |
| chrX.1:101295991 | chrX.1 | 101295991 | 8970 | E016188 | right | TTAA | NA | NA |
| chrX.1:101295994 | chrX.1 | 101295994 | 1 | E016188 | left | TTAA | NA | NA |
| chr1-0.1:219759257 | chr1-0.1 | 219759257 | 8971 | E016189 | right | TTAA | Ppp1r9a | intron |
| chr1-0.1:219759258 | chr1-0.1 | 219759258 | 3 | E016189 | left | TAATTGC | Ppp1r9a | intron |
| chr1-1.1:140740862 | chr1-1.1 | 140740862 | 1 | E016189 | left | TTAA | Scfd2 | intron |
| chr1-1.1:140740865 | chr1-1.1 | 140740865 | 8970 | E016189 | right | TTAA | Scfd2 | intron |
| chr2.1:314328926 | chr2.1 | 314328926 | 8970 | E016189 | right | TTAA | Iqgap2 | intron |
| chr2.1:314328929 | chr2.1 | 314328929 | 1 | E016189 | left | TTAA | Iqgap2 | intron |
| chr3.1:96419652 | chr3.1 | 96419652 | 1 | E016189 | left | TTAA | NA | NA |
| chr3.1:96419655 | chr3.1 | 96419655 | 8970 | E016189 | right | TTAA | NA | NA |
| chr3.1:196283803 | chr3.1 | 196283803 | 8970 | E016189 | right | TTAA | NA | NA |
| chr3.1:196283806 | chr3.1 | 196283806 | 1 | E016189 | left | TTAA | NA | NA |
| chr3.1:212114786 | chr3.1 | 212114786 | 8971 | E016189 | right | TTAA | Gpr158 | intron |

| | | | | | | | | |
|---|---|---|---|---|---|---|---|---|
| chr3.1:212114789 | chr3.1 | 212114789 | 1 | E016189 | left | TTAA | Gpr158 | intron |
| chr3.1:212412938 | chr3.1 | 212412938 | 597 | E016189 | unknown | GGGCGTT | NA | NA |
| chr3.1:212412948 | chr3.1 | 212412948 | 7562 | E016189 | unknown | TAAGGAA | NA | NA |
| chr3.1:262246229 | chr3.1 | 262246229 | 1 | E016189 | left | TTAA | NA | NA |
| chr3.1:262246232 | chr3.1 | 262246232 | 8970 | E016189 | right | TTAA | NA | NA |
| chr4.1:22715683 | chr4.1 | 22715683 | 1 | E016189 | left | TTAA | NA | NA |
| chr4.1:22715686 | chr4.1 | 22715686 | 8970 | E016189 | right | TTAA | NA | NA |
| chr4.1:53738467 | chr4.1 | 53738467 | 8970 | E016189 | right | TTAA | Dtx3l | intron |
| chr4.1:53738467 | chr4.1 | 53738467 | 1 | E016189 | left | TTAA | Dtx3l | intron |
| chr4.1:109169458 | chr4.1 | 109169458 | 8970 | E016189 | right | TTAA | NA | NA |
| chr4.1:109169461 | chr4.1 | 109169461 | 1 | E016189 | left | TTAA | NA | NA |
| chr6.1:99375047 | chr6.1 | 99375047 | 8970 | E016189 | right | TTAA | Nfe2l2 | intron |
| chr6.1:99375050 | chr6.1 | 99375050 | 1 | E016189 | left | TTAA | Nfe2l2 | intron |
| chr6.1:114195352 | chr6.1 | 114195352 | 1 | E016189 | left | TTAA | LOC118239066 | intron |
| chr6.1:114195355 | chr6.1 | 114195355 | 8970 | E016189 | right | TTAA | LOC118239066 | intron |
| chr6.1:124012692 | chr6.1 | 124012692 | 5 | E016189 | left | ATACTAA | NA | NA |
| chr6.1:124012696 | chr6.1 | 124012696 | 8969 | E016189 | right | TAACAGC | NA | NA |
| chr7.1:36012513 | chr7.1 | 36012513 | 1 | E016189 | left | TCTGTTA | NA | NA |
| chr7.1:36012517 | chr7.1 | 36012517 | 8971 | E016189 | right | TTAA | NA | NA |
| chr8.1:28568486 | chr8.1 | 28568486 | 8970 | E016189 | right | TTAA | Zfand4 | intron |
| chr8.1:45072694 | chr8.1 | 45072694 | 1 | E016189 | left | TTAA | LOC103160634 | intron |
| chr8.1:45072697 | chr8.1 | 45072697 | 8970 | E016189 | right | TTAA | LOC103160634 | intron |
| chr8.1:65423626 | chr8.1 | 65423626 | 1 | E016189 | left | TTAA | NA | NA |
| chr8.1:65423629 | chr8.1 | 65423629 | 8970 | E016189 | right | TTAA | NA | NA |
| chrX.1:105128330 | chrX.1 | 105128330 | 8970 | E016189 | right | TTAA | Wwc3 | intron |
| chrX.1:105128333 | chrX.1 | 105128333 | 1 | E016189 | left | TTAA | Wwc3 | intron |
| chrX.1:121720594 | chrX.1 | 121720594 | 8970 | E016189 | right | TTAA | NA | NA |
| chrX.1:121720597 | chrX.1 | 121720597 | 1 | E016189 | left | TTAA | NA | NA |
| chr1-0.1:73189686 | chr1-0.1 | 73189686 | 1 | E016191 | left | TTAA | Pdgfc | intron |
| chr1-0.1:73189689 | chr1-0.1 | 73189689 | 8970 | E016191 | right | TTAA | Pdgfc | intron |
| chr1-0.1:209506072 | chr1-0.1 | 209506072 | 8970 | E016191 | right | TTAA | NA | NA |
| chr1-0.1:209506075 | chr1-0.1 | 209506075 | 1 | E016191 | left | TTAA | NA | NA |
| chr1-1.1:103814601 | chr1-1.1 | 103814601 | 8970 | E016191 | right | TTAA | Htt | intron |
| chr1-1.1:103814604 | chr1-1.1 | 103814604 | 1 | E016191 | left | TTAA | Htt | intron |
| chr1-1.1:271879422 | chr1-1.1 | 271879422 | 8970 | E016191 | right | TTAA | Ubac2 | intron |
| chr1-1.1:271879425 | chr1-1.1 | 271879425 | 1 | E016191 | left | TTAA | Ubac2 | intron |
| chr10.1:29613024 | chr10.1 | 29613024 | 1 | E016191 | left | ATAGTTA | LOC113837489 | intron |
| chr10.1:29613028 | chr10.1 | 29613028 | 8971 | E016191 | right | TTAA | LOC113837489 | intron |
| chr2.1:81571949 | chr2.1 | 81571949 | 8970 | E016191 | right | TTAA | NA | NA |
| chr2.1:81571952 | chr2.1 | 81571952 | 1 | E016191 | left | TTAA | NA | NA |
| chr2.1:337543292 | chr2.1 | 337543292 | 8971 | E016191 | right | TTAA | LOC113834131 | intron |
| chr3.1:27469712 | chr3.1 | 27469712 | 3 | E016191 | left | CTCCTAA | Fas | intron |
| chr3.1:27469715 | chr3.1 | 27469715 | 8969 | E016191 | right | CTAAGGT | Fas | intron |
| chr3.1:34626894 | chr3.1 | 34626894 | 1 | E016191 | left | TTAA | LOC100770307 | intron |
| chr3.1:34626897 | chr3.1 | 34626897 | 8970 | E016191 | right | TTAA | LOC100770307 | intron |
| chr3.1:43699072 | chr3.1 | 43699072 | 8970 | E016191 | right | TTAA | NA | NA |
| chr3.1:43699075 | chr3.1 | 43699075 | 1 | E016191 | left | TTAA | NA | NA |
| chr3.1:101707677 | chr3.1 | 101707677 | 8971 | E016191 | right | TTAA | Pak1 | intron |
| chr3.1:101707681 | chr3.1 | 101707681 | 1 | E016191 | left | TAACACA | Pak1 | intron |
| chr3.1:228861794 | chr3.1 | 228861794 | 1 | E016191 | left | TTAA | NA | NA |
| chr3.1:228861797 | chr3.1 | 228861797 | 8970 | E016191 | right | TTAA | NA | NA |
| chr3.1:269445104 | chr3.1 | 269445104 | 1 | E016191 | left | TTAA | Cenpp | intron |
| chr3.1:269445107 | chr3.1 | 269445107 | 8970 | E016191 | right | TTAA | Cenpp | intron |
| chr3.1:276893869 | chr3.1 | 276893869 | 1637 | E016191 | unknown | TGTGTTT | Spock1 | intron |
| chr3.1:276901394 | chr3.1 | 276901394 | 8960 | E016191 | right | TAGTGTT | Spock1 | intron |
| chr4.1:116007728 | chr4.1 | 116007728 | 1 | E016191 | left | TTAA | LOC107980100 | intron |
| chr4.1:116007731 | chr4.1 | 116007731 | 8970 | E016191 | right | TTAA | LOC107980100 | intron |
| chr4.1:144592237 | chr4.1 | 144592237 | 8971 | E016191 | right | TTAA | Arhgap32 | intron |
| chr4.1:144592242 | chr4.1 | 144592242 | 1 | E016191 | left | AACCTTT | Arhgap32 | intron |
| chr4.1:192302036 | chr4.1 | 192302036 | 8970 | E016191 | right | TCAGTTA | NA | NA |
| chr4.1:192302040 | chr4.1 | 192302040 | 1 | E016191 | left | TTAA | NA | NA |
| chr4.1:208797885 | chr4.1 | 208797885 | 1 | E016191 | left | TTAA | Il20rb | intron |
| chr4.1:208797888 | chr4.1 | 208797888 | 8970 | E016191 | right | TTAA | Il20rb | intron |
| chr5.1:38587998 | chr5.1 | 38587998 | 8971 | E016191 | right | TTAA | Nmnat2 | intron |
| chr5.1:38588002 | chr5.1 | 38588002 | 1 | E016191 | left | TAACTGA | Nmnat2 | intron |
| chr6.1:58948413 | chr6.1 | 58948413 | 1 | E016191 | left | TTAA | Exd1 | intron |
| chr6.1:58948416 | chr6.1 | 58948416 | 8970 | E016191 | right | TTAA | Exd1 | intron |
| chr7.1:88854464 | chr7.1 | 88854464 | 1 | E016191 | left | TTAA | LOC103158874 | intron |
| chr7.1:88854467 | chr7.1 | 88854467 | 8970 | E016191 | right | TTAA | LOC103158874 | intron |
| chr8.1:74569665 | chr8.1 | 74569665 | 8970 | E016191 | right | TTAA | NA | NA |
| chr8.1:74569668 | chr8.1 | 74569668 | 1 | E016191 | left | TTAA | NA | NA |
| chr1-0.1:187051693 | chr1-0.1 | 187051693 | 8970 | E016192 | right | TTAA | Dbf4 | intron |
| chr1-0.1:187051696 | chr1-0.1 | 187051696 | 1 | E016192 | left | TTAA | Dbf4 | intron |
| chr1-0.1:234472688 | chr1-0.1 | 234472688 | 8971 | E016192 | right | TTAA | Glra3 | intron |
| chr1-0.1:234472692 | chr1-0.1 | 234472692 | 1 | E016192 | left | TAACACA | Glra3 | intron |
| chr1-1.1:8338242 | chr1-1.1 | 8338242 | 8970 | E016192 | right | TTAA | LOC103159864 | intron |
| chr1-1.1:8338245 | chr1-1.1 | 8338245 | 1 | E016192 | left | TTAA | LOC103159864 | intron |
| chr1-1.1:11631356 | chr1-1.1 | 11631356 | 8970 | E016192 | right | TTAA | LOC113832259 | intron |
| chr1-1.1:11631359 | chr1-1.1 | 11631359 | 1 | E016192 | left | TTAA | LOC113832259 | intron |

| | | | | | | | | |
|---|---|---|---|---|---|---|---|---|
| chr1-1.1:31066466 | chr1-1.1 | 31066466 | 1 | E016192 | left | TTAA | NA | NA |
| chr1-1.1:31066469 | chr1-1.1 | 31066469 | 8970 | E016192 | right | TTAA | NA | NA |
| chr1-1.1:206743449 | chr1-1.1 | 206743449 | 1 | E016192 | left | TTAA | NA | NA |
| chr1-1.1:206743452 | chr1-1.1 | 206743452 | 8970 | E016192 | right | TTAA | NA | NA |
| chr1-1.1:217847300 | chr1-1.1 | 217847300 | 8967 | E016192 | right | TTTATAG | NA | NA |
| chr1-1.1:217847302 | chr1-1.1 | 217847302 | 8970 | E016192 | right | TATAGTA | NA | NA |
| chr2.1:14100504 | chr2.1 | 14100504 | 8970 | E016192 | right | TTAA | Capzb | intron |
| chr2.1:14100507 | chr2.1 | 14100507 | 1 | E016192 | left | TTAA | Capzb | intron |
| chr2.1:16759196 | chr2.1 | 16759196 | 8970 | E016192 | right | TTAA | Kdm1a | intron |
| chr2.1:16759199 | chr2.1 | 16759199 | 1 | E016192 | left | TTAA | Kdm1a | intron |
| chr2.1:20381046 | chr2.1 | 20381046 | 1 | E016192 | left | TTAA | NA | NA |
| chr2.1:20381049 | chr2.1 | 20381049 | 8970 | E016192 | right | TTAA | NA | NA |
| chr2.1:57815724 | chr2.1 | 57815724 | 8970 | E016192 | right | TTAA | NA | NA |
| chr2.1:57815727 | chr2.1 | 57815727 | 1 | E016192 | left | TTAA | NA | NA |
| chr2.1:249905491 | chr2.1 | 249905491 | 1 | E016192 | left | TTAA | NA | NA |
| chr2.1:249908666 | chr2.1 | 249908666 | 8965 | E016192 | right | TTGCTTT | NA | NA |
| chr2.1:261376315 | chr2.1 | 261376315 | 1 | E016192 | left | TTAA | NA | NA |
| chr2.1:261376318 | chr2.1 | 261376318 | 8970 | E016192 | right | TTAA | NA | NA |
| chr2.1:340181813 | chr2.1 | 340181813 | 1 | E016192 | left | TTAA | Dab2 | intron |
| chr2.1:340181816 | chr2.1 | 340181816 | 8970 | E016192 | right | TTAA | Dab2 | intron |
| chr2.1:391025418 | chr2.1 | 391025418 | 1 | E016192 | left | TTAA | Pus7l | intron |
| chr2.1:391025421 | chr2.1 | 391025421 | 8970 | E016192 | right | TTAA | Pus7l | intron |
| chr2.1:454984118 | chr2.1 | 454984118 | 1 | E016192 | left | GTTGTTA | NA | NA |
| chr2.1:454984122 | chr2.1 | 454984122 | 8971 | E016192 | right | TTAA | NA | NA |
| chr4.1:48566986 | chr4.1 | 48566986 | 1 | E016192 | left | TTAA | Lsamp | intron |
| chr4.1:48566989 | chr4.1 | 48566989 | 8970 | E016192 | right | TTAA | Lsamp | intron |
| chr7.1:95612503 | chr7.1 | 95612503 | 1 | E016192 | left | ATATATA | Ccl1 | intron |
| chr7.1:95612504 | chr7.1 | 95612504 | 8966 | E016192 | right | TATATAA | Ccl1 | intron |
| chr1-0.1:140835108 | chr1-0.1 | 140835108 | 1 | E017223 | left | TTAA | Rab21 | intron |
| chr1-0.1:140835111 | chr1-0.1 | 140835111 | 8984 | E017223 | right | TTAA | Rab21 | intron |
| chr2.1:234695066 | chr2.1 | 234695066 | 8984 | E017223 | right | TGTGTTA | Nfatc1 | intron |
| chr2.1:234695070 | chr2.1 | 234695070 | 1 | E017223 | left | TTAA | Nfatc1 | intron |
| chr3.1:48041632 | chr3.1 | 48041632 | 2 | E017223 | left | GCATTAT | LOC100768487 | intron |
| chr3.1:48041633 | chr3.1 | 48041633 | 8984 | E017223 | right | CATTATT | LOC100768487 | intron |
| chr4.1:17687476 | chr4.1 | 17687476 | 1 | E017223 | left | TTAA | NA | NA |
| chr4.1:17687479 | chr4.1 | 17687479 | 8984 | E017223 | right | TTAA | NA | NA |
| chr5.1:12321934 | chr5.1 | 12321934 | 8984 | E017223 | right | TTAA | NA | NA |
| chr5.1:12321937 | chr5.1 | 12321937 | 1 | E017223 | left | TTAA | NA | NA |
| chr5.1:116358896 | chr5.1 | 116358896 | 1 | E017223 | left | TTAA | NA | exon |
| chr5.1:116358899 | chr5.1 | 116358899 | 8984 | E017223 | right | TTAA | NA | exon |
| chr5.1:181478819 | chr5.1 | 181478819 | 1 | E017223 | left | TTAA | NA | NA |
| chr6.1:154022906 | chr6.1 | 154022906 | 1 | E017223 | left | TTAA | Flt3lg | intron |
| chr6.1:154022909 | chr6.1 | 154022909 | 8984 | E017223 | right | TTAA | Flt3lg | intron |
| chr8.1:16466597 | chr8.1 | 16466597 | 5704 | E017223 | unknown | TTAA | LOC100773926 | intron |
| chr8.1:16771721 | chr8.1 | 16771721 | 10750 | E017223 | unknown | TCCCTTT | NA | NA |
| chr1-0.1:36951324 | chr1-0.1 | 36951324 | 1 | E017227 | left | TTAA | NA | NA |
| chr1-1.1:32890390 | chr1-1.1 | 32890390 | 1 | E017227 | left | TTAA | NA | NA |
| chr1-1.1:32890394 | chr1-1.1 | 32890394 | 8984 | E017227 | right | TAACTAC | NA | NA |
| chr2.1:182547757 | chr2.1 | 182547757 | 8984 | E017227 | right | TGAGTTA | NA | NA |
| chr2.1:182547761 | chr2.1 | 182547761 | 1 | E017227 | left | TTAA | NA | NA |
| chr3.1:196892971 | chr3.1 | 196892971 | 7 | E017227 | left | ATTTATT | Gse1 | intron |
| chr3.1:196892975 | chr3.1 | 196892975 | 8984 | E017227 | right | TTAA | Gse1 | intron |
| chr4.1:209017778 | chr4.1 | 209017778 | 8984 | E017227 | right | TTAA | Stag1 | intron |
| chr4.1:209017781 | chr4.1 | 209017781 | 1 | E017227 | left | TTAA | Stag1 | intron |
| chr7.1:92077916 | chr7.1 | 92077916 | 8984 | E017227 | right | TTAA | Nlk | intron |
| chr7.1:92077919 | chr7.1 | 92077919 | 1 | E017227 | left | TTAA | Nlk | intron |
| chr8.1:44361663 | chr8.1 | 44361663 | 8984 | E017227 | right | TTAA | NA | NA |
| chr1-0.1:24076866 | chr1-0.1 | 24076866 | 1 | E017228 | left | TTAA | Cisd2 | intron |
| chr1-0.1:48302150 | chr1-0.1 | 48302150 | 8984 | E017228 | right | CACTAAT | NA | NA |
| chr1-0.1:48302152 | chr1-0.1 | 48302152 | 2 | E017228 | left | CTAATCT | NA | NA |
| chr1-0.1:83891723 | chr1-0.1 | 83891723 | 1 | E017228 | left | TTAA | NA | NA |
| chr1-0.1:83891726 | chr1-0.1 | 83891726 | 8984 | E017228 | right | TTAA | NA | NA |
| chr1-0.1:151628218 | chr1-0.1 | 151628218 | 1 | E017228 | left | TTAA | NA | NA |
| chr1-0.1:151628221 | chr1-0.1 | 151628221 | 8984 | E017228 | right | TTAA | NA | NA |
| chr1-0.1:179751642 | chr1-0.1 | 179751642 | 1 | E017228 | left | TTAA | NA | NA |
| chr1-0.1:179751645 | chr1-0.1 | 179751645 | 8984 | E017228 | right | TTAA | NA | NA |
| chr1-0.1:231083110 | chr1-0.1 | 231083110 | 1 | E017228 | left | TTAA | LOC118239533 | NA |
| chr1-0.1:231083114 | chr1-0.1 | 231083114 | 8984 | E017228 | right | TAACTGA | LOC118239533 | NA |
| chr1-1.1:139704777 | chr1-1.1 | 139704777 | 1 | E017228 | left | TTAA | Fryl | intron |
| chr1-1.1:139704780 | chr1-1.1 | 139704780 | 8984 | E017228 | right | TTAA | Fryl | intron |
| chr1-1.1:149089963 | chr1-1.1 | 149089963 | 1 | E017228 | left | TTAA | NA | NA |
| chr1-1.1:149089966 | chr1-1.1 | 149089966 | 8984 | E017228 | right | TTAA | NA | NA |
| chr1-1.1:192176752 | chr1-1.1 | 192176752 | 8984 | E017228 | right | TTAA | Ccdc66 | intron |
| chr1-1.1:192176755 | chr1-1.1 | 192176755 | 1 | E017228 | left | TTAA | Ccdc66 | intron |
| chr1-1.1:229394125 | chr1-1.1 | 229394125 | 8984 | E017228 | right | TTAA | NA | NA |
| chr1-1.1:229394128 | chr1-1.1 | 229394128 | 1 | E017228 | left | TTAA | NA | NA |
| chr1-1.1:271724469 | chr1-1.1 | 271724469 | 1 | E017228 | left | TTAA | NA | NA |
| chr1-1.1:271724472 | chr1-1.1 | 271724472 | 8984 | E017228 | right | TTAA | NA | NA |
| chr2.1:6301897 | chr2.1 | 6301897 | 1 | E017228 | left | AACCTCA | NA | NA |

| | | | | | | | | |
|---|---|---|---|---|---|---|---|---|
| chr2.1:60149710 | chr2.1 | 60149710 | 1 | E017228 | left | TTAA | Mllt3 | intron |
| chr2.1:60149713 | chr2.1 | 60149713 | 8984 | E017228 | right | TTAA | Mllt3 | intron |
| chr2.1:185686658 | chr2.1 | 185686658 | 8984 | E017228 | right | TTAA | NA | NA |
| chr2.1:230808265 | chr2.1 | 230808265 | 1 | E017228 | left | TTAA | NA | NA |
| chr2.1:230808268 | chr2.1 | 230808268 | 8984 | E017228 | right | TTAA | NA | NA |
| chr2.1:253382640 | chr2.1 | 253382640 | 8984 | E017228 | right | TTAA | Tbc1d32 | intron |
| chr2.1:256477570 | chr2.1 | 256477570 | 8984 | E017228 | right | TTAA | LOC100760022 | intron |
| chr2.1:256477573 | chr2.1 | 256477573 | 1 | E017228 | left | TTAA | LOC100760022 | intron |
| chr2.1:263792049 | chr2.1 | 263792049 | 1 | E017228 | left | TTAA | Ak9 | intron |
| chr2.1:263792052 | chr2.1 | 263792052 | 8984 | E017228 | right | TTAA | Ak9 | intron |
| chr2.1:334200785 | chr2.1 | 334200785 | 1 | E017228 | left | TTAA | NA | NA |
| chr2.1:334200788 | chr2.1 | 334200788 | 8984 | E017228 | right | TTAA | NA | NA |
| chr2.1:393261776 | chr2.1 | 393261776 | 1043 | E017228 | unknown | TAGAGTG | NA | NA |
| chr2.1:393265254 | chr2.1 | 393265254 | 8984 | E017228 | right | TTAA | NA | NA |
| chr2.1:402387667 | chr2.1 | 402387667 | 1 | E017228 | left | TTAGTTA | LOC103158689 | intron |
| chr2.1:402387671 | chr2.1 | 402387671 | 8985 | E017228 | right | TTAA | LOC103158689 | intron |
| chr2.1:426025094 | chr2.1 | 426025094 | 8980 | E017228 | right | TTCCTTT | NA | NA |
| chr2.1:426025097 | chr2.1 | 426025097 | 1 | E017228 | left | TTAA | NA | NA |
| chr3.1:17419739 | chr3.1 | 17419739 | 8984 | E017228 | right | TTAA | NA | exon |
| chr3.1:27384006 | chr3.1 | 27384006 | 8984 | E017228 | right | TTAA | NA | NA |
| chr3.1:27384009 | chr3.1 | 27384009 | 1 | E017228 | left | TTAA | NA | NA |
| chr3.1:179942324 | chr3.1 | 179942324 | 1 | E017228 | left | TTAA | NA | NA |
| chr3.1:179942327 | chr3.1 | 179942327 | 8984 | E017228 | right | TTAA | NA | NA |
| chr3.1:181639380 | chr3.1 | 181639380 | 1 | E017228 | left | CACGTTA | Slc9a5 | intron |
| chr3.1:181639384 | chr3.1 | 181639384 | 8986 | E017228 | right | TTAA | Slc9a5 | intron |
| chr4.1:59088787 | chr4.1 | 59088787 | 1 | E017228 | left | TTAA | Atp13a3 | intron |
| chr4.1:93547196 | chr4.1 | 93547196 | 8984 | E017228 | right | TTAA | Coro1c | intron |
| chr4.1:93547199 | chr4.1 | 93547199 | 1 | E017228 | left | TTAA | Coro1c | intron |
| chr4.1:134272229 | chr4.1 | 134272229 | 1 | E017228 | left | TTAA | NA | NA |
| chr4.1:134272232 | chr4.1 | 134272232 | 8984 | E017228 | right | TTAA | NA | NA |
| chr4.1:152761808 | chr4.1 | 152761808 | 1 | E017228 | left | TTAA | LOC103158664 | intron |
| chr4.1:152761811 | chr4.1 | 152761811 | 8984 | E017228 | right | TTAA | LOC103158664 | intron |
| chr4.1:156342033 | chr4.1 | 156342033 | 2 | E017228 | left | TTCGTTA | NA | NA |
| chr4.1:156342039 | chr4.1 | 156342039 | 10172 | E017228 | unknown | ATTTATT | NA | NA |
| chr4.1:162809095 | chr4.1 | 162809095 | 8984 | E017228 | right | TTAA | Layn | intron |
| chr4.1:162809098 | chr4.1 | 162809098 | 1 | E017228 | left | TTAA | Layn | intron |
| chr4.1:167977935 | chr4.1 | 167977935 | 8984 | E017228 | right | TTAA | Hmg20a | intron |
| chr4.1:167977938 | chr4.1 | 167977938 | 1 | E017228 | left | TTAA | Hmg20a | intron |
| chr4.1:201525104 | chr4.1 | 201525104 | 1 | E017228 | left | TTAA | Plod2 | intron |
| chr4.1:201525107 | chr4.1 | 201525107 | 8984 | E017228 | right | TTAA | Plod2 | intron |
| chr4.1:209006482 | chr4.1 | 209006482 | 1 | E017228 | left | TTAA | Stag1 | intron |
| chr4.1:209008862 | chr4.1 | 209008862 | 1 | E017228 | left | TTAA | Stag1 | intron |
| chr4.1:209008865 | chr4.1 | 209008865 | 8984 | E017228 | right | TTAA | Stag1 | intron |
| chr5.1:26239045 | chr5.1 | 26239045 | 1 | E017228 | left | TTAA | Ildr2 | intron |
| chr5.1:26239048 | chr5.1 | 26239048 | 8984 | E017228 | right | TTAA | Ildr2 | intron |
| chr5.1:39790713 | chr5.1 | 39790713 | 1 | E017228 | left | TTAA | Edem3 | intron |
| chr5.1:39790716 | chr5.1 | 39790716 | 8984 | E017228 | right | TTAA | Edem3 | intron |
| chr5.1:168805586 | chr5.1 | 168805586 | 8980 | E017228 | right | TGACTAG | Lamb1 | intron |
| chr5.1:168805588 | chr5.1 | 168805588 | 2 | E017228 | left | ACTAGAA | Lamb1 | intron |
| chr6.1:95499639 | chr6.1 | 95499639 | 8984 | E017228 | right | TTAA | NA | NA |
| chr6.1:95499642 | chr6.1 | 95499642 | 1 | E017228 | left | TTAA | NA | NA |
| chr7.1:106949499 | chr7.1 | 106949499 | 8984 | E017228 | right | TTAA | Spag9 | intron |
| chr7.1:106949502 | chr7.1 | 106949502 | 1 | E017228 | left | TTAA | Spag9 | intron |
| chr8.1:2375892 | chr8.1 | 2375892 | 1 | E017228 | left | TTAA | LOC113830878 | intron |
| chr8.1:2375896 | chr8.1 | 2375896 | 8984 | E017228 | right | TAACACA | LOC113830878 | intron |
| chr8.1:24709382 | chr8.1 | 24709382 | 8985 | E017228 | right | TTAA | NA | NA |
| chr8.1:24709386 | chr8.1 | 24709386 | 1 | E017228 | left | TAACACT | NA | NA |
| chr8.1:76304752 | chr8.1 | 76304752 | 8984 | E017228 | right | TTAA | Gng12 | intron |
| chr8.1:76304755 | chr8.1 | 76304755 | 1 | E017228 | left | TTAA | Gng12 | intron |
| chr9.1:21197536 | chr9.1 | 21197536 | 8984 | E017228 | right | TTAA | NA | NA |
| chr9.1:21197539 | chr9.1 | 21197539 | 1 | E017228 | left | TTAA | NA | NA |
| chrX.1:75920606 | chrX.1 | 75920606 | 1 | E017228 | left | TTAA | NA | NA |
| chrX.1:75920609 | chrX.1 | 75920609 | 8984 | E017228 | right | TTAA | NA | NA |
| chr1-0.1:75025373 | chr1-0.1 | 75025373 | 1 | E017231 | left | TTTGTTA | Rapgef2 | intron |
| chr1-0.1:75025377 | chr1-0.1 | 75025377 | 8985 | E017231 | right | TTAA | Rapgef2 | intron |
| chr1-0.1:142132955 | chr1-0.1 | 142132955 | 2 | E017231 | left | TCATTAG | NA | NA |
| chr1-0.1:142132958 | chr1-0.1 | 142132958 | 8980 | E017231 | right | TTAGAGG | NA | NA |
| chr1-0.1:198211386 | chr1-0.1 | 198211386 | 8984 | E017231 | right | TTAA | NA | NA |
| chr1-0.1:198211387 | chr1-0.1 | 198211387 | 2 | E017231 | left | TTAA | NA | NA |
| chr2.1:48758618 | chr2.1 | 48758618 | 8984 | E017231 | right | TTAA | Cachd1 | intron |
| chr2.1:48758621 | chr2.1 | 48758621 | 1 | E017231 | left | TTAA | Cachd1 | intron |
| chr2.1:459338128 | chr2.1 | 459338128 | 1 | E017231 | left | TTAA | NA | NA |
| chr2.1:459338131 | chr2.1 | 459338131 | 8984 | E017231 | right | TTAA | NA | NA |
| chr3.1:53747234 | chr3.1 | 53747234 | 1 | E017231 | left | TTAA | NA | NA |
| chr3.1:53747237 | chr3.1 | 53747237 | 8984 | E017231 | right | TTAA | NA | NA |
| chr4.1:189515710 | chr4.1 | 189515710 | 1 | E017231 | left | TTAA | LOC100754516 | intron |
| chr1-1.1:2058696 | chr1-1.1 | 2058696 | 1 | E017233 | left | TTAA | Septin10 | intron |
| chr1-1.1:2058699 | chr1-1.1 | 2058699 | 8984 | E017233 | right | TTAA | Septin10 | intron |
| chr1-1.1:33335620 | chr1-1.1 | 33335620 | 8984 | E017233 | right | TTAA | NA | NA |

| | | | | | | | | |
|---|---|---|---|---|---|---|---|---|
| chr1-1.1:33335623 | chr1-1.1 | 33335623 | 1 | E017233 | left | TTAA | NA | NA |
| chr2.1:64692410 | chr2.1 | 64692410 | 8984 | E017233 | right | TTAA | NA | NA |
| chr2.1:356728353 | chr2.1 | 356728353 | 1 | E017233 | left | GAGAGGG | Retreg1 | intron |
| chr2.1:356728360 | chr2.1 | 356728360 | 8985 | E017233 | right | TTAA | Retreg1 | intron |
| chr2.1:455758160 | chr2.1 | 455758160 | 8984 | E017233 | right | TTAA | Ptprm | intron |
| chr2.1:455758163 | chr2.1 | 455758163 | 1 | E017233 | left | TTAA | Ptprm | intron |
| chr3.1:29048615 | chr3.1 | 29048615 | 8984 | E017233 | right | TTAA | LOC113834741 | intron |
| chr3.1:29048618 | chr3.1 | 29048618 | 1 | E017233 | left | TTAA | LOC113834741 | intron |
| chr3.1:271777368 | chr3.1 | 271777368 | 8984 | E017233 | right | TTAA | NA | NA |
| chr3.1:271777371 | chr3.1 | 271777371 | 1 | E017233 | left | TTAA | NA | NA |
| chr4.1:209221053 | chr4.1 | 209221053 | 1 | E017233 | left | TTAA | Stag1 | intron |
| chr4.1:209221056 | chr4.1 | 209221056 | 8984 | E017233 | right | TTAA | Stag1 | intron |
| chr4.1:221779250 | chr4.1 | 221779250 | 8985 | E017233 | right | TTAA | Dync1li1 | intron |
| chr4.1:221779254 | chr4.1 | 221779254 | 1 | E017233 | left | TAACAGT | Dync1li1 | intron |
| chr5.1:28100485 | chr5.1 | 28100485 | 8984 | E017233 | right | TTAA | Slc19a2 | intron |
| chr5.1:28100488 | chr5.1 | 28100488 | 1 | E017233 | left | TTAA | Slc19a2 | intron |
| chr6.1:64096073 | chr6.1 | 64096073 | 8984 | E017233 | right | TTAA | NA | NA |
| chr6.1:64096076 | chr6.1 | 64096076 | 1 | E017233 | left | TTAA | NA | NA |
| chr6.1:145950851 | chr6.1 | 145950851 | 1 | E017233 | left | TTAA | Pbx3 | intron |
| chr6.1:145950854 | chr6.1 | 145950854 | 8984 | E017233 | right | TTAA | Pbx3 | intron |
| chr8.1:45385319 | chr8.1 | 45385319 | 1 | E017233 | left | TTAA | NA | NA |
| chrX.1:25946621 | chrX.1 | 25946621 | 8983 | E017233 | right | CTAATAT | LOC103159158 | intron |
| chrX.1:25948314 | chrX.1 | 25948314 | 1445 | E017233 | unknown | ACAGAAC | LOC103159158 | intron |
| chr2.1:326297039 | chr2.1 | 326297039 | 8984 | E017242 | right | TTAA | Elovl7 | intron |
| chr2.1:326297042 | chr2.1 | 326297042 | 1 | E017242 | left | TTAA | Elovl7 | intron |
| chr2.1:398367643 | chr2.1 | 398367643 | 8984 | E017242 | right | TTAA | LOC100774159 | intron |
| chr2.1:398367646 | chr2.1 | 398367646 | 1 | E017242 | left | TTAA | LOC100774159 | intron |
| chr2.1:402166873 | chr2.1 | 402166873 | 8984 | E017242 | right | TTAA | NA | NA |
| chr2.1:402166876 | chr2.1 | 402166876 | 1 | E017242 | left | TTAA | NA | NA |
| chr3.1:10577161 | chr3.1 | 10577161 | 8984 | E017242 | right | TTAA | NA | NA |
| chr3.1:10577164 | chr3.1 | 10577164 | 1 | E017242 | left | TTAA | NA | NA |
| chr3.1:28770894 | chr3.1 | 28770894 | 8990 | E017242 | right | TTAA | Pten | intron |
| chr3.1:28770897 | chr3.1 | 28770897 | 1 | E017242 | left | TTAA | Pten | intron |
| chr3.1:29354657 | chr3.1 | 29354657 | 8984 | E017242 | right | TTAA | Sgms1 | intron |
| chr3.1:29354660 | chr3.1 | 29354660 | 1 | E017242 | left | TTAA | Sgms1 | intron |
| chr3.1:69471910 | chr3.1 | 69471910 | 8984 | E017242 | right | TTAA | Ikzf5 | intron |
| chr4.1:103441041 | chr4.1 | 103441041 | 8984 | E017242 | right | TTAA | NA | NA |
| chr4.1:103441044 | chr4.1 | 103441044 | 1 | E017242 | left | TTAA | NA | NA |
| chr4.1:129080050 | chr4.1 | 129080050 | 1 | E017242 | left | TTAA | Deup1 | intron |
| chr4.1:129080053 | chr4.1 | 129080053 | 8984 | E017242 | right | TTAA | Deup1 | intron |
| chr5.1:102130411 | chr5.1 | 102130411 | 8984 | E017242 | right | TTAA | NA | NA |
| chr5.1:102130414 | chr5.1 | 102130414 | 1 | E017242 | left | TTAA | NA | NA |
| chr5.1:154601942 | chr5.1 | 154601942 | 8984 | E017242 | right | TTTGTTA | NA | NA |
| chr5.1:154601946 | chr5.1 | 154601946 | 1 | E017242 | left | TTAA | NA | NA |
| chr7.1:26062590 | chr7.1 | 26062590 | 1 | E017242 | left | TTAA | NA | NA |
| chr7.1:26062593 | chr7.1 | 26062593 | 8984 | E017242 | right | TTAA | NA | NA |
| chr1-0.1:34342417 | chr1-0.1 | 34342417 | 1 | E018244 | left | TTAA | NA | NA |
| chr1-0.1:34342420 | chr1-0.1 | 34342420 | 8984 | E018244 | right | TTAA | NA | NA |
| chr1-0.1:246371998 | chr1-0.1 | 246371998 | 1 | E018244 | left | TTAA | NA | NA |
| chr1-0.1:246372001 | chr1-0.1 | 246372001 | 8984 | E018244 | right | TTAA | NA | NA |
| chr1-1.1:36369807 | chr1-1.1 | 36369807 | 1 | E018244 | left | TTAA | NA | NA |
| chr1-1.1:36369810 | chr1-1.1 | 36369810 | 8984 | E018244 | right | TTAA | NA | NA |
| chr1-1.1:50923579 | chr1-1.1 | 50923579 | 8985 | E018244 | right | TTAA | Rbpms | intron |
| chr1-1.1:50923583 | chr1-1.1 | 50923583 | 1 | E018244 | left | TAACAGA | Rbpms | intron |
| chr2.1:87527381 | chr2.1 | 87527381 | 8985 | E018244 | right | ATTGTTA | NA | NA |
| chr2.1:87527386 | chr2.1 | 87527386 | 1 | E018244 | left | TAACTTC | NA | NA |
| chr2.1:213957319 | chr2.1 | 213957319 | 8984 | E018244 | right | TTAA | NA | NA |
| chr2.1:408368040 | chr2.1 | 408368040 | 1 | E018244 | left | TTAA | Stk17b | intron |
| chr2.1:408368044 | chr2.1 | 408368044 | 8984 | E018244 | right | TAACACT | Stk17b | intron |
| chr3.1:148322245 | chr3.1 | 148322245 | 8984 | E018244 | right | TTAA | NA | NA |
| chr4.1:85129243 | chr4.1 | 85129243 | 1 | E018244 | left | TTAA | NA | NA |
| chr4.1:85129246 | chr4.1 | 85129246 | 8984 | E018244 | right | TTAA | NA | NA |
| chr4.1:99012634 | chr4.1 | 99012634 | 8984 | E018244 | right | TTAA | LOC118238653 | intron |
| chr4.1:99012637 | chr4.1 | 99012637 | 1 | E018244 | left | TTAA | LOC118238653 | intron |
| chr5.1:45998007 | chr5.1 | 45998007 | 8984 | E018244 | right | TTAA | NA | NA |
| chr5.1:45998010 | chr5.1 | 45998010 | 2 | E018244 | left | TTAA | NA | NA |
| chr7.1:109013539 | chr7.1 | 109013539 | 8986 | E018244 | right | TTAA | NA | NA |
| chr7.1:109013543 | chr7.1 | 109013543 | 1 | E018244 | left | TAACGGA | NA | NA |
| chr8.1:56003050 | chr8.1 | 56003050 | 8984 | E018244 | right | TTAA | Antxr1 | intron |
| chr8.1:56003053 | chr8.1 | 56003053 | 1 | E018244 | left | TTAA | Antxr1 | intron |
| chr1-0.1:226176806 | chr1-0.1 | 226176806 | 8984 | E018245 | right | TTAA | NA | NA |
| chr1-0.1:226176809 | chr1-0.1 | 226176809 | 12 | E018245 | left | AACAATC | NA | NA |
| chr1-1.1:36667596 | chr1-1.1 | 36667596 | 8984 | E018245 | right | TTAA | NA | NA |
| chr1-1.1:36667599 | chr1-1.1 | 36667599 | 1 | E018245 | left | TTAA | NA | NA |
| chr2.1:312016890 | chr2.1 | 312016890 | 1 | E018245 | left | TTAA | NA | NA |
| chr2.1:361728955 | chr2.1 | 361728955 | 8984 | E018245 | right | TTAA | Atpsckmt | intron |
| chr2.1:361728961 | chr2.1 | 361728961 | 5 | E018245 | left | CCCGGCG | Atpsckmt | intron |
| chr4.1:41242870 | chr4.1 | 41242870 | 8984 | E018245 | right | TTAA | NA | NA |
| chr4.1:41242873 | chr4.1 | 41242873 | 1 | E018245 | left | TTAA | NA | NA |

| | | | | | | | | |
|---|---|---|---|---|---|---|---|---|
| chr5.1:34946952 | chr5.1 | 34946952 | 8985 | E018245 | right | TTAA | NA | NA |
| chr5.1:34946957 | chr5.1 | 34946957 | 1 | E018245 | left | AACCACT | NA | NA |
| chr7.1:8670352 | chr7.1 | 8670352 | 8984 | E018245 | right | TTAA | Ltbp1 | intron |
| chr7.1:8670355 | chr7.1 | 8670355 | 1 | E018245 | left | TTAA | Ltbp1 | intron |
| chr8.1:24814837 | chr8.1 | 24814837 | 8984 | E018245 | right | TTAA | NA | NA |
| chr8.1:24814840 | chr8.1 | 24814840 | 1 | E018245 | left | TTAA | NA | NA |
| chr8.1:64490655 | chr8.1 | 64490655 | 1 | E018245 | left | TTAGGTT | Tcf7l1 | intron |
| chr8.1:64490660 | chr8.1 | 64490660 | 8985 | E018245 | right | TTAA | Tcf7l1 | intron |
| chr8.1:64492397 | chr8.1 | 64492397 | 8984 | E018245 | right | TTAA | Tcf7l1 | intron |
| chr8.1:64492400 | chr8.1 | 64492400 | 1 | E018245 | left | TTAA | Tcf7l1 | intron |
| chr1-0.1:88369768 | chr1-0.1 | 88369768 | 1 | E018246 | left | TTAA | NA | exon |
| chr1-0.1:88369771 | chr1-0.1 | 88369771 | 8984 | E018246 | right | TTAA | NA | exon |
| chr1-0.1:119670285 | chr1-0.1 | 119670285 | 8984 | E018246 | right | TTAA | Prkci | intron |
| chr1-0.1:119670288 | chr1-0.1 | 119670288 | 1 | E018246 | left | TTAA | Prkci | intron |
| chr1-1.1:257295748 | chr1-1.1 | 257295748 | 1 | E018246 | left | TTAA | NA | NA |
| chr1-1.1:257295751 | chr1-1.1 | 257295751 | 8990 | E018246 | right | TTAA | NA | NA |
| chr2.1:90747944 | chr2.1 | 90747944 | 1 | E018246 | left | TTAA | NA | exon |
| chr2.1:90747947 | chr2.1 | 90747947 | 8984 | E018246 | right | TTAA | NA | exon |
| chr3.1:55304499 | chr3.1 | 55304499 | 1 | E018246 | left | TTAA | NA | NA |
| chr3.1:237168402 | chr3.1 | 237168402 | 1 | E018246 | left | TTAA | NA | NA |
| chr3.1:237168405 | chr3.1 | 237168405 | 8984 | E018246 | right | TTAA | NA | NA |
| chr5.1:147453537 | chr5.1 | 147453537 | 8984 | E018246 | right | TTAA | NA | NA |
| chr5.1:147453540 | chr5.1 | 147453540 | 1 | E018246 | left | TTAA | NA | NA |
| chr6.1:57471233 | chr6.1 | 57471233 | 8984 | E018246 | right | GGCGTTA | Ubr1 | intron |
| chr6.1:57471237 | chr6.1 | 57471237 | 1 | E018246 | left | TTAA | Ubr1 | intron |
| chr7.1:43246420 | chr7.1 | 43246420 | 8985 | E018246 | right | TTAA | Pkmyt1 | intron |
| chr7.1:43246425 | chr7.1 | 43246425 | 1 | E018246 | left | AACCAGC | Pkmyt1 | intron |
| chr7.1:45593400 | chr7.1 | 45593400 | 1 | E018246 | left | TTAA | Capn15 | intron |
| chr7.1:45593403 | chr7.1 | 45593403 | 8984 | E018246 | right | TTAA | Capn15 | intron |
| chr1-0.1:27564297 | chr1-0.1 | 27564297 | 1 | E018247 | left | TTAA | LOC118238018 | intron |
| chr1-0.1:27564301 | chr1-0.1 | 27564301 | 8984 | E018247 | right | TAACTGA | LOC118238018 | intron |
| chr1-0.1:40437242 | chr1-0.1 | 40437242 | 8984 | E018247 | right | TACTGTT | Snx7 | intron |
| chr1-0.1:40437246 | chr1-0.1 | 40437246 | 1 | E018247 | left | GTTGTTA | Snx7 | intron |
| chr1-0.1:235387091 | chr1-0.1 | 235387091 | 1 | E018247 | left | TTAA | NA | NA |
| chr1-0.1:235387094 | chr1-0.1 | 235387094 | 8984 | E018247 | right | TTAA | NA | NA |
| chr1-0.1:244990375 | chr1-0.1 | 244990375 | 1 | E018247 | left | TTAA | NA | exon |
| chr1-0.1:244990378 | chr1-0.1 | 244990378 | 8984 | E018247 | right | TTAA | NA | exon |
| chr1-1.1:190861050 | chr1-1.1 | 190861050 | 1153 | E018247 | unknown | TTAA | NA | NA |
| chr1-1.1:190861053 | chr1-1.1 | 190861053 | 8984 | E018247 | right | TTAA | NA | NA |
| chr10.1:28248657 | chr10.1 | 28248657 | 1 | E018247 | left | TTAA | LOC103159000 | intron |
| chr10.1:28248660 | chr10.1 | 28248660 | 8984 | E018247 | right | TTAA | LOC103159000 | intron |
| chr2.1:20640275 | chr2.1 | 20640275 | 2 | E018247 | left | CTCTAGA | NA | NA |
| chr2.1:20640277 | chr2.1 | 20640277 | 8980 | E018247 | right | CTAGAAT | NA | NA |
| chr2.1:34311037 | chr2.1 | 34311037 | 8984 | E018247 | right | TTAA | St3gal3 | intron |
| chr2.1:34311040 | chr2.1 | 34311040 | 1 | E018247 | left | TTAA | St3gal3 | intron |
| chr2.1:48418650 | chr2.1 | 48418650 | 1 | E018247 | left | TTAA | Jak1 | intron |
| chr2.1:48418653 | chr2.1 | 48418653 | 8984 | E018247 | right | TTAA | Jak1 | intron |
| chr2.1:140588974 | chr2.1 | 140588974 | 6 | E018247 | left | TCCTCAC | NA | NA |
| chr2.1:140588979 | chr2.1 | 140588979 | 8985 | E018247 | right | TTAA | NA | NA |
| chr2.1:154138195 | chr2.1 | 154138195 | 8984 | E018247 | right | TTAA | Sdcbp | intron |
| chr2.1:154138198 | chr2.1 | 154138198 | 1 | E018247 | left | TTAA | Sdcbp | intron |
| chr2.1:284279632 | chr2.1 | 284279632 | 8984 | E018247 | right | TTAA | Cnksr3 | intron |
| chr2.1:284279635 | chr2.1 | 284279635 | 1 | E018247 | left | TTAA | Cnksr3 | intron |
| chr2.1:398776959 | chr2.1 | 398776959 | 1 | E018247 | left | TTAA | NA | NA |
| chr2.1:398776962 | chr2.1 | 398776962 | 8984 | E018247 | right | TTAA | NA | NA |
| chr3.1:166847779 | chr3.1 | 166847779 | 8984 | E018247 | right | TTAA | NA | NA |
| chr3.1:166847782 | chr3.1 | 166847782 | 1 | E018247 | left | TTAA | NA | NA |
| chr4.1:81566619 | chr4.1 | 81566619 | 8984 | E018247 | right | CATGTTA | NA | CDS |
| chr4.1:81566623 | chr4.1 | 81566623 | 1 | E018247 | left | TTAA | NA | CDS |
| chr4.1:175143539 | chr4.1 | 175143539 | 8984 | E018247 | right | TTAA | Aagab | intron |
| chr4.1:175143542 | chr4.1 | 175143542 | 1 | E018247 | left | TTAA | Aagab | intron |
| chr5.1:52590412 | chr5.1 | 52590412 | 1 | E018247 | left | TTAA | NA | NA |
| chr5.1:52590415 | chr5.1 | 52590415 | 8984 | E018247 | right | TTAA | NA | NA |
| chr6.1:76063191 | chr6.1 | 76063191 | 8984 | E018247 | right | TTAA | Iftap | intron |
| chr6.1:76063199 | chr6.1 | 76063199 | 4 | E018247 | left | ACTGTGG | Iftap | intron |
| chr6.1:95365528 | chr6.1 | 95365528 | 1 | E018247 | left | TTAA | Ppp1r1c | intron |
| chr6.1:95365531 | chr6.1 | 95365531 | 8984 | E018247 | right | TTAA | Ppp1r1c | intron |
| chr2.1:248605530 | chr2.1 | 248605530 | 1 | E018249 | left | TTAA | Cenpw | intron |
| chr2.1:248605533 | chr2.1 | 248605533 | 8984 | E018249 | right | TTAA | Cenpw | intron |
| chr2.1:411290137 | chr2.1 | 411290137 | 1 | E018249 | left | TTAA | Satb2 | intron |
| chr2.1:411290140 | chr2.1 | 411290140 | 8984 | E018249 | right | TTAA | Satb2 | intron |
| chr3.1:32400651 | chr3.1 | 32400651 | 5 | E018249 | left | CCCTATT | NA | NA |
| chr3.1:32400653 | chr3.1 | 32400653 | 8984 | E018249 | right | TTAA | NA | NA |
| chr3.1:57628660 | chr3.1 | 57628660 | 4 | E018249 | left | TTAA | Osbpl5 | intron |
| chr3.1:57628661 | chr3.1 | 57628661 | 8984 | E018249 | right | TTAA | Osbpl5 | intron |
| chr3.1:103402042 | chr3.1 | 103402042 | 8984 | E018249 | right | TTAA | NA | NA |
| chr3.1:103402045 | chr3.1 | 103402045 | 1 | E018249 | left | TTAA | NA | NA |
| chr3.1:244800229 | chr3.1 | 244800229 | 8984 | E018249 | right | TTAA | NA | NA |
| chr3.1:244800232 | chr3.1 | 244800232 | 1 | E018249 | left | TTAA | NA | NA |

| | | | | | | | | |
|---|---|---|---|---|---|---|---|---|
| chr4.1:109223356 | chr4.1 | 109223356 | 8984 | E018249 | right | TTAA | Vkorc1l1 | intron |
| chr4.1:109223359 | chr4.1 | 109223359 | 1 | E018249 | left | TTAA | Vkorc1l1 | intron |
| chr6.1:68061425 | chr6.1 | 68061425 | 1 | E018249 | left | TTAA | Lgr4 | intron |
| chr6.1:68061428 | chr6.1 | 68061428 | 8984 | E018249 | right | TTAA | Lgr4 | intron |
| chr7.1:4761376 | chr7.1 | 4761376 | 8984 | E018249 | right | TTAA | NA | NA |
| chr7.1:4761379 | chr7.1 | 4761379 | 1 | E018249 | left | TTAA | NA | NA |
| chr8.1:57015004 | chr8.1 | 57015004 | 8984 | E018249 | right | TCTGTTA | NA | NA |
| chr8.1:57015008 | chr8.1 | 57015008 | 1 | E018249 | left | TTAA | NA | NA |
| chr8.1:75538440 | chr8.1 | 75538440 | 1 | E018249 | left | TTAA | NA | exon |
| chr1-0.1:101389944 | chr1-0.1 | 101389944 | 8985 | E018250 | right | TTAA | NA | NA |
| chr1-0.1:101389948 | chr1-0.1 | 101389948 | 1 | E018250 | left | TAACTGC | NA | NA |
| chr1-1.1:10915607 | chr1-1.1 | 10915607 | 8984 | E018250 | right | TTAA | Inpp4a | intron |
| chr1-1.1:10915610 | chr1-1.1 | 10915610 | 1 | E018250 | left | TTAA | Inpp4a | intron |
| chr1-1.1:41634166 | chr1-1.1 | 41634166 | 8984 | E018250 | right | TTAA | NA | NA |
| chr1-1.1:41634169 | chr1-1.1 | 41634169 | 1 | E018250 | left | TTAA | NA | NA |
| chr1-1.1:270228220 | chr1-1.1 | 270228220 | 8984 | E018250 | right | TTAA | NA | NA |
| chr1-1.1:270228223 | chr1-1.1 | 270228223 | 1 | E018250 | left | TTAA | NA | NA |
| chr2.1:17160353 | chr2.1 | 17160353 | 2 | E018250 | left | TTATTAG | NA | NA |
| chr2.1:17160355 | chr2.1 | 17160355 | 8984 | E018250 | right | ATTAGTG | NA | NA |
| chr2.1:322781613 | chr2.1 | 322781613 | 8984 | E018250 | right | TTAA | Adamts6 | intron |
| chr2.1:322781616 | chr2.1 | 322781616 | 1 | E018250 | left | TTAA | Adamts6 | intron |
| chr2.1:337459368 | chr2.1 | 337459368 | 1 | E018250 | left | TTAA | NA | NA |
| chr2.1:337459371 | chr2.1 | 337459371 | 8984 | E018250 | right | TTAA | NA | NA |
| chr3.1:78606259 | chr3.1 | 78606259 | 1 | E018250 | left | TTAA | Ubfd1 | intron |
| chr3.1:78606262 | chr3.1 | 78606262 | 8984 | E018250 | right | TTAA | Ubfd1 | intron |
| chr3.1:187345758 | chr3.1 | 187345758 | 8984 | E018250 | right | TTAA | St3gal2 | intron |
| chr3.1:187345761 | chr3.1 | 187345761 | 1 | E018250 | left | TTAA | St3gal2 | intron |
| chr5.1:113674547 | chr5.1 | 113674547 | 1 | E018250 | left | TTAA | Cep128 | intron |
| chr5.1:113674550 | chr5.1 | 113674550 | 8984 | E018250 | right | TTAA | Cep128 | intron |
| chr5.1:150961794 | chr5.1 | 150961794 | 1 | E018250 | left | TTAA | G2e3 | intron |
| chr5.1:150961797 | chr5.1 | 150961797 | 8984 | E018250 | right | TTAA | G2e3 | intron |
| chr6.1:475797 | chr6.1 | 475797 | 8984 | E018250 | right | TTAA | Zbtb46 | intron |
| chr6.1:475800 | chr6.1 | 475800 | 1 | E018250 | left | TTAA | Zbtb46 | intron |
| chr6.1:144934972 | chr6.1 | 144934972 | 1 | E018250 | left | TTAA | Ralgps1 | intron |
| chr6.1:144934975 | chr6.1 | 144934975 | 8984 | E018250 | right | TTAA | Ralgps1 | intron |
| chr8.1:29566646 | chr8.1 | 29566646 | 8984 | E018250 | right | ATCTGCA | Syn2 | intron |
| chr8.1:29566656 | chr8.1 | 29566656 | 1 | E018250 | left | TTAA | Syn2 | intron |
| chr8.1:58303039 | chr8.1 | 58303039 | 1 | E018250 | left | AAAGTTA | NA | NA |
| chr8.1:58303043 | chr8.1 | 58303043 | 8985 | E018250 | right | TTAA | NA | NA |
| chrX.1:115648620 | chrX.1 | 115648620 | 1 | E018250 | left | TTAA | NA | NA |
| chrX.1:115648623 | chrX.1 | 115648623 | 8984 | E018250 | right | TTAA | NA | NA |
| chr1-0.1:64773575 | chr1-0.1 | 64773575 | 8984 | E019257 | right | TTAA | NA | NA |
| chr1-0.1:64773578 | chr1-0.1 | 64773578 | 1 | E019257 | left | TTAA | NA | NA |
| chr1-0.1:95366125 | chr1-0.1 | 95366125 | 8984 | E019257 | right | TTAA | Nbea | intron |
| chr1-0.1:95366128 | chr1-0.1 | 95366128 | 1 | E019257 | left | TTAA | Nbea | intron |
| chr1-0.1:177405375 | chr1-0.1 | 177405375 | 1 | E019257 | left | TTAA | NA | NA |
| chr1-0.1:177405378 | chr1-0.1 | 177405378 | 8984 | E019257 | right | TTAA | NA | NA |
| chr1-0.1:232966599 | chr1-0.1 | 232966599 | 1 | E019257 | left | TTAA | NA | NA |
| chr1-0.1:232966602 | chr1-0.1 | 232966602 | 8984 | E019257 | right | TTAA | NA | NA |
| chr1-1.1:19111579 | chr1-1.1 | 19111579 | 8984 | E019257 | right | TTAA | NA | NA |
| chr1-1.1:19111582 | chr1-1.1 | 19111582 | 1 | E019257 | left | TTAA | NA | NA |
| chr1-1.1:67490208 | chr1-1.1 | 67490208 | 1 | E019257 | left | TTAA | Tmco3 | intron |
| chr1-1.1:67490211 | chr1-1.1 | 67490211 | 8984 | E019257 | right | TTAA | Tmco3 | intron |
| chr1-1.1:107103242 | chr1-1.1 | 107103242 | 8984 | E019257 | right | TTAA | Nsg1 | intron |
| chr1-1.1:107103245 | chr1-1.1 | 107103245 | 1 | E019257 | left | TTAA | Nsg1 | intron |
| chr2.1:105329254 | chr2.1 | 105329254 | 8984 | E019257 | right | TTAA | NA | NA |
| chr2.1:105329257 | chr2.1 | 105329257 | 1 | E019257 | left | TTAA | NA | NA |
| chr2.1:123350723 | chr2.1 | 123350723 | 1 | E019257 | left | TTAA | NA | NA |
| chr2.1:123350726 | chr2.1 | 123350726 | 8984 | E019257 | right | TTAA | NA | NA |
| chr2.1:395107029 | chr2.1 | 395107029 | 1 | E019257 | left | TTAA | NA | NA |
| chr2.1:395107032 | chr2.1 | 395107032 | 8984 | E019257 | right | TTAA | NA | NA |
| chr3.1:70337525 | chr3.1 | 70337525 | 8984 | E019257 | right | TTAA | Ate1 | intron |
| chr4.1:152904949 | chr4.1 | 152904949 | 8980 | E019257 | right | ATACTAA | NA | NA |
| chr4.1:152904952 | chr4.1 | 152904952 | 6 | E019257 | left | CTAAGGA | NA | NA |
| chr6.1:78868878 | chr6.1 | 78868878 | 1 | E019257 | left | GTTGTTA | NA | NA |
| chr6.1:78868882 | chr6.1 | 78868882 | 8985 | E019257 | right | TTAA | NA | NA |
| chr9.1:25501741 | chr9.1 | 25501741 | 8984 | E019257 | right | TTAA | NA | NA |
| chr9.1:25501744 | chr9.1 | 25501744 | 1 | E019257 | left | TTAA | NA | NA |
| chr1-0.1:61277560 | chr1-0.1 | 61277560 | 1 | E019260 | left | TTAA | Mllt11 | intron |
| chr1-0.1:61277563 | chr1-0.1 | 61277563 | 8984 | E019260 | right | TTAA | Mllt11 | intron |
| chr10.1:28873990 | chr10.1 | 28873990 | 8984 | E019260 | right | TTAA | NA | NA |
| chr10.1:28874023 | chr10.1 | 28874023 | 1729 | E019260 | unknown | GTGTGAC | NA | NA |
| chr2.1:93652550 | chr2.1 | 93652550 | 5 | E019260 | left | AGAATAA | Grin3a | intron |
| chr2.1:93652553 | chr2.1 | 93652553 | 8983 | E019260 | right | ATAAACA | Grin3a | intron |
| chr2.1:395887666 | chr2.1 | 395887666 | 1 | E019260 | left | TTAA | NA | NA |
| chr2.1:395887669 | chr2.1 | 395887669 | 8984 | E019260 | right | TTAA | NA | NA |
| chrX.1:64673227 | chrX.1 | 64673227 | 1 | E019260 | left | TTAA | Atp7a | intron |
| chrX.1:64673230 | chrX.1 | 64673230 | 8984 | E019260 | right | TTAA | Atp7a | intron |
| chr1-0.1:91469064 | chr1-0.1 | 91469064 | 1 | E019268 | left | TTAA | NA | NA |

| | | | | | | | | |
|---|---|---|---|---|---|---|---|---|
| chr1-0.1:91469067 | chr1-0.1 | 91469067 | 8984 | E019268 | right | TTAA | NA | NA |
| chr1-0.1:239510004 | chr1-0.1 | 239510004 | 8984 | E019268 | right | TTAA | LOC103160224 | intron |
| chr1-1.1:155458251 | chr1-1.1 | 155458251 | 1 | E019268 | left | TTAA | NA | NA |
| chr2.1:279506002 | chr2.1 | 279506002 | 1 | E019268 | left | TTAA | NA | CDS |
| chr2.1:279506005 | chr2.1 | 279506005 | 8984 | E019268 | right | TTAA | NA | CDS |
| chr4.1:167970822 | chr4.1 | 167970822 | 1 | E019268 | left | TTAA | Hmg20a | intron |
| chr4.1:167970825 | chr4.1 | 167970825 | 8984 | E019268 | right | TTAA | Hmg20a | intron |
| chr5.1:38435251 | chr5.1 | 38435251 | 8984 | E019268 | right | TTAA | NA | NA |
| chr5.1:38435254 | chr5.1 | 38435254 | 1 | E019268 | left | TTAA | NA | NA |
| chr5.1:52829772 | chr5.1 | 52829772 | 5 | E019268 | left | TCAGTAT | NA | CDS |
| chr5.1:52829775 | chr5.1 | 52829775 | 8980 | E019268 | right | GTATGTC | NA | CDS |
| chr7.1:86922265 | chr7.1 | 86922265 | 8984 | E019268 | right | TTAA | NA | NA |
| chr7.1:86922268 | chr7.1 | 86922268 | 1 | E019268 | left | TTAA | NA | NA |
| chr7.1:114835423 | chr7.1 | 114835423 | 1 | E019268 | left | TTAA | LOC103164366 | intron |
| chr7.1:114835426 | chr7.1 | 114835426 | 8984 | E019268 | right | TTAA | LOC103164366 | intron |
| chr1-0.1:175172933 | chr1-0.1 | 175172933 | 1 | E019275 | left | TTAA | Magi2 | intron |
| chr1-0.1:175172936 | chr1-0.1 | 175172936 | 8984 | E019275 | right | TTAA | Magi2 | intron |
| chr1-1.1:57803505 | chr1-1.1 | 57803505 | 8984 | E019275 | right | TTAA | Fnta | intron |
| chr1-1.1:57803508 | chr1-1.1 | 57803508 | 1 | E019275 | left | TTAA | Fnta | intron |
| chr2.1:425378276 | chr2.1 | 425378276 | 8984 | E019275 | right | TTAA | NA | NA |
| chr2.1:425378279 | chr2.1 | 425378279 | 1 | E019275 | left | TTAA | NA | NA |
| chr3.1:93667465 | chr3.1 | 93667465 | 1 | E019275 | left | TTAA | NA | NA |
| chr3.1:93667467 | chr3.1 | 93667467 | 8984 | E019275 | right | TTAA | NA | NA |
| chr4.1:174329154 | chr4.1 | 174329154 | 1 | E019275 | left | TTAA | Cln6 | intron |
| chr8.1:59005310 | chr8.1 | 59005310 | 8984 | E019275 | right | TTAA | NA | NA |
| chr8.1:59005313 | chr8.1 | 59005313 | 1 | E019275 | left | TTAA | NA | NA |
| chrX.1:40638716 | chrX.1 | 40638716 | 1 | E019275 | left | TTAA | Tbc1d8b | intron |
| chrX.1:40638719 | chrX.1 | 40638719 | 8984 | E019275 | right | TTAA | Tbc1d8b | intron |
| chrX.1:87493780 | chrX.1 | 87493780 | 8984 | E019275 | right | TTAA | NA | NA |
| chrX.1:87493783 | chrX.1 | 87493783 | 1 | E019275 | left | TTAA | NA | NA |
| chr3.1:80339249 | chr3.1 | 80339249 | 8984 | E020278 | right | TTAA | NA | CDS |
| chr3.1:80339252 | chr3.1 | 80339252 | 8984 | E020278 | right | TTAA | NA | CDS |
| chr3.1:101376185 | chr3.1 | 101376185 | 1 | E020278 | left | GCAGTTA | NA | NA |
| chr3.1:243224128 | chr3.1 | 243224128 | 1 | E020278 | left | TTAA | Aoah | intron |
| chr4.1:208826585 | chr4.1 | 208826585 | 8984 | E020278 | right | TTAA | NA | NA |
| chr4.1:208826588 | chr4.1 | 208826588 | 1 | E020278 | left | TTAA | NA | NA |
| chr5.1:27325093 | chr5.1 | 27325093 | 1 | E020278 | left | TTAA | NA | NA |
| chr5.1:27325096 | chr5.1 | 27325096 | 8984 | E020278 | right | TTAA | NA | NA |
| chr6.1:128457637 | chr6.1 | 128457637 | 1 | E020278 | left | TTAA | NA | NA |
| chr6.1:128457640 | chr6.1 | 128457640 | 8984 | E020278 | right | TTAA | NA | NA |
| chr8.1:18110072 | chr8.1 | 18110072 | 1 | E020278 | left | TTAA | LOC118239313 | intron |
| chrX.1:51333927 | chrX.1 | 51333927 | 1 | E020278 | left | TTAA | NA | NA |
| chrX.1:51333930 | chrX.1 | 51333930 | 8984 | E020278 | right | TTAA | NA | NA |
| chr1-0.1:52612709 | chr1-0.1 | 52612709 | 8984 | E020284 | right | TTAA | NA | NA |
| chr1-0.1:52612712 | chr1-0.1 | 52612712 | 1 | E020284 | left | TTAA | NA | NA |
| chr1-0.1:64944485 | chr1-0.1 | 64944485 | 1 | E020284 | left | TTAA | NA | NA |
| chr1-0.1:64944488 | chr1-0.1 | 64944488 | 8984 | E020284 | right | TTAA | NA | NA |
| chr1-0.1:75023596 | chr1-0.1 | 75023596 | 8984 | E020284 | right | TTAA | Rapgef2 | intron |
| chr1-0.1:75023599 | chr1-0.1 | 75023599 | 1 | E020284 | left | TTAA | Rapgef2 | intron |
| chr1-0.1:95555040 | chr1-0.1 | 95555040 | 1 | E020284 | left | TTAA | NA | NA |
| chr1-0.1:95555043 | chr1-0.1 | 95555043 | 8984 | E020284 | right | TTAA | NA | NA |
| chr1-0.1:140422263 | chr1-0.1 | 140422263 | 1 | E020284 | left | TTAA | NA | NA |
| chr1-0.1:140422266 | chr1-0.1 | 140422266 | 8984 | E020284 | right | TTAA | NA | NA |
| chr1-0.1:150922679 | chr1-0.1 | 150922679 | 8984 | E020284 | right | TTAA | NA | NA |
| chr1-0.1:150922682 | chr1-0.1 | 150922682 | 1 | E020284 | left | TTAA | NA | NA |
| chr1-0.1:186928402 | chr1-0.1 | 186928402 | 8984 | E020284 | right | TTAA | Adam22 | intron |
| chr1-0.1:186928405 | chr1-0.1 | 186928405 | 1 | E020284 | left | TTAA | Adam22 | intron |
| chr1-0.1:229497783 | chr1-0.1 | 229497783 | 8984 | E020284 | right | TTAA | NA | CDS |
| chr1-0.1:229497786 | chr1-0.1 | 229497786 | 1 | E020284 | left | TTAA | NA | CDS |
| chr1-0.1:256479600 | chr1-0.1 | 256479600 | 1 | E020284 | left | AAAGTTA | LOC103160738 | intron |
| chr1-0.1:256479604 | chr1-0.1 | 256479604 | 8985 | E020284 | right | TTAA | LOC103160738 | intron |
| chr1-1.1:69691575 | chr1-1.1 | 69691575 | 5798 | E020284 | unknown | GTCTTTA | NA | NA |
| chr1-1.1:69691576 | chr1-1.1 | 69691576 | 1 | E020284 | left | TTAA | NA | NA |
| chr1-1.1:103663864 | chr1-1.1 | 103663864 | 1 | E020284 | left | TTAA | Grk4 | intron |
| chr1-1.1:103663867 | chr1-1.1 | 103663867 | 8984 | E020284 | right | TTAA | Grk4 | intron |
| chr1-1.1:121283195 | chr1-1.1 | 121283195 | 8985 | E020284 | right | TTAA | NA | NA |
| chr1-1.1:121283199 | chr1-1.1 | 121283199 | 1 | E020284 | left | TAACTTT | NA | NA |
| chr1-1.1:147576771 | chr1-1.1 | 147576771 | 8985 | E020284 | right | TTAA | Adgrl3 | intron |
| chr1-1.1:147576775 | chr1-1.1 | 147576775 | 1 | E020284 | left | TAACTCT | Adgrl3 | intron |
| chr1-1.1:203250335 | chr1-1.1 | 203250335 | 1 | E020284 | left | TTAA | NA | NA |
| chr1-1.1:203250339 | chr1-1.1 | 203250339 | 8984 | E020284 | right | TAACTGG | NA | NA |
| chr1-1.1:219694033 | chr1-1.1 | 219694033 | 1 | E020284 | left | TTAA | Kif13b | intron |
| chr1-1.1:219694036 | chr1-1.1 | 219694036 | 8984 | E020284 | right | TTAA | Kif13b | intron |
| chr1-1.1:231094832 | chr1-1.1 | 231094832 | 8984 | E020284 | right | TTAA | LOC113832677 | intron |
| chr1-1.1:231094835 | chr1-1.1 | 231094835 | 1 | E020284 | left | TTAA | LOC113832677 | intron |
| chr1-1.1:270459267 | chr1-1.1 | 270459267 | 1 | E020284 | left | TTAA | NA | exon |
| chr1-1.1:270459270 | chr1-1.1 | 270459270 | 8984 | E020284 | right | TTAA | NA | exon |
| chr2.1:22920319 | chr2.1 | 22920319 | 1 | E020284 | left | TTAA | Pef1 | intron |
| chr2.1:22920322 | chr2.1 | 22920322 | 8984 | E020284 | right | TTAA | Pef1 | intron |

| | | | | | | | | |
|---|---|---|---|---|---|---|---|---|
| chr2.1:29977123 | chr2.1 | 29977123 | 8984 | E020284 | right | TTAA | Trit1 | intron |
| chr2.1:29977126 | chr2.1 | 29977126 | 1 | E020284 | left | TTAA | Trit1 | intron |
| chr2.1:32037533 | chr2.1 | 32037533 | 8984 | E020284 | right | TTAA | LOC118238296 | intron |
| chr2.1:32037536 | chr2.1 | 32037536 | 1 | E020284 | left | TTAA | LOC118238296 | intron |
| chr2.1:34342299 | chr2.1 | 34342299 | 8984 | E020284 | right | TTAA | St3gal3 | intron |
| chr2.1:34342302 | chr2.1 | 34342302 | 1 | E020284 | left | TTAA | St3gal3 | intron |
| chr2.1:44704464 | chr2.1 | 44704464 | 8984 | E020284 | right | TTAA | NA | NA |
| chr2.1:44704467 | chr2.1 | 44704467 | 1 | E020284 | left | TTAA | NA | NA |
| chr2.1:60668568 | chr2.1 | 60668568 | 8984 | E020284 | right | TTAA | NA | NA |
| chr2.1:60668571 | chr2.1 | 60668571 | 1 | E020284 | left | TTAA | NA | NA |
| chr2.1:162889090 | chr2.1 | 162889090 | 8984 | E020284 | right | TTAA | NA | NA |
| chr2.1:162889093 | chr2.1 | 162889093 | 1 | E020284 | left | TTAA | NA | NA |
| chr2.1:204592040 | chr2.1 | 204592040 | 1 | E020284 | left | TTAA | NA | NA |
| chr2.1:204592043 | chr2.1 | 204592043 | 8984 | E020284 | right | TTAA | NA | NA |
| chr2.1:236322733 | chr2.1 | 236322733 | 8983 | E020284 | right | ACATTAG | NA | NA |
| chr2.1:236322736 | chr2.1 | 236322736 | 5 | E020284 | left | TTAGTAT | NA | NA |
| chr2.1:334262861 | chr2.1 | 334262861 | 8984 | E020284 | right | TTAA | NA | NA |
| chr2.1:334262864 | chr2.1 | 334262864 | 1 | E020284 | left | TTAA | NA | NA |
| chr2.1:459222184 | chr2.1 | 459222184 | 1 | E020284 | left | TTAA | LOC103161380 | intron |
| chr2.1:459222188 | chr2.1 | 459222188 | 8984 | E020284 | right | TAACTGC | LOC103161380 | intron |
| chr3.1:55005391 | chr3.1 | 55005391 | 8984 | E020284 | right | TTAA | LOC103160214 | intron |
| chr3.1:55005394 | chr3.1 | 55005394 | 1 | E020284 | left | TTAA | LOC103160214 | intron |
| chr3.1:58445499 | chr3.1 | 58445499 | 1 | E020284 | left | TTAA | LOC118238546 | intron |
| chr3.1:58445502 | chr3.1 | 58445502 | 8984 | E020284 | right | TTAA | LOC118238546 | intron |
| chr3.1:106380215 | chr3.1 | 106380215 | 1 | E020284 | left | TTAA | LOC103162587 | intron |
| chr3.1:106380218 | chr3.1 | 106380218 | 8984 | E020284 | right | TTAA | LOC103162587 | intron |
| chr3.1:109204836 | chr3.1 | 109204836 | 1 | E020284 | left | TTAA | NA | NA |
| chr3.1:109204839 | chr3.1 | 109204839 | 8984 | E020284 | right | TTAA | NA | NA |
| chr3.1:123682321 | chr3.1 | 123682321 | 8984 | E020284 | right | TTAA | LOC113834919 | intron |
| chr3.1:123682324 | chr3.1 | 123682324 | 1 | E020284 | left | TTAA | LOC113834919 | intron |
| chr3.1:129616051 | chr3.1 | 129616051 | 8984 | E020284 | right | TTAA | Lrrc28 | intron |
| chr3.1:129616054 | chr3.1 | 129616054 | 1 | E020284 | left | TTAA | Lrrc28 | intron |
| chr3.1:148503561 | chr3.1 | 148503561 | 8984 | E020284 | right | TTAA | Slc17a6 | intron |
| chr3.1:148503564 | chr3.1 | 148503564 | 1 | E020284 | left | TTAA | Slc17a6 | intron |
| chr3.1:158114264 | chr3.1 | 158114264 | 1 | E020284 | left | TTAA | Il15 | intron |
| chr3.1:158114267 | chr3.1 | 158114267 | 8984 | E020284 | right | TTAA | Il15 | intron |
| chr3.1:187238980 | chr3.1 | 187238980 | 1 | E020284 | left | TTAA | Sf3b3 | intron |
| chr3.1:187238983 | chr3.1 | 187238983 | 8984 | E020284 | right | TTAA | Sf3b3 | intron |
| chr3.1:202717918 | chr3.1 | 202717918 | 8984 | E020284 | right | TAACAGC | NA | exon |
| chr4.1:46242188 | chr4.1 | 46242188 | 1 | E020284 | left | TTAA | Usf3 | intron |
| chr4.1:46242191 | chr4.1 | 46242191 | 8984 | E020284 | right | TTAA | Usf3 | intron |
| chr4.1:192862096 | chr4.1 | 192862096 | 8984 | E020284 | right | TTAA | NA | NA |
| chr4.1:192862099 | chr4.1 | 192862099 | 1 | E020284 | left | TTAA | NA | NA |
| chr4.1:214377883 | chr4.1 | 214377883 | 1 | E020284 | left | TTAA | Wdr82 | intron |
| chr4.1:214377886 | chr4.1 | 214377886 | 8984 | E020284 | right | TTAA | Wdr82 | intron |
| chr5.1:68922646 | chr5.1 | 68922646 | 8984 | E020284 | right | TTAA | Epb41l5 | intron |
| chr5.1:68922649 | chr5.1 | 68922649 | 1 | E020284 | left | TTAA | Epb41l5 | intron |
| chr5.1:170749294 | chr5.1 | 170749294 | 8984 | E020284 | right | TTAA | NA | NA |
| chr5.1:170749297 | chr5.1 | 170749297 | 1 | E020284 | left | TTAA | NA | NA |
| chr6.1:59740900 | chr6.1 | 59740900 | 1 | E020284 | left | TTAA | NA | NA |
| chr6.1:59740903 | chr6.1 | 59740903 | 8984 | E020284 | right | TTAA | NA | NA |
| chr6.1:107890556 | chr6.1 | 107890556 | 8983 | E020284 | right | TACTAGA | NA | NA |
| chr6.1:107890558 | chr6.1 | 107890558 | 5 | E020284 | left | CTAGAAT | NA | NA |
| chr7.1:8936602 | chr7.1 | 8936602 | 8985 | E020284 | right | TTAA | Ttc27 | intron |
| chr7.1:8936606 | chr7.1 | 8936606 | 1 | E020284 | left | TAACTGG | Ttc27 | intron |
| chr7.1:47070730 | chr7.1 | 47070730 | 1 | E020284 | left | TTAA | Crebrf | intron |
| chr7.1:47070733 | chr7.1 | 47070733 | 8984 | E020284 | right | TTAA | Crebrf | intron |
| chr7.1:110317615 | chr7.1 | 110317615 | 8984 | E020284 | right | TTAA | Kpnb1 | intron |
| chr7.1:110317616 | chr7.1 | 110317616 | 1 | E020284 | left | TTAA | Kpnb1 | intron |
| chr1-0.1:49516251 | chr1-0.1 | 49516251 | 1 | E020285 | left | AAAGGTT | NA | NA |
| chr1-0.1:49516256 | chr1-0.1 | 49516256 | 8985 | E020285 | right | TTAA | NA | NA |
| chr1-0.1:177181929 | chr1-0.1 | 177181929 | 8984 | E020285 | right | TTAA | LOC103162716 | intron |
| chr1-0.1:177181932 | chr1-0.1 | 177181932 | 1 | E020285 | left | TTAA | LOC103162716 | intron |
| chr1-0.1:212443708 | chr1-0.1 | 212443708 | 1 | E020285 | left | TTAA | Bmt2 | intron |
| chr1-0.1:212443711 | chr1-0.1 | 212443711 | 8984 | E020285 | right | TTAA | Bmt2 | intron |
| chr2.1:36264531 | chr2.1 | 36264531 | 1 | E020285 | left | TTAA | Lrrc41 | intron |
| chr2.1:36264535 | chr2.1 | 36264535 | 8984 | E020285 | right | TAACTCT | Lrrc41 | intron |
| chr2.1:191649892 | chr2.1 | 191649892 | 8984 | E020285 | right | TTAA | NA | NA |
| chr2.1:191649895 | chr2.1 | 191649895 | 1 | E020285 | left | TTAA | NA | NA |
| chr2.1:370331063 | chr2.1 | 370331063 | 1 | E020285 | left | TTAA | Trappc9 | intron |
| chr2.1:370331067 | chr2.1 | 370331067 | 8984 | E020285 | right | TAACAGG | Trappc9 | intron |
| chr3.1:251007116 | chr3.1 | 251007116 | 1 | E020285 | left | TTAA | NA | NA |
| chr5.1:69124923 | chr5.1 | 69124923 | 1 | E020285 | left | TTAA | NA | NA |
| chr5.1:69124926 | chr5.1 | 69124926 | 8984 | E020285 | right | TTAA | NA | NA |
| chr5.1:166158331 | chr5.1 | 166158331 | 1 | E020285 | left | TTAA | Ahr | intron |
| chr5.1:166158334 | chr5.1 | 166158334 | 8984 | E020285 | right | TTAA | Ahr | intron |
| chr6.1:47900699 | chr6.1 | 47900699 | 1 | E020285 | left | AGAGGTT | Itpa | intron |
| chr6.1:47900704 | chr6.1 | 47900704 | 8985 | E020285 | right | TTAA | Itpa | intron |
| chr6.1:76240448 | chr6.1 | 76240448 | 10781 | E020285 | unknown | TTAA | NA | NA |

| | | | | | | | | |
|---|---|---|---|---|---|---|---|---|
| chr6.1:76240452 | chr6.1 | 76240452 | 1 | E020285 | left | TAACTCT | NA | NA |
| chr6.1:137828327 | chr6.1 | 137828327 | 1 | E020285 | left | TTAA | Sec16a | intron |
| chr6.1:137828330 | chr6.1 | 137828330 | 1539 | E020285 | unknown | TTAA | Sec16a | intron |
| chr6.1:144240687 | chr6.1 | 144240687 | 1 | E020285 | left | TTAA | NA | NA |
| chr6.1:144240690 | chr6.1 | 144240690 | 8984 | E020285 | right | TTAA | NA | NA |
| chr7.1:35418620 | chr7.1 | 35418620 | 1 | E020285 | left | TTAA | NA | NA |
| chr7.1:35418623 | chr7.1 | 35418623 | 8984 | E020285 | right | TTAA | NA | NA |
| chr7.1:67226405 | chr7.1 | 67226405 | 1 | E020285 | left | AAAGTTA | Fstl4 | intron |
| chr7.1:67226409 | chr7.1 | 67226409 | 8985 | E020285 | right | TTAA | Fstl4 | intron |
| chr8.1:46103716 | chr8.1 | 46103716 | 8984 | E020285 | right | TTAA | Mitf | intron |
| chr8.1:46103719 | chr8.1 | 46103719 | 1 | E020285 | left | TTAA | Mitf | intron |
| chr8.1:64716237 | chr8.1 | 64716237 | 8984 | E020285 | right | TTAA | NA | NA |
| chr8.1:64716240 | chr8.1 | 64716240 | 1 | E020285 | left | TTAA | NA | NA |
| chr8.1:69538945 | chr8.1 | 69538945 | 1 | E020285 | left | TTAA | Ctnna2 | intron |
| chr8.1:69538948 | chr8.1 | 69538948 | 8984 | E020285 | right | TTAA | Ctnna2 | intron |
| chr1-0.1:174469334 | chr1-0.1 | 174469334 | 8984 | E020293 | right | GCTGTTA | Magi2 | intron |
| chr1-0.1:174469338 | chr1-0.1 | 174469338 | 1 | E020293 | left | TTAA | Magi2 | intron |
| chr1-1.1:247083874 | chr1-1.1 | 247083874 | 2 | E020293 | left | TGATTAT | NA | NA |
| chr1-1.1:247083876 | chr1-1.1 | 247083876 | 8984 | E020293 | right | ATTATGA | NA | NA |
| chr2.1:397470 | chr2.1 | 397470 | 8984 | E020293 | right | TTAA | Pusl1 | intron |
| chr2.1:397473 | chr2.1 | 397473 | 1 | E020293 | left | TTAA | Pusl1 | intron |
| chr2.1:5553476 | chr2.1 | 5553476 | 8984 | E020293 | right | TTAA | NA | NA |
| chr2.1:5553479 | chr2.1 | 5553479 | 1 | E020293 | left | TTAA | NA | NA |
| chr3.1:10577161 | chr3.1 | 10577161 | 8984 | E020293 | right | TTAA | NA | NA |
| chr4.1:215523312 | chr4.1 | 215523312 | 8984 | E020293 | right | TTAA | CUNH3orf18 | intron |
| chr5.1:29303078 | chr5.1 | 29303078 | 8984 | E020293 | right | TTAA | Fmo1 | intron |
| chr5.1:29303081 | chr5.1 | 29303081 | 1 | E020293 | left | TTAA | Fmo1 | intron |
| chr5.1:80113116 | chr5.1 | 80113116 | 1 | E020293 | left | TTAA | NA | NA |
| chr5.1:80113119 | chr5.1 | 80113119 | 8984 | E020293 | right | TTAA | NA | NA |
| chr8.1:11486536 | chr8.1 | 11486536 | 1 | E020293 | left | TTAA | Eps8 | intron |
| chrX.1:67832296 | chrX.1 | 67832296 | 1 | E020293 | left | TTAA | Hdac8 | intron |
| chrX.1:67832299 | chrX.1 | 67832299 | 8984 | E020293 | right | TTAA | Hdac8 | intron |

| IS_location | count |
|---|---|
| CDS | 8 |
| Exon | 11.5 |
| Intron | 312 |
| NA | 316 |

| type | count |
| --- | --- |
| left | 632 |
| right | 631 |
| unknown | 32 |

| clone no. | integration_sites | HC | LC | Mean Copies |
|---|---|---|---|---|
| E003102 | 53.50 | 33.72 | 31.23 | 32.48 |
| E013146 | 8.50 | 7.91 | 9.09 | 8.50 |
| E014156 | 20.00 | 18.40 | 17.36 | 17.88 |
| E014157 | 7.50 | 10.55 | 10.98 | 10.77 |
| E014158 | 13.50 | 16.54 | 21.87 | 19.20 |
| E014160 | 12.00 | 69.09 | 63.92 | 66.51 |
| E014163 | 5.50 | 7.53 | 7.98 | 7.75 |
| E014168 | 26.50 | 22.71 | 26.68 | 24.69 |
| E016186 | 13.50 | 12.03 | 19.20 | 15.61 |
| E016188 | 16.00 | 18.73 | 19.79 | 19.26 |
| E016191 | 21.50 | 22.03 | 19.04 | 20.53 |
| E016192 | 18.00 | 24.22 | 19.81 | 22.01 |
| E006206 | 9.00 | 6.29 | 7.72 | 7.00 |
| E007215 | 10.00 | 9.92 | 8.16 | 9.04 |
| E017223 | 8.50 | 13.00 | 11.90 | 12.45 |
| E017227 | 6.00 | 8.60 | 14.84 | 11.72 |
| E017233 | 13.00 | 11.86 | 19.53 | 15.70 |
| E018244 | 12.00 | 11.40 | 10.98 | 11.19 |
| E018247 | 19.00 | 11.92 | 11.84 | 11.88 |
| E018249 | 10.50 | 10.92 | 11.41 | 11.17 |
| E018250 | 16.00 | 14.30 | 14.89 | 14.59 |
| E020278 | 6.50 | 6.32 | 8.41 | 7.37 |
| E020284 | 47.50 | 53.51 | 57.94 | 55.72 |
| E020285 | 17.50 | 15.83 | 18.08 | 16.96 |
| E003104 | 8.50 | 8.56 | 8.29 | 8.42 |
| E003115 | 7.50 | 8.77 | 8.81 | 8.79 |
| E004130 | 9.00 | 9.58 | 9.78 | 9.68 |
| E004134 | 5.00 | 5.61 | 5.84 | 5.72 |
| E013141 | 6.00 | 6.38 | 6.48 | 6.43 |
| E013145 | 10.00 | 9.40 | 9.84 | 9.62 |
| E014154 | 4.50 | 7.65 | 8.43 | 8.04 |
| E014159 | 22.50 | 11.51 | 11.50 | 11.50 |
| E014162 | 8.50 | 10.05 | 10.35 | 10.20 |
| E014167 | 8.50 | 8.38 | 10.02 | 9.20 |
| E015170 | 7.50 | 7.40 | 7.88 | 7.64 |
| E016189 | 19.50 | 34.08 | 32.68 | 33.38 |
| E006200 | 11.00 | 10.63 | 11.89 | 11.26 |
| E006204 | 6.00 | 7.23 | 8.11 | 7.67 |
| E017228 | 42.50 | 23.43 | 23.52 | 23.47 |
| E017231 | 6.50 | 8.76 | 7.81 | 8.28 |
| E017242 | 11.50 | 18.12 | 19.14 | 18.63 |
| E018245 | 9.50 | 8.46 | 8.90 | 8.68 |
| E018246 | 9.50 | 10.53 | 10.63 | 10.58 |
| E019257 | 13.50 | 14.06 | 13.54 | 13.80 |
| E019260 | 5.00 | 13.21 | 14.13 | 13.67 |
| E019268 | 8.00 | 8.63 | 7.60 | 8.12 |

| | | | | |
|---|---|---|---|---|
| E019275 | 7.50 | 8.66 | 8.63 | 8.65 |
| E020293 | 8.50 | 8.42 | 8.35 | 8.38 |
